# Benchmarking single cell transcriptome matching methods for incremental growth of cell atlases

**DOI:** 10.1101/2025.04.10.648034

**Authors:** Joyce Hu, Beverly Peng, Ajith V. Pankajam, Bingfang Xu, Vikrant Anil Deshpande, Andreas Bueckle, Bruce W. Herr, Christopher L. Dupont, Katy Börner, Richard H. Scheuermann, Yun Zhang

**Author notes:** These authors contributed equally to this manuscript (Joyce Hu, Beverly Peng, Yun Zhang). Corresponding author: Yun Zhang.

## Abstract

**Background:** The advancement of single cell technologies has driven significant progress in constructing a multiscale, pan-organ Human Reference Atlas for healthy human cells. Many multi-faceted cell atlases for different organs, species, and diseases now exist, though challenges remain in harmonizing cell types and unifying nomenclature among respective cell atlases. Multiple machine learning and artificial intelligence methods, including models pre-trained on large-scale cell atlas datasets, are publicly available for single cell community users to computationally map their cell clusters to the cell atlases.

**Results:** This study benchmarks seven computational tools for cell type matching and label transfer – Azimuth, CellTypist, CellHint, FR-Match, scArches, scPred, and singleR – in ten organ systems. Using healthy lung as an exemplary organ, when matching the well-annotated cell types in two atlases – the Human Lung Cell Atlas (HLCA) and the LungMAP Single-Cell Reference (CellRef), variations in the matching accuracy were observed, especially in rare cell types, underlining the need for a consensus strategy using a selective set of computational methods. In the meta-analysis, the benchmarked methods were used to incrementally integrate 61 cell types from HLCA and 48 from CellRef, resulting in a cell meta-atlas of 41 matched, 20 HLCA-, and 7 CellRef-specific cell types. Similar approach revealed 25 matched cell types existed in two independent kidney atlases. Generalizability of the benchmarking performances were further demonstrated in a variety of organ systems.

**Conclusion:** This study reveals complementing strengths of the benchmarked methods and presents a framework for incremental growth of cell types in cell atlases.

## Introduction

The advancement of single cell transcriptomic technologies has led to the rapid growth of single cell/nucleus RNA-seq (sc/snRNA-seq) datasets. Several large international consortia have made these data publicly available to accelerate the cellular characterization of complex tissues. As of March 2026, the CZ CellxGene Discover platform [1] (https://cellxgene.cziscience.com) has a data corpus of 149 million unique cells from 2,076 datasets; the Human BioMolecular Atlas Program (HuBMAP) [2] data portal (https://portal.hubmapconsortium.org) contains 5,640 datasets for 31 organs; the Single Cell Portal [3] (https://singlecell.broadinstitute.org) features 972 studies of ∼74 million cells. These efforts have contributed to the ongoing endeavors to build comprehensive reference cell atlases for human tissues (e.g., the Human Cell Atlas (HCA) [4] and Human Reference Atlas (HRA) [5, 6]) at single cell resolution. One of the goals of these atlasing projects is to achieve a community-driven consensus of cell phenotypes (cell types and cell states) in the human body. While there is a growing consensus on gene names and anatomical structures, there is less agreement on the nomenclature of cell phenotypes and their defining characteristics, making the harmonization of cell types between datasets, technologies, and atlases challenging.

The review by Hemberg et al. highlights achievements and challenges in the field of single cell biology, including the need to benchmark the rapidly expanding set of computational tools and to conduct meaningful meta-analyses for biological insights [7]. Over the past few years, numerous computational methods have been developed for cell type integration based on batch correction and label transfer approaches. Batch correction methods, such as Harmony [8], BBKNN [9], and scVI [10], aim to eliminate experimental batch effects and technical noises while preserving true biological variation. Label transfer methods, such as Seurat [11], LIGER [12], and scANVI [13] aim to jointly model heterogeneous cells across datasets, modalities, and conditions to form a shared latent space for nearest neighbor search. Luecken et al. [14] benchmarked 16 batch correction and label transfer methods and revealed complementing properties of these methods. Combining the strengths of these methods, the first versions of cell atlases were constructed by integrating data from selected high-quality studies and refining cell type annotation with expert curation. For example, scVI and scANVI-based methods, suggested by Luecken et al., were optimized to build the integrated Human Lung Cell Atlas (HLCA) v1.0 [15] using 14 published and unpublished datasets by the computational and domain experts from the HCA Lung Network, where dataset-specific batch effects were removed and consensus cell type annotations were defined after correcting mislabeled and underlabeled cells.

Following a similar strategy, the Human CellCards Multi-Study CellRef 1.0 Atlas [16] was built by the LungMAP Consortium. Similarly for kidney, the Human Kidney Atlas 1.0 (HKA) [17] was built by cross-consortia efforts, including Kidney Precision Medicine Project (KPMP), HuBMAP, and HCA. The multimodal benchmarking dataset for renal cortex (mBDRC) [18] was generated as part of the CZI Seed Networks for HCA. These data-driven and well-curated cell atlases form the foundation for building a cell phenotype knowledgebase that covers all healthy cell types and generalizes to data from future disease-related studies.

So far, there is not a preferred strategy to update a cell atlas when new data become available or when merging two overlapping cell atlases. The current strategy, which requires data integration and joint clustering, is problematic for two reasons. First, every update will require a re-analysis of all cells, including the cells that are already in the atlas. Re-analysis will introduce slightly different groupings of cells, so that studies which used an old version of the atlas will not be reproducible with an updated version, contradicting the reproducibility desired for reference cell atlases. Second, the joint clustering and manual annotation procedure is not efficient for the growing size of integrated datasets, as those computationally derived cell clusters will need to be re-annotated using marker genes and domain knowledge, which require manual curation to identify rare and/or novel cell types in the rapidly growing datasets.

To support the reproducibility of reference cell atlases and to construct a sustainable cell type knowledgebase, we propose a strategy for “incremental” growth of the cell types across cell atlases. This approach compares cell types between the query and reference datasets, so common cell types are merged, and novel cell types are added to the cell type knowledgebase. This strategy requires computational methods to perform comparative pattern matching at the cell type/cluster level. As an alternative to the cell-level label transfer methods, cell type/cluster-level matching methods, such as FR-Match [19, 20] and CellHint [21], will complement and potentially improve the long-term atlas building.

In this study, we present a framework for constructing a meta-atlas of healthy human cell types using benchmarked computational methods for incremental cell type knowledge growth (**Figure 1**) using lung as an exemplary organ. Similar analysis was applied to two kidney atlases and eight additional community-contributed atlases to illustrate the generalizability of the framework. For the healthy adult human lung, we harmonize cell types identified in two highly-recognized lung atlases – the Human Lung Cell Atlas (HLCA) [16] and the LungMAP Single-Cell Reference (CellRef) [15]. Evaluating all available methods is not feasible due to time and resource constraints. Instead, seven widely used methods were selected to cover different categories of cell type matching and label transfer approaches (**Figure 1A-B**). These seven include: pre-trained methods – Azimuth [22], CellTypist [23], CellHint [21], and scArches [24], and standalone methods – scPred [25], singleR [26], and FR-Match [19, 20]. Based on the initial benchmarking results, five methods were then used to integrate the 61 cell types identified in the HLCA with the 48 cell types identified in the CellRef.

**Figure 1.**
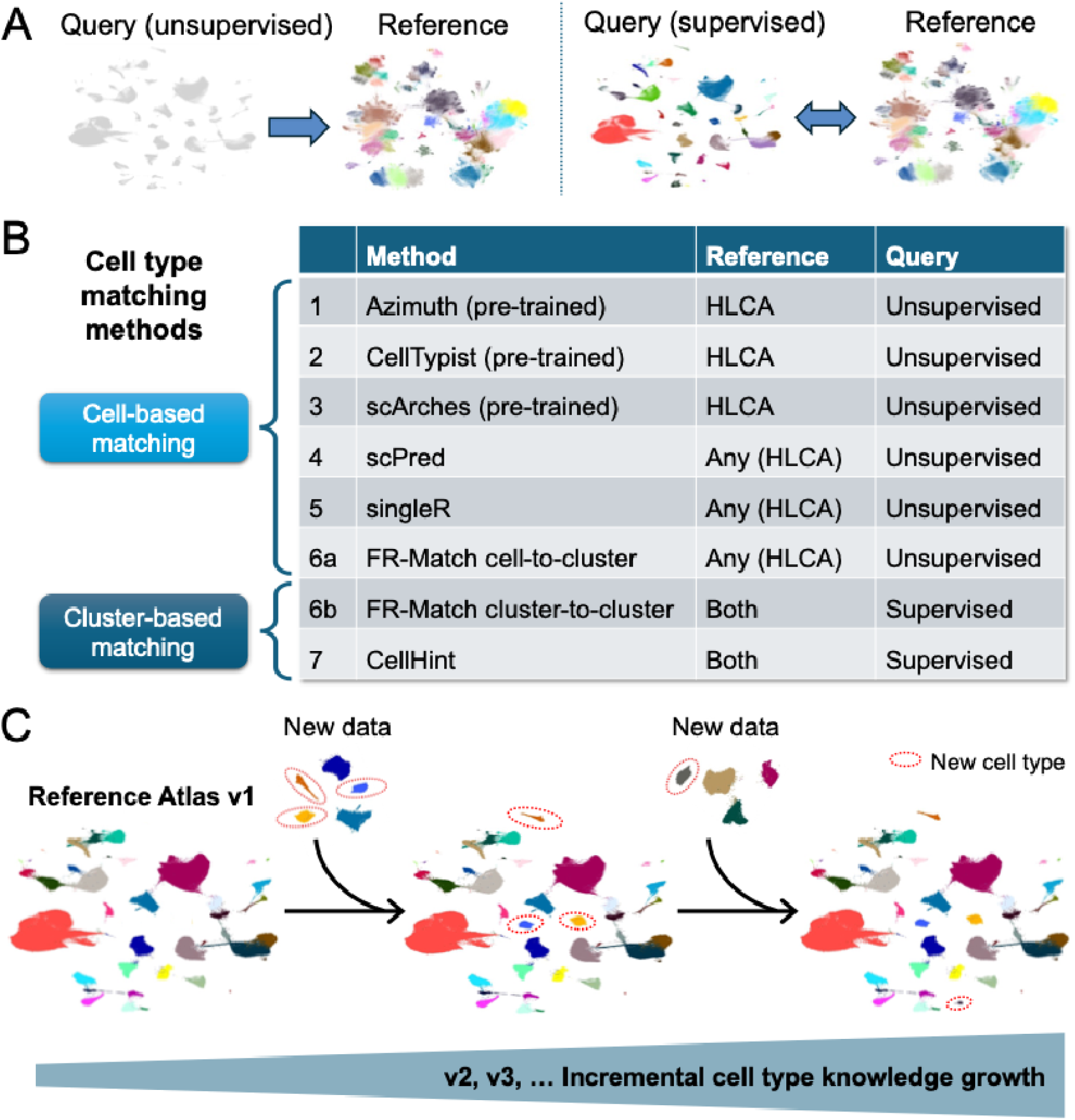
Study overview. Constructing an integrated human lung reference atlas using computational cell type matching between lung atlas datasets from single cell transcriptomic profiling, in support of incremental growth of a cell type knowledgebase. **(A)** Unsupervised and supervised cell type matching between query and reference datasets. The unsupervised cell type matching is when the query dataset has no cluster / cell type labels, so that the matching direction can only be in one direction from the query to the reference. The supervised cell type matching is when both datasets have annotated cell types, so that two-way cell type matching is possible to crossmatch the cell types in the two datasets. **(B)** Cell-based and cluster-based cell type matching methods. Azimuth, CellTypist, and scArches perform cell-based nearest neighbor label transfer, i.e., from query cell to reference cell, and assign the reference cell label to the query cell, using models pre-trained on a pre-defined reference dataset. The lung models in these three methods were pre-trained using the HLCA dataset as the reference. ScPred, singleR, and FR-Match are standalone methods for annotating scRNA-seq data using any reference data. For benchmarking, the same reference, HLCA, was used for these methods. FR-Match has both cell-based and cluster-based matching options: cell-to-cluster is from query cell to reference cluster and cluster-to-cluster is from query cluster to reference cluster. If both datasets are optimally-clustered, FR-Match cluster-to-cluster performs two-way matching (i.e., CellRef← → HLCA). CellHint performs cluster-level cell type harmonization with any datasets of choice. **(C)** A sustainable workflow for continued evolution of reference atlases requires a strategy that identifies new cell types from new datasets to be integrated to the reference atlas while preserving the cell type memberships of existing cells, to support the incremental growth of the knowledgebase.

To use Azimuth, CellTypist, and scArches, models pre-trained on the same reference dataset (i.e., HLCA) are publicly available. One potential shortcoming of these models is that they may not explicitly recognize novel cell types that are not in the reference during the pre-training process. To complement these pre-trained models, standalone methods were used for further evaluating the matching performance with potential candidates for dataset-specific cell types. All cell-based methods were used to match cell types from CellRef as the query dataset to HLCA as the reference dataset (i.e., CellRef → HLCA).

FR-Match has two matching options: cell-to-cluster and cluster-to-cluster. The cluster-based matching takes both datasets as query and reference, allowing for reciprocal two-way matching (HLCA <--> CellRef). The one-way (CellRef → HLCA) FR-Match cell-to-cluster matching was benchmarked with other cell-based methods. The two-way FR-Match cluster-to-cluster matching was benchmarked against CellHint – a CellTypist method that is designed for cluster-level cell type harmonization. To construct a meta-atlas of lung cell types, the cell-level matching results were summarized into cluster-level results, and the cell meta-atlas (i.e., matched cell types and dataset-specific cell types) was curated by examining all matching results and their data visualizations.

## Results

### Characteristics of methods and key datasets

In this study, we introduce the notions of unsupervised and supervised cell type matching and cell-based and cluster-based matching for describing the different matching methods. **Figure 1A** illustrates the unsupervised and supervised cell type matching scenarios. The setup of the cell type matching problem is to compare a query dataset to a reference dataset. The reference dataset is always an annotated dataset with clusters labeled by cell type names (visualized as colored dots on UMAP or tSNE plots). If the query dataset is unannotated (visualized as uncolored dots, but the grouping of cells can be obtained from the standard single cell analysis workflow), we refer the matching as unsupervised cell type matching. If the query dataset is also annotated, we refer the matching as supervised cell type matching. One advantage of the supervised matching is that matching direction can be two-way, i.e., query → reference and reference → query. Supervised matching is useful when matching cell types between two well-annotated cell atlases.

Key datasets used in this study are summarized in **Table 1**. Detailed description of key datasets is in **Methods**.

**Table 1.** Summary of healthy adult human lung and kidney atlases.

| Dataset | Version (Year) | Organ | Consortium | # cells/<br>nuclei | # genes | # cell<br>types | Platform | Sample<br>type |
| --- | --- | --- | --- | --- | --- | --- | --- | --- |
| HLCA<br>core | 1.0<br>(2023) | Lung | HCA | 584,944 | 28,024 | 61 | 10x 3' v1/v2/v3,<br>10x 5' v1/v2 | Cells &<br>nuclei |
| CellRef<br>human | 1.0<br>(2023) | Lung | LungMAP | 347,970 | 32,284 | 48 | 10x 3' v2/v3, 10x<br>5' v1 | Cells &<br>nuclei |
| HKA<br>normal | 1.0<br>(2023) | Kidney | KPMP,<br>HuBMAP, HCA | 107,701 | 33,826 | 75 | 10x 3' v3 | Cells &<br>nuclei |
| mBDRC | preprint<br>(2025) | Kidney | HCA | 97,125 | 36,972 | 36 | 10x 3' v3, 10x 5'<br>v1, 10x multiome | Cells &<br>nuclei |

We benchmark two major categories of methods: cell-based matching and cluster-based matching (**Figure 1B)**. One criterion used to select the cell-based methods is that the method must output a quantitative metric that reflects the matching confidence, i.e., “confidence score” (**Methods**). The confidence score provides the evidence for assigning a match (i.e., high confidence score = match, low confidence score = no match) and allows techniques such as ROC analysis to systematically evaluate the model performance across the spectrum of varying confidence threshold values. Based on this criterion, we selected the machine learning classification-based method, scPred [25], correlation-based method, singleR [26], the cell-to-cluster version of the graph-based method FR-Match [17] and pre-trained models: Azimuth [22], CellTypist [23], and scArches [24]. Azimuth and CellTypist provide multiple organ-specific models; the corresponding lung models pre-trained using the HLCA reference were used for benchmarking. ScArches is a pre-trained scvi-tools-based model that was used to build the joint embedding of the HLCA reference by the original authors. The pre-trained part of these models are the latent space embeddings and pre-tuned algorithms trained using the chosen reference, so that the input query data will be projected to the pre-trained latent space and annotated to the reference cell types by the pre-tuned algorithm. Despite the different underlying algorithms, these methods all perform cell-based matching (i.e., assigning each query cell to a reference cell type). The pre-trained models require minimal user input as no tuning of the algorithm is needed and no clustering is needed to prepare the query data. The downsides of pre-trained models are that the results are constrained to the reference dataset used for the pre-training and it is difficult for the users to directly assess the result quality, as only latent space embeddings are available in the pre-trained models.

The second category of methods is the cluster-based matching approach used in CellHint cell type harmonization [19] and FR-Match cluster-to-cluster two-way matching [16]. CellHint allows automatic cell type harmonization using predictive clustering tree and tree splitting algorithms. FR-Match utilizes a reduced feature marker gene space determined by the NS-Forest marker gene selection algorithm [27, 28], as a data-driven way to represent reference cell types in a maximally informative reduced dimensional space. For the query dataset, these methods utilize the upstream clustering results that group similar query cells together, making use of the aggregated properties of similar cells. When the query dataset is optimally clustered, cluster-based matching methods can be used to perform supervised matching between the query and reference datasets.

The cell-based methods were first benchmarked in a cross-validation setting using the HLCA dataset. Then, top performing cell-based methods and cluster-based methods were used to perform cell type matching between HLCA and CellRef. All results are cross-compared at the cluster-level after the cell-based matching results were summarized to the cluster-level.

Differences in the matching results were closely examined in the HLCA and CellRef UMAPs to resolve any discrepancies. Following this procedure, a collection of 68 cell types (41 matched cell types, 20 HLCA-specific cell types, and 7 CellRef-specific cell types) was collated into the resulting lung meta-atlas.

Last, we present a lung meta-atlas after integrating the cell types between HLCA and CellRef. We propose a strategy for incrementally growing the meta-atlas by adding new cell types from future studies while preserving the knowledge and cell types in the original atlases (**Figure 1C**). The incremental knowledge growth strategy will take advantage of the high-quality, community-approved v1 cell atlases with relatively coherent and validated cell type annotations. By performing meta-analysis comparison of the cell atlases, the meta-atlas can serve as a baseline for continued updating of the cell type collection from new single cell datasets and for adding new terms to the Cell Ontology [29–32] when there is sufficient evidence.

### Benchmarking of the cell-based methods on the HLCA dataset

The performance of Azimuth, CellTypist, scArches, scPred, singleR, and FR-Match cell-to-cluster matching methods were first benchmarked on the HLCA reference using cross-validation setting, where the HLCA cells were randomly split into 10 subsets (folds) within each cluster.

Each fold was then used as a query dataset to be assigned with the predicted annotations. After matching all folds, the predicted annotations of all cells were pooled and compared with the true annotations in the HLCA dataset (**Supplementary Figure 1**).

In the HLCA dataset, the abundances of the 61 cell types varied, with the alveolar macrophages being the most abundant cell type (68,487 cells) and the lymphatic EC proliferating cell being the least abundant (28 cells) (**Figure 2A**). Notably, 55% of cells are contained within the top 7 most abundant cell types, highlighting the large variation in cell abundances (cluster sizes) among the cell types. Potential effects of unequal cluster sizes when evaluating the matching performance are worth noting. For example, if training a model to perform perfect classification for the top 7 most abundant cell types in HLCA, the overall accuracy will be 0.55 (i.e., all cells in the top 7 cell types get correct classification), even if the remaining 54 out of 61 cell types all get misclassified. To mitigate the impact of the largely unequal cluster sizes, both overall performance metrics and per cluster performance metrics were calculated in the benchmarking comparison.

**Figure 2.**
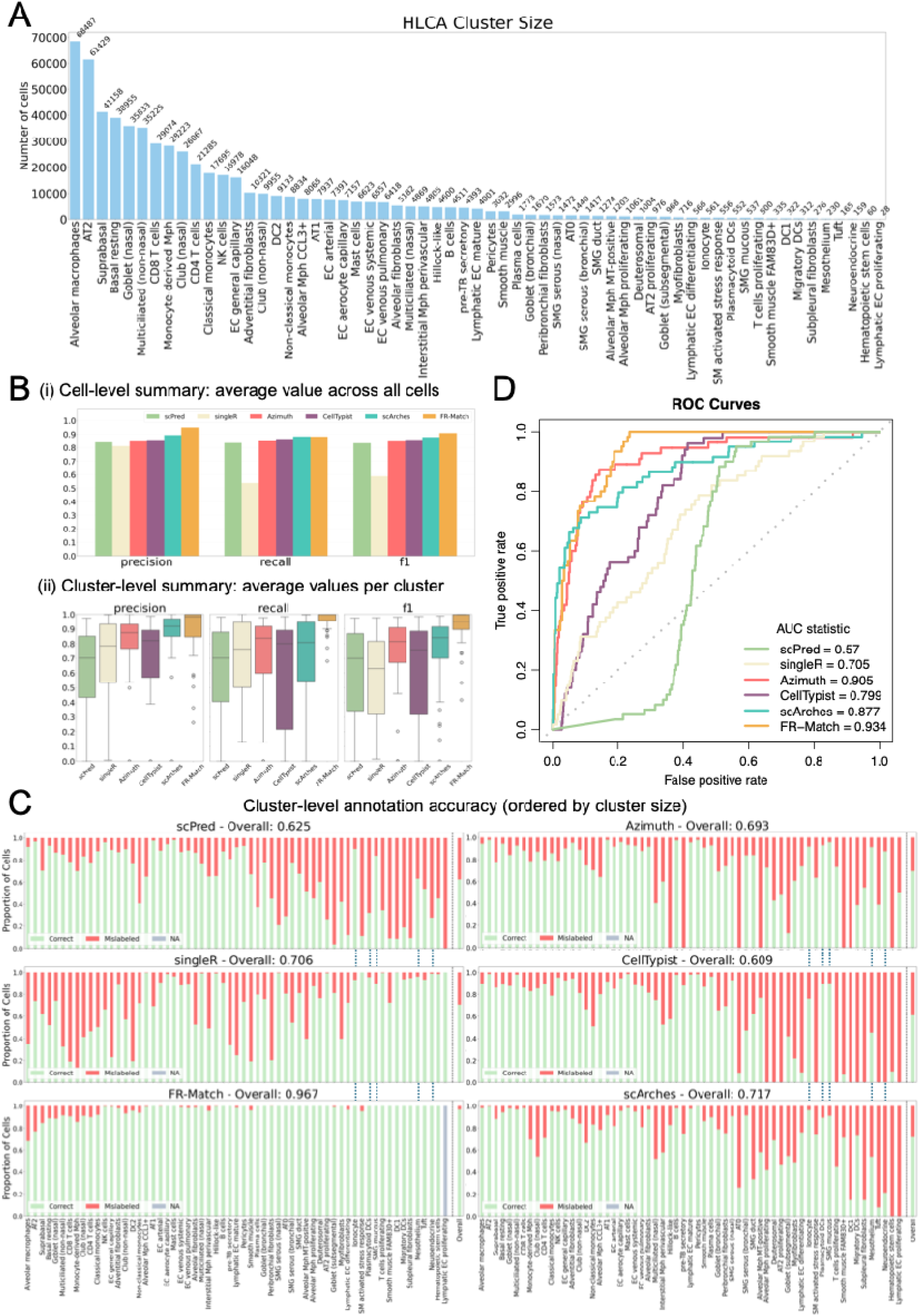
Benchmarking performance of cell-based annotation/matching methods in HLCA using 10-fold cross-validation. **(A)** Distribution of cluster sizes of the cell types represented in the HLCA dataset. **(B)** Precision, recall, and F1-score for each method (scPred, singleR, Azimuth, CellTypist, scArches, and FR-Match cell-to-cluster): (i) overall performance averaged by all cells and (ii) performance by each cluster. **(C)** Bar plots of cluster-level annotation accuracy for each method, sorted by cluster size with the proportion of cells correctly (green) or incorrectly (red) labelled. Due to a lack of cells in the smallest cluster after a 10-fold split (less than 3 cells in each fold), FR-Match filtered out this cluster as having too few cells for confident matching (see **Methods**). **(D)** ROC curve and AUC values for each method.

For benchmarking, each of the 10 folds of data was considered as the query set iteratively for predicting cell type annotations using the six cell-based methods. After pooling results for all cells by each method, the overall metrics in **Figure 2B(i)**, indicate that all methods achieved high overall precision (above 0.8) and five methods also had high recall and F1 scores. (SingleR was not benchmarkable with the other methods since it includes a fine-tuning step after running correlation analysis on all annotations, which disassociate the correlation-based confidence scores with predictions.) When these results were evaluated per cluster (i.e. grouped by each true HLCA cell type), the performance metrics showed variable patterns (**Figure 2B(ii)**). In contrast to the high overall F1-scores, the median F1-score when calculated per cluster for singleR (0.630), scPred (0.700), CellTypist (0.758), Azimuth (0.815), and scArches (0.841) were much lower than FR-Match (0.952); similar patterns were observed in precision and recall. The box plots of the per-cluster metrics for scPred, singleR, Azimuth, CellTypist, and scArches also showed wider ranges, suggesting more inconsistency in the cluster-level performance. The long whiskers and outliers in these box plots indicate that while the overall F1-score is high, certain clusters are low performing.

The variation of the cluster-level performance is further illustrated by the bar plots in **Figure 2C** which depict the proportion of correct-to-incorrect annotations made for each cell type, ordered by cluster size. FR-Match achieved the highest average accuracy (96.7%), followed by scArches (71.7%), singleR (70.6%), Azimuth (69.3%), scPred (62.5%), and CellTypist (60.9%). Looking at the trend with cluster sizes, scPred, Azimuth, CellTypist, and scArches follow similar patterns where larger clusters tended to have higher proportions of correct annotations and smaller clusters tended to have higher proportions of incorrect annotations. SingleR and FR-Match follows a reverse trend where larger clusters tended to have relatively higher proportion of incorrect annotations than smaller clusters, while the ratio of correct annotations for smaller clusters were remarkably better than the other methods. The relationship observed suggests that the discrepancy between the overall results and the cluster-level results in **Figure 2B** may be attributed to the high performance of larger clusters masking the low performance of smaller clusters for some methods. Therefore, when selecting the most appropriate metrics to evaluate the cell type annotation methods, it is important to not only evaluate overall performance but also to consider the method’s effectiveness in accurately identifying rarer cell types.

Utilizing the confidence scores produced by each of the methods, receiver operating characteristic (ROC) analysis was conducted in this cross-validation setting. Each of the six methods employs their own methodology for producing a confidence score to quantify a prediction’s certainty. Details about these methodologies are provided in the **Methods** section. The underlying assumption of a confidence score is that a method would assign higher confidence to the correct predictions and lower confidence to the incorrect ones, which can be used to construct ROC curves to assess the trade-offs between true positive rate and false positive rate at various confidence thresholds. FR-Match exhibited the highest performance with an area under the curve (AUC) score of 0.934, followed by Azimuth (AUC = 0.905), scArches (AUC = 0.877), CellTypist (AUC = 0.799), singleR (AUC = 0.705), and scPred (AUC = 0.57) (**Figure 2D**). To further understand the ROC results, colorized ROC curves are shown in **Supplementary Figure 2** and heatmaps of confidence scores are shown in **Supplementary Figure 3**. See **Methods** for more details. As an alternative to Figure 2C, the variation of cluster-level performance can also be viewed in **Supplementary Figure 4**, as heatmaps of the cell type–to–cell type confusion matrices for each method. The heatmaps in **Supplementary Figure 3-4** complement each other revealing the misclassification patterns between closely related cell types.

In summary, we found that (1) although almost all methods achieved high overall cell-based performance, the cluster-level performance varied from method to method, with lower performance on smaller clusters obscured in the overall evaluation; and (2) the ROC analysis demonstrated that method performance was driven and could be diagnosed by the confidence scores calculated in each method. Based on the cross-validation and ROC analysis, four of the cell-based methods – Azimuth, CellTypist, scArches, and FR-Match – were selected for further study.

### Performance on rare cell types

In **Figure 2C**, some rare cell types were well-resolved by many methods, e.g., ionocyte, plasmacytoid DCs, and SMG mucous, and some showed varying performance across methods, e.g., mesothelium and neuroendocrine. To understand the potential reasons, the feature spaces and cluster quality metrics were examined in **Supplementary Figure 5.** The unsupervised models follow the general steps of finding highly variable genes (HVG) and constructing latent spaces based on the HVGs for dimensionality reduction, whereas FR-Match uses the supervised feature space (i.e., gene space defined by NS-Forest, see **Methods**). Among the pre-trained models, only Azimuth has made its HVGs accessible in the reference data object.

**Supplementary Figure 5A** shows the 427 overlapping genes between the 3000 HVGs used by Azimuth and the 610 features (top 10 binary genes per cluster) used by FR-Match. It is expected that more features are selected as HVGs for larger clusters (i.e., cells from the larger clusters contribute more to the variance calculation) and significantly fewer features are selected as HVGs for smaller clusters (p-value = 0 by Spearman correlation test). This suggests that the HVG-derived latent space would unequally over-represent information in the larger clusters and down-represent information in the smaller clusters, thus, poorer performance in the rare cell types following the unsupervised HVG selection. **Supplementary Figure 5B** shows the average binary scores for the 10 binary genes used by FR-Match per cluster. Binary score is a metric that quantifies the on-and-off binary expression pattern of a gene in the target cluster and off-target clusters [27, 28], ranging from 0 to 1. The numerical difference of the binary scores among clusters is small, with an overall average score of 0.97. The rank-based correlation test with cluster size showed significance and the negative correlation value indicates consistently higher scores for rare cell types. The use of per-cluster binary scores for feature selection ensures a strong and fair representation of all cell types in the feature space used by FR-Match. **Supplementary Figure 5C-D** are metrics that could be used to understand the cluster quality and **Supplementary Figure 5E** shows the relationship between the two metrics. The F-beta score is a classification metric that quantifies the accuracy of its marker genes for the target cluster, ranging from 0 to 1, and the silhouette score is a widely used metric to evaluate the quality of clustering, taking values from −1 to 1. Considering both metrics, a well-resolved cell cluster would be expected to have good classification markers and a well-segregated cluster boundary in the UMAP visualization or an equivalent embedding. **Supplementary Figure 5C-D** show these two metrics are not impacted by cluster size. **Supplementary Figure 5E** shows there is no significant correlation between the two metrics, suggesting multiple independent measures should be considered when evaluating the cluster qualities. The highlighted clusters, however, had both high F-beta score and high silhouette score, which are rare cell types that had good performance in Azimuth but varying performance in CellTypist and scArches, among selected methods. In summary, this additional analysis suggests the potential reasons for why FR-Match showed better performance in the rare cell types and indicative metrics (such as F-beta score and silhouette score) that could partially explain the Azimuth performance in rare cell types but not the other methods. Future studies are needed to identify the root cause of the performance differences among the clusters with respect to cluster sizes in different methods.

### Benchmarking of the cell-based methods across datasets

The four selected cell-based methods were then used for matching of the CellRef dataset as query to the HLCA reference (CellRef → HLCA). Each query cell was assigned to a reference cell type or “unassigned” (see **Methods**), resulting in cell-level predictions for the 347,970 CellRef cells with corresponding confidence scores for each method (**Supplementary Table 1)**.

Again, cluster sizes had an impact on the matching results. In CellRef, there are 48 cell types, whose cluster sizes are shown in **Figure 3A**. The most abundant cell type is alveolar macrophage with 101,380 cells and the least abundant cell type is chondrocyte with only 6 cells. Due to inconsistent nomenclatures used to annotate cell types in the two studies, instead of validating the predicted HLCA-derived labels with the original CellRef labels, each cell’s prediction by one method was cross-compared with its corresponding predictions from the other three methods. An agreement score of 4 is given to a cell where all four methods gave the same prediction, 3 where three methods agreed upon the prediction, and so forth. In **Figure 3B**, 76.7% of cell predictions were agreed upon by all four methods, and an additional 14.7% showed a majority 3 out of 4 agreement. 8.4% of annotations were agreed between only two methods, and 0.2% had no agreement at all. While agreement among methods does not necessarily confirm the accuracy of the predictions, the high level of agreement across methods suggests that the predictions for most cells are probably correct.

**Figure 3.**
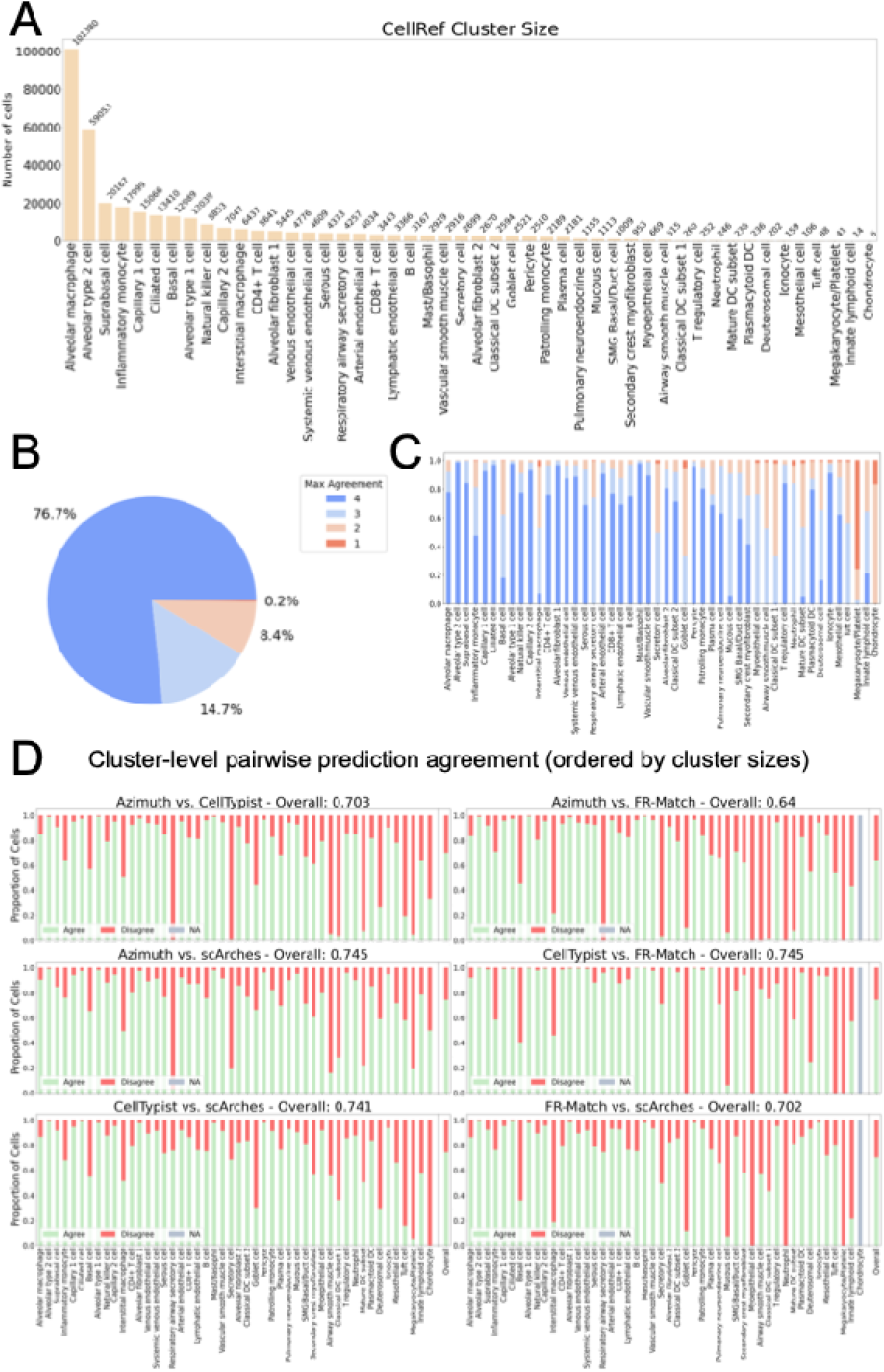
Benchmarking of the cell-based methods across the CellRef and HLCA datasets. The cell type annotation/matching is from CellRef as the query dataset to HLCA as the reference dataset (CellRef → HLCA). **(A)** Distribution of cluster sizes of cell types represented in the CellRef dataset. **(B)** Summary of level of prediction agreement across all methods (Azimuth, CellTypist, scArches, and FR-Match cell-to-cluster) at the cell-level. **(C)** Stacked bar plot depicting level of prediction agreement across all methods at the cluster-level, ordered by cluster size. **(D)** Proportion of agreement between two methods’ predictions for each cluster, ordered by cluster size. Note that chondrocyte was filtered out in FR-Match due to its small cluster size.

As observed above, it would be informative to look at the prediction agreement at the cluster-level. As expected, when looking at the agreement levels for individual clusters, there was a trend where larger clusters had higher agreement, and smaller clusters had lower agreement (**Figure 3C**). In this figure, cells are grouped by their original CellRef labels. Looking at the most abundant cluster, 91.56% of alveolar macrophage cells’ predictions had majority agreement among all four methods (score of 4 or 3). As this cluster consists of 29.13% of the dataset, alveolar macrophage and other large, high agreement clusters may have biased the overall cell-level results. Similarly, all four methods assigned the same predictions for almost all cells in abundant alveolar type 2 cell and alveolar type 1 cell clusters (98.38% and 97.51% of cells had a score of 4, respectively). In contrast, there was no majority agreement (score of 2 or 1) in clusters such as megakaryocyte/platelet and chondrocyte. Although the 0.2% of cell-level predictions that had no method agreement (score of 1) is a low number, most of those predictions are found within the megakaryocyte/platelet and chondrocyte clusters (containing 41 and 6 cells respectively). Both clusters, although small, have high proportions of little-to-no method agreement (score of 2 or 1) for its predictions (97.56% and 100% respectively), which further points out the difficulties in accurately predicting rare cell types.

Assessing the pairwise agreement between methods revealed additional insights into the performance for individual methods (**Figure 3D**). Overall, a general trend between cluster size and prediction agreement was observed for each pair, where the larger clusters tended to have more agreements. While there was a higher proportion of disagreement (red) towards the right of these plots, there was also low agreement for some moderate sized clusters. Some disagreement could be attributed to certain methods. For example, Azimuth showed a high disagreement with all other methods about predictions for respiratory airway secretory cell and secretory cell; in contrast, the other method pairs have high agreement on the prediction for these two cell types. Thus, there is no agreement score of 4 in **Figure 3C** for these two types. While we cannot tell from these plots whether Azimuth correctly predicted for these two cell types while other methods were incorrect, or vice versa, this example demonstrates how even among similarly performing algorithms, methods may still vary in their ability to predict specific cell types, regardless of how well the clusters are represented in the dataset. In the case of basal cell and intestinal macrophage types, which have sufficient cells (12,989 and 6,437 cells respectively), relatively low and variable agreement was observed between method pairs, especially in the pairs of FR-Match and others, resulting in an agreement score of 4 with only 17.92% and 7.49% of cells (**Figure 3C**). The matching results of basal cell and intestinal macrophage types will be further explored in the next section. Interestingly, the highest overall agreement achieved in this pairwise comparison are the pair of Azimuth and scArches (0.745), and the pair of CellTypist and FR-Match (0.745), suggesting some similarity in these pairs of methods.

### Cell-based matching results of the lung cell atlases

Cell-based matching of CellRef as query and HLCA as reference results in each CellRef cell being assigned with an HLCA cell type label prediction. **Figure 4A** shows UMAP plots in which CellRef cells in the CellRef UMAP coordinates are colored by the HLCA label predictions by each method. Overall, Azimuth predicted 54 cell types out of the 61 HLCA cell types; CellTypist predicted 47; scArches predicted 61; and FR-Match predicted 49. While the original CellRef contains 48 cell types of its own annotation, the predictions to all 61 HLCA cell types by scArches may suggest that the scArches model is overfitted as it was trained by the HLCA team. Indeed, some of the cell types reported in the HLCA dataset were exclusively derived from the nasal swab specimens, which were not used in the CellRef study. While a global evaluation of these UMAPs (i)-(iv) suggest that the matching results are very similar (**Figure 4A**), since 76.7% of cells had agreed predictions by all four methods (**Figure 3B**), the differences between the methods require more careful examinations.

**Figure 4.**
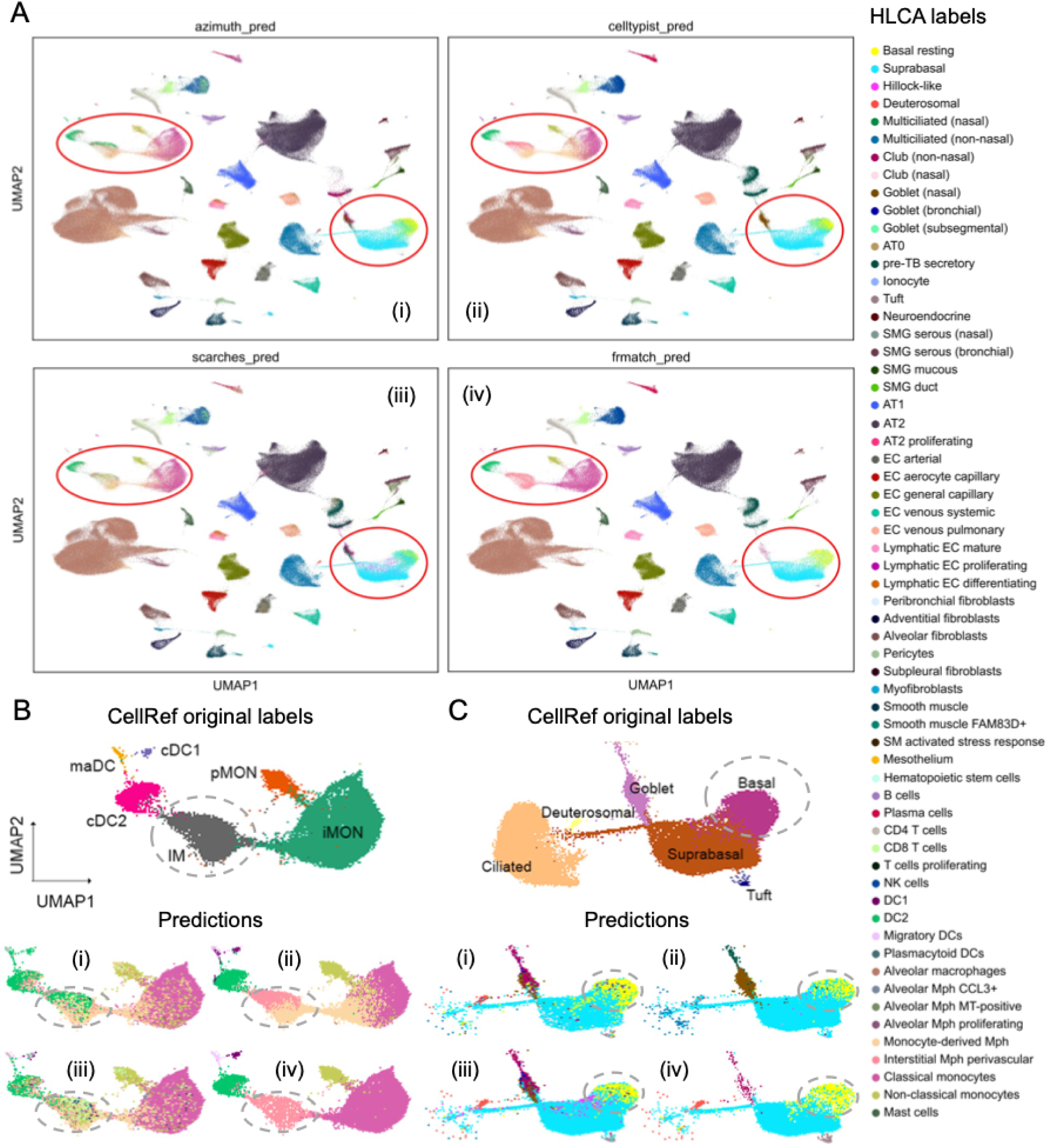
UMAP of cell-based matching results of CellRef cells to the HLCA reference. **(A)** CellRef cells in the CellRef UMAP embeddings are colored by HLCA labels predicted by Azimuth (i), CellTypist (ii), scArches (iii), and FR-Match cell-to-cluster (iv). Red circles highlight regions with major differences. **(B-C)** Detailed views of the highlighted regions in (A) for comparison of the CellRef original labels and the HLCA predictions for these cells. Methods used in (i)-(iv) are the same as in (A). The dotted circles indicate the major differences in interstitial macrophage (IM) and basal cell. Cells with the original label of cDC1, maDC, cDC2, IM, iMON, pMON are plotted with predictions in the “IM” region plot in (B). Cells with the original label of deuterosomal, goblet, suprabasal, tuft, and basal are plotted with predictions in the “basal” region plot in (C).

Two regions where major differences were observed are highlighted in **Figure 4A** and shown in detail in **Figure 4B-C**, where the original CellRef labels and the predictions are compared. One region is the interstitial macrophage (“IM”) region in CellRef (**Figure 4B**), where the well-segregated IM cluster is matched to a mixture of HLCA cell types by Azimuth and scArches, to two HLCA cell types split into roughly equal sizes by CellTypist, and to one HLCA cell type uniformly colored by FR-Match. The FR-Match result supports the well-segregated shape of the query cluster (interstitial macrophage in CellRef) and suggests a uniform matching to the interstitial Mph perivascular in HLCA. The CellTypist result is also interesting, where the beige color in the predictions is the monocyte-derived Mph in HLCA, suggesting a transitional state of cell types from the inflammatory monocyte (iMON) to interstitial macrophage (IM) in CellRef.

The other region showing major differences is the “basal” region in CellRef (**Figure 4C**), where there is a separation of suprabasal and basal in the CellRef original labels, but spillover of suprabasal (cyan color) into the “basal” region (yellow color) in the predictions. The FR-Match result had the least spillover of the cyan color because those spillover cells were likely of low confidence, thus were “unassigned” using FR-Match (**Supplementary Figures 6 and 7**). All predicted labels as well as the original CellRef and HLCA labels for the “IM” and “basal” regions are in **Supplementary Figure 6**, where the “unassigned” cells in FR-Match are colored white, implying a clean-up of the low confidence cells.

In **Supplementary Figure 7A**, cells in the UMAP plots are colored by their confidence scores by each method. While most of the cells had high confidence scores in each plot, all methods produced weakly matched cells spread over many clusters. FR-Match produced some regions packed with very low confidence cells (i.e., dark islands in the plot), including the spillover “basal” region, where many of the spillover cells are “unassigned” (**Supplementary Figure 7B**).

Labeling the spillover cells as “unassigned”, reflecting the low matching confidence, is more informative than a fuzzy matching to a similar label, e.g., suprabasal. Another “unassigned” pattern in the FR-Match plot (**Supplementary Figure 7B**) is highlighted by the black arrows where the “unassigned” cells form an entire cluster that is isolated from others, suggesting CellRef cell types that are not in the HLCA reference. Lastly, due to a different version of HLCA annotations being used in the Azimuth training model, the Azimuth-only labels had decent confidence scores (**Supplementary Figure 7A-B**), suggesting an update is needed for the Azimuth lung model to be consistent with the cell type annotation in the HLCA literature.

### Cluster-level analysis of the cell-based matching results

As previously shown, the cluster-level pairwise prediction agreement varied a lot among clusters (**Figure 3D**). By aggregating the cell-level predictions per query cluster, **Figure 5** shows the proportion of CellRef cells matched between a pair of the original CellRef cell type (horizontal axis) and the predicted HLCA cell type (vertical axis) for each method. The HLCA cell types are ordered by the hierarchical clustering such that similar clusters are grouped next to each other according to their gene expression profiles (**Methods**). The CellRef cell types are ordered accordingly so that the highest matched proportions would ideally align along the diagonal. These results are also visualized in **Supplementary Figure 8**, as heatmaps of the cell type–to–cell type confusion matrices.

**Figure 5.**
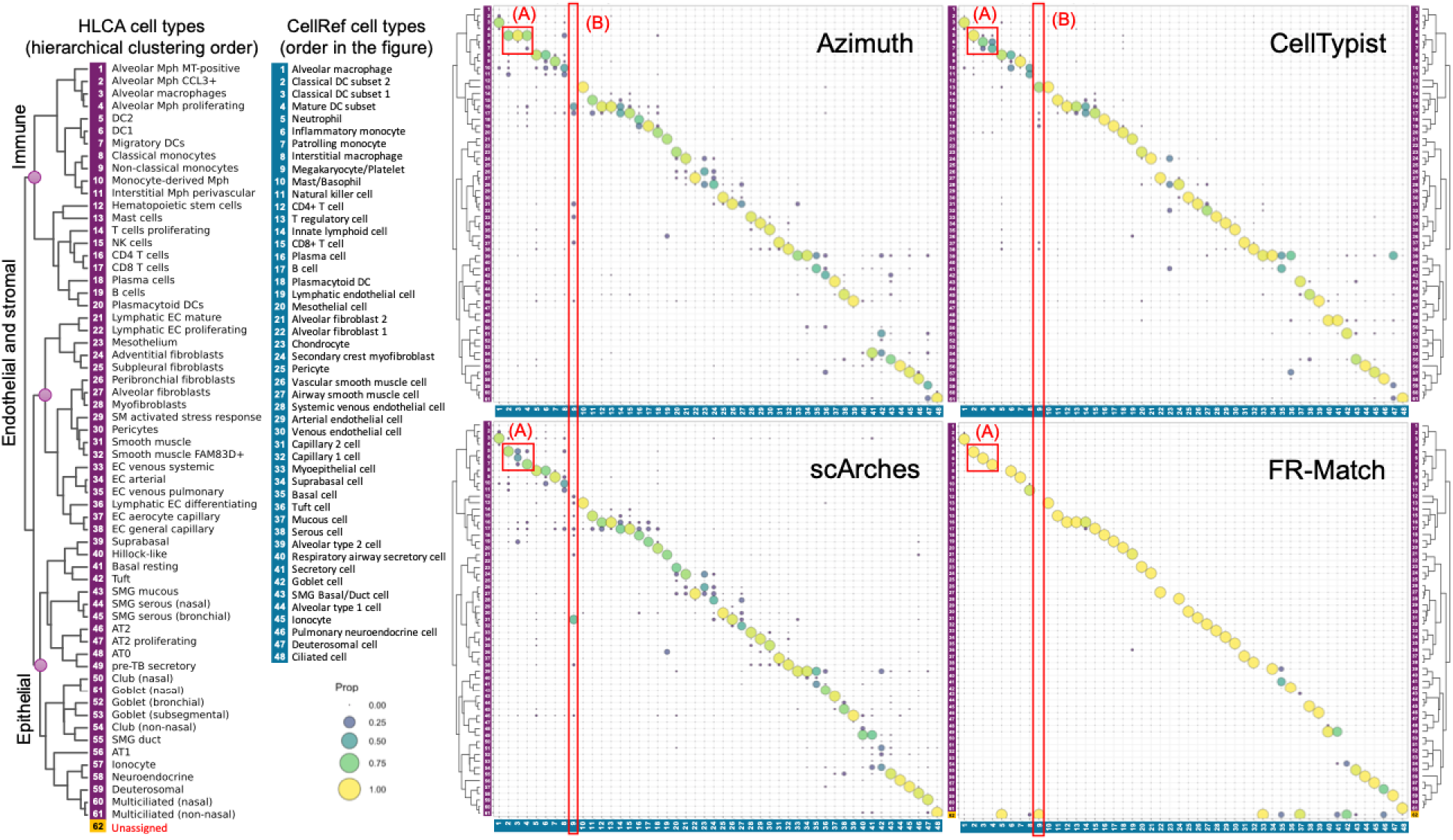
Matching between the 48 cell types in CellRef and the 61 cell types in HLCA. Cell-based matching results are summarized at the cluster-level. The 48 CellRef cell types are on the horizontal axis; the 61 HLCA cell types are on the vertical axis. Each circle in the plot is the proportion of cells in the original CellRef cluster matched to each of the corresponding HLCA clusters (CellRef as query to HLCA as reference), so that the proportions in each column sum up to 1. The dendrogram to the left shows the hierarchical clustering order of the HLCA cell types, which groups similar cell types near each other in the dendrogram based on their transcriptional profiles (**Methods**). The CellRef cell types are ordered according to the HLCA dendrogram order so that the matching proportions in the figure align close to the diagonal; an off-diagonal circle suggests a mismatch. In the FR-Match figure, “unassigned” cells are collected in the bottom row, indicating the statistically weak matching of these cells to all HLCA clusters. The highlighted boxes are examples of different matching patterns by the methods for dendritic cell types (A) and for “unassigned” cells (B).

In **Figure 5**, FR-Match showed the cleanest aggregated cluster-level matching pattern with the lowest off-diagonal proportions. Two examples are highlighted to reveal the different matching patterns among the methods. In the highlighted box (A), the three dendritic cell (DC) types of CellRef are matched differently by all methods. Azimuth matched all three types to DC2 in HLCA; CellTypist clearly matched classical DC subset 2 to DC2 but not the other two types; scArches showed relatively smaller proportions of matched cells, with some matched to other DC types and some matched to non-DC types; and FR-Match identified unique 1-to-1 matches for each of the three types.

The highlighted box (B) illustrates the matching pattern for an “unassigned” cell type by FR-Match. For the megakaryoctye/platelet type in CellRef, it was matched poorly to many cell types in HLCA by Azimuth and scArches and was matched to the HLCA mast cells by CellTypist. Only FR-Match determined that those poorly matched cells should be “unassigned”. The disagreement among Azimuth, CellTypist and scArches also indicates that there is no consistent match to the megakaryoctye/platelet type and should be “unassigned”. These examples suggest that further refinements are needed to fine tune the pre-trained models of Azimuth, CellTypist, and scArches for more accurate matching (e.g., for the DC types) and to address the scenario where weakly matched cells are forced to match to some cell type in the reference.

### Cluster-based matching of cell types using FR-Match and CellHint

FR-Match also performs cluster-to-cluster matching, which allows two-way matching (i.e., from CellRef as query to HLCA as reference and vice versa, denoted as CellRef ← → HLCA), by utilizing the cell type annotations in both datasets. The two-way matching results are shown in **Figure 6A**, and results of each direction are shown in **Supplementary Figure 9**. The two-way matches indicate that the pair of the matched CellRef and HLCA clusters are found by using both the CellRef as reference and the HLCA as reference. In **Figure 6A**, the two-way matches found by the FR-Match cluster-to-cluster matching strategy exhibit a very clean matching pattern, where most of the two-way matches are 1-to-1 matches and only a few are 1-to-2 or 2-to-1 matches. The cluster-to-cluster results are easier to interpret and not impacted by the varying cluster sizes as each cluster is assessed as a whole in the cluster-based method.

**Figure 6.**
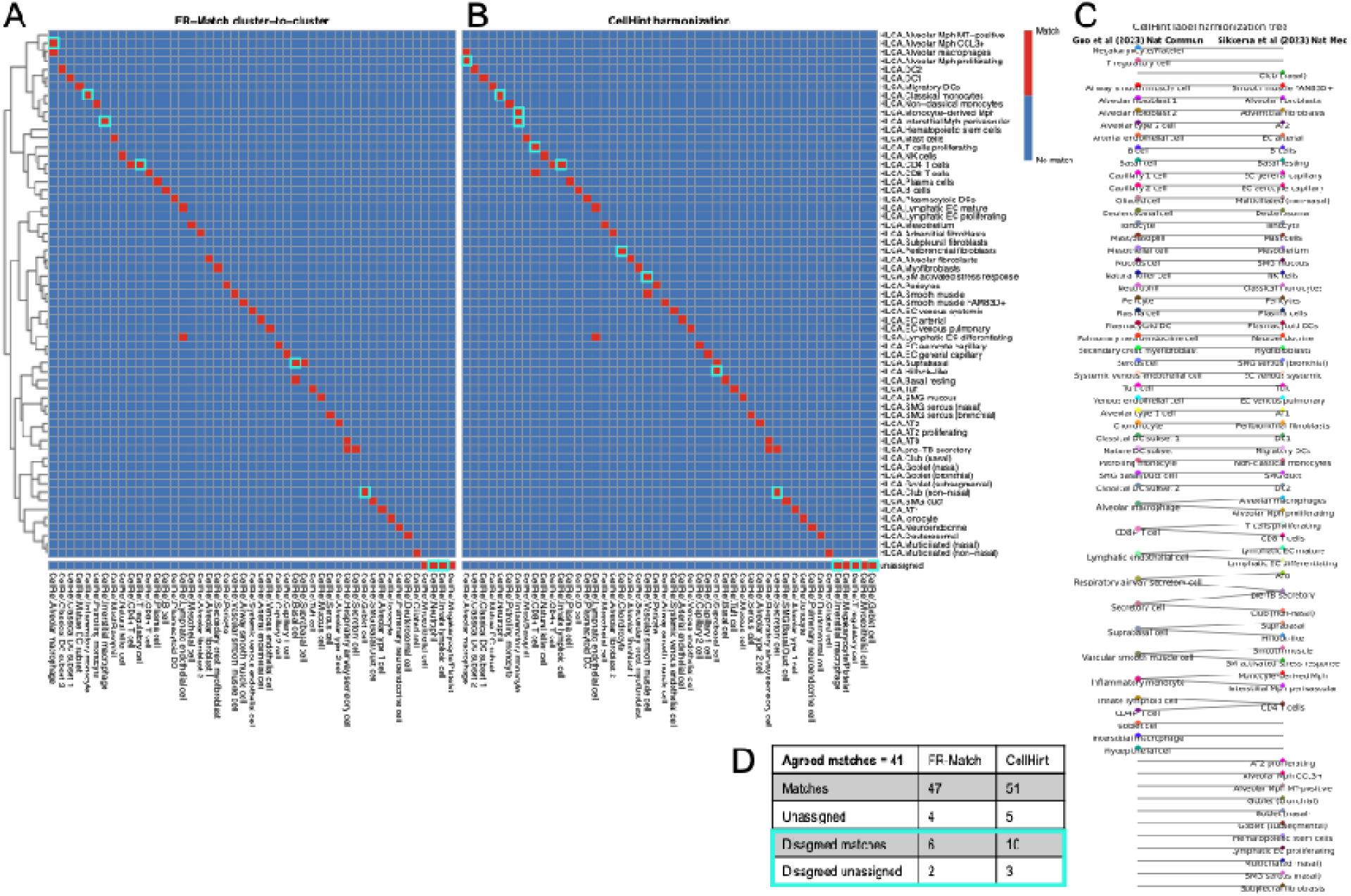
Cluster-based matching of healthy lung cell types. **(A)** FR-Match cluster-to-cluster results of the CellRef ← → HLCA cell type matching. Two-way matches of the CellRef and HLCA cell types. CellRef as query to HLCA as reference and HLCA as query to CellRef as reference were both performed by FR-Match. The red blocks indicate two-way matches where the matched/unassigned cell types are agreed by both matching directions. **(B)** CellHint harmonization results of CellRef and HLCA. Rows are the HLCA cell types ordered by the dendrogram. Columns in (A) and (B) are ordered differently, each optimizing the matches along the diagonal and grouping the unassigned at the end. **(C)** Tree plot outputted from CellHint. Each red block in (B) correspond to an edge in (C). **(D)** Total count table and disagreements between FR-Match and CellHint. Cyan boxes correspond to disagreements.

Although it is possible to have one-to-many or many-to-one matches, there are fewer of these in the cluster-based results than in the cell-based results. It is worth noting that the “unassigned” row at the bottom also shows two-way matches, suggesting that these cell types are only found in the CellRef atlas. Similarly, the empty rows in the figure indicate the cell types that are only found in the HLCA atlas.

CellHint, a sister method to CellTypist, conducts guided cell type harmonization across cell atlas datasets [21]. **Figure 6B** shows the CellHint matches for the CellRef and HLCA harmonization, visualized analogous to **Figure 6A** to ease comparison, converted from CellHint’s tree plot visualization in **Figure 6C**. Differences between the FR-Match and CellHint results are highlighted in the cyan boxes. **Figure 6D** summarizes the total match and unassigned counts and their disagreements between the two methods. CellHint tends to result in more matches than FR-Match. The two methods had 41 agreed matches and 2 agreed unassigned types (myoepithelial cell and megakaryocyte/platelet, both of which are rare cell types in CellRef). The agreement between the two methods is substantial (87% of the FR-Match matches and 80% of the CellHint matches). Some of the disagreements could help disambiguate the 1-to-2 or 2-to-1 matches, reciprocally. For example, the reciprocal results suggests that the unique 1-to-1 match of CellRef alveolar macrophage and HLCA alveolar macrophages is correct, thus ruling out the other match in the 1-to-2 matching patterns in each other’s matching results. Compared to the cell-based matching approach, the cluster-based matching approach can produce more concise results, which are much easier to compare and interpret, focusing on the cluster as a whole instead of the individual cells, while the later might be prone to the clustering uncertainty to various degrees.

### Recommended matching for the lung cell type meta-atlas across CellRef and HLCA

For the lung cell atlases, a comprehensive meta-analysis of benchmarked cell-based matching results (**Figure 5**) and the cluster-based results (**Figure 6**) was conducted to construct a meta-atlas of lung cell types across CellRef and HLCA. For the cell-based results, a computational strategy was applied to determine high confidence matches based on the matching proportions (**Methods**), so that all cell-based results could be uniformly processed into a binary “match” (Y) or “no match” (N) format between the cell types. Then, the two-way cluster-to-cluster matches from FR-Match and cell type harmonization from CellHint were combined with the high confidence matches from the cell-based matching results to provide evidence for determining the recommended matching in the meta-atlas. **Figure 7** shows the cross-comparison table for the recommended matching of the cell types between CellRef and HLCA. Candidates of “HLCA2CellRef_Match” are suggested by the agreement of the cluster-based matching method and at least one cell-based matching method; the remaining are candidates for atlas-specific cell types. All candidate pairs and any match to the atlas-specific candidates by a single method were carefully examined by looking at their marker gene expression patterns using the marker gene selection algorithm NS-Forest [27, 28] (**Methods**) before determining the final matching recommendation. 28 matches were identified by all methods with all Y’s; the remaining were determined by majority voting and examination on the marker gene expression pattern. The complete matching table with characterizing marker genes is in **Supplementary Table 2**.

**Figure 7.**
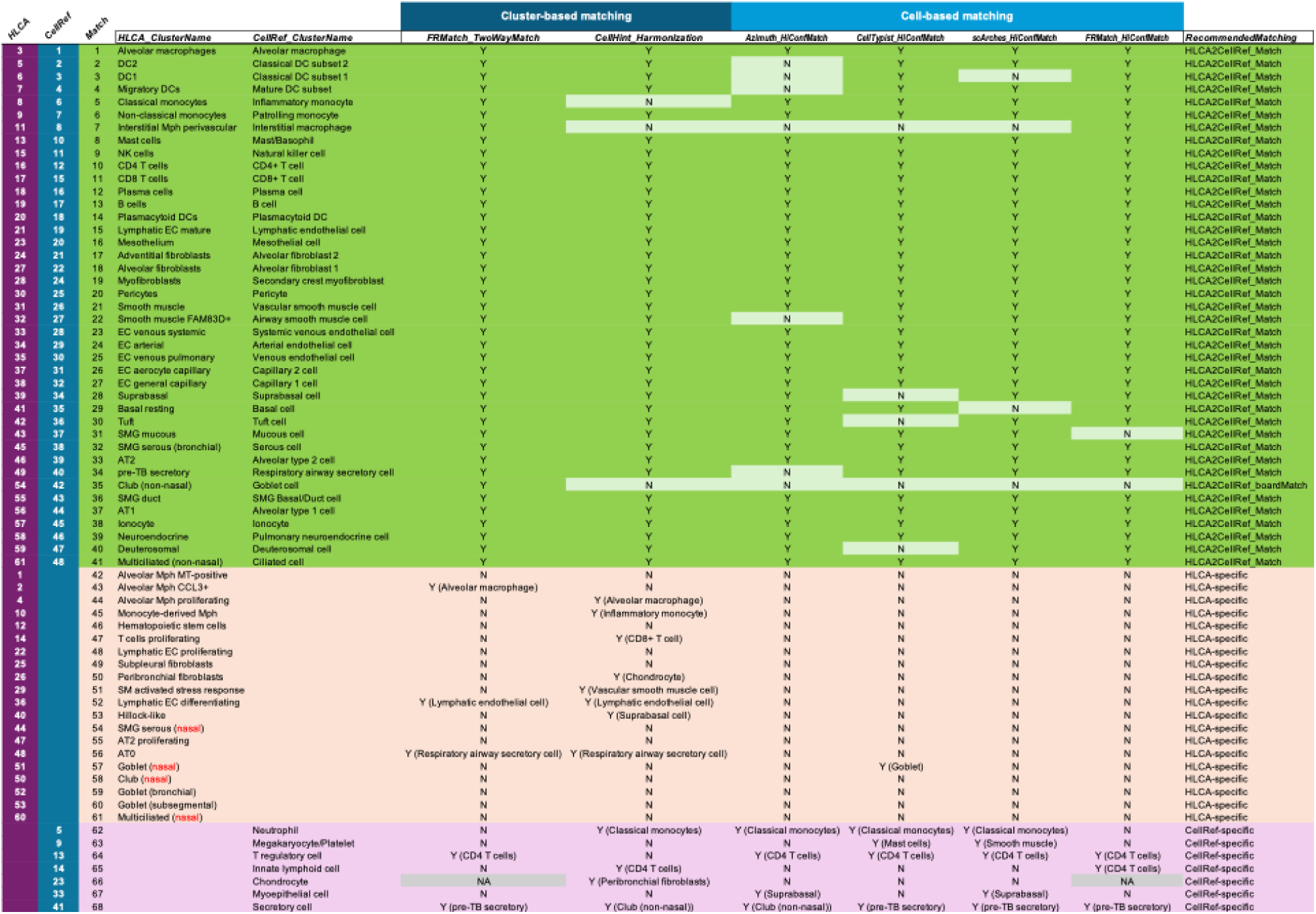
Recommended matching of the lung cell types between CellRef and HLCA. 41 recommended cell type matches (cluster-based matching + cell-based matching) out of 61 HLCA cell types and 48 CellRef cell types. An additional 20 cell types are unique to HLCA and 7 cell types are unique to CellRef.

**Figure 8** shows the use of NS-Forest marker genes to disambiguate potential matches. The NS-Forest marker gene combinations form unique “barcode” (i.e., on-and-off) patterns to characterize different cell types in a scRNA-seq dataset. **Figure 8A** are the “barcode” plots of the HLCA AT0 and pre-TB secretory types and the CellRef respiratory airway secretory cell and secretory cell types. The four cell types form three pairs of matches in both FR-Match and CellHint cluster-based matching results (**Figure 6A-B**). Using the “barcode” plots, the expression combination of the three marker genes – *SCGB3A2*, *SCGB1A1*, and *SFTPB* can clearly delineate the recommended matching pair – HLCA pre-TB secretory and CellRef respiratory airway secretory cell – from the other two types. The matched pair have high expression in all three markers (i.e., *SCGB3A2*^high^*SCGB1A1*^high^*SFTPB*^high^), whereas the HLCA AT0 is *SCGB3A2*^high^*SCGB1A1*^low^*SFTPB*^high^, and the CellRef secretory cell is *SCGB3A2*^low^*SCGB1A1*^high^*SFTPB*^low^ in the corresponding datasets. Another example is the CellRef regulatory T cells, which is recommended as a CellRef-specific type, despite the potential matching to the HLCA CD4 T cells by all methods. Although these are very similar cell types, markers like *FOXP3* and *CTLA4* can distinguish the T regulatory cells (**Figure 8B**). A special case of “HLCA2CellRef_broadMatch” is recommended between the HLCA club (non-nasal) and CellRef goblet cell types, because the CellRef goblet cell type is likely under-partitioned, containing both club cells expressing *SCGB1A1*, the secretoglobin 1A1 (also known as club cell secretory protein) [33], and goblet cells expressing the well-known marker *MUC5AC* [34] (**Figure 8C**). In these two atlases, there are three location-specific goblet types (nasal, bronchial, and subsegmental) in HLCA, thus none of which were matched to the under-partitioned goblet cell type in CellRef. There is no club cell annotated in CellRef, which is likely grouped in the goblet cell type in the CellRef atlas. It is also worth noting that none of the four “nasal” cell types in HLCA are matched to anything in CellRef in the cross-comparison table (**Figure 7**), which is consistent with the fact that nasal specimens were not included in the CellRef experiments. However, that was not the case in the individual matching results of Azimuth, CellTypist, and scArches (**Figure 5**), where nasal cell types were weakly matched to similar cell types in different tissue locations. In contrast, cell-based FR-Match did not match “nasal” cell types (**Figure 5-6**). By using the supervised feature space of NS-Forest markers, the FR-Match results are easily interpretable. “Barcode” plots of matched types can be used to understand and validate the matching results. In **Supplementary Figure 10**, the marker gene profiles show highly consistent patterns across datasets. For example, the HLCA markers of *AGER* for AT1, *SFTPA1* for AT2, *KRT15* and *KRT17* for resting basal, and *HIGD1B* and *KCNK3* for pericytes show very high specificity in the matched CellRef types; and vice versa. Benchmarking of the NS-Forest markers and markers sets selected by other methods can be found in our earlier study [28]. More description of “barcode” plots is in **Methods**.

**Figure 8.**
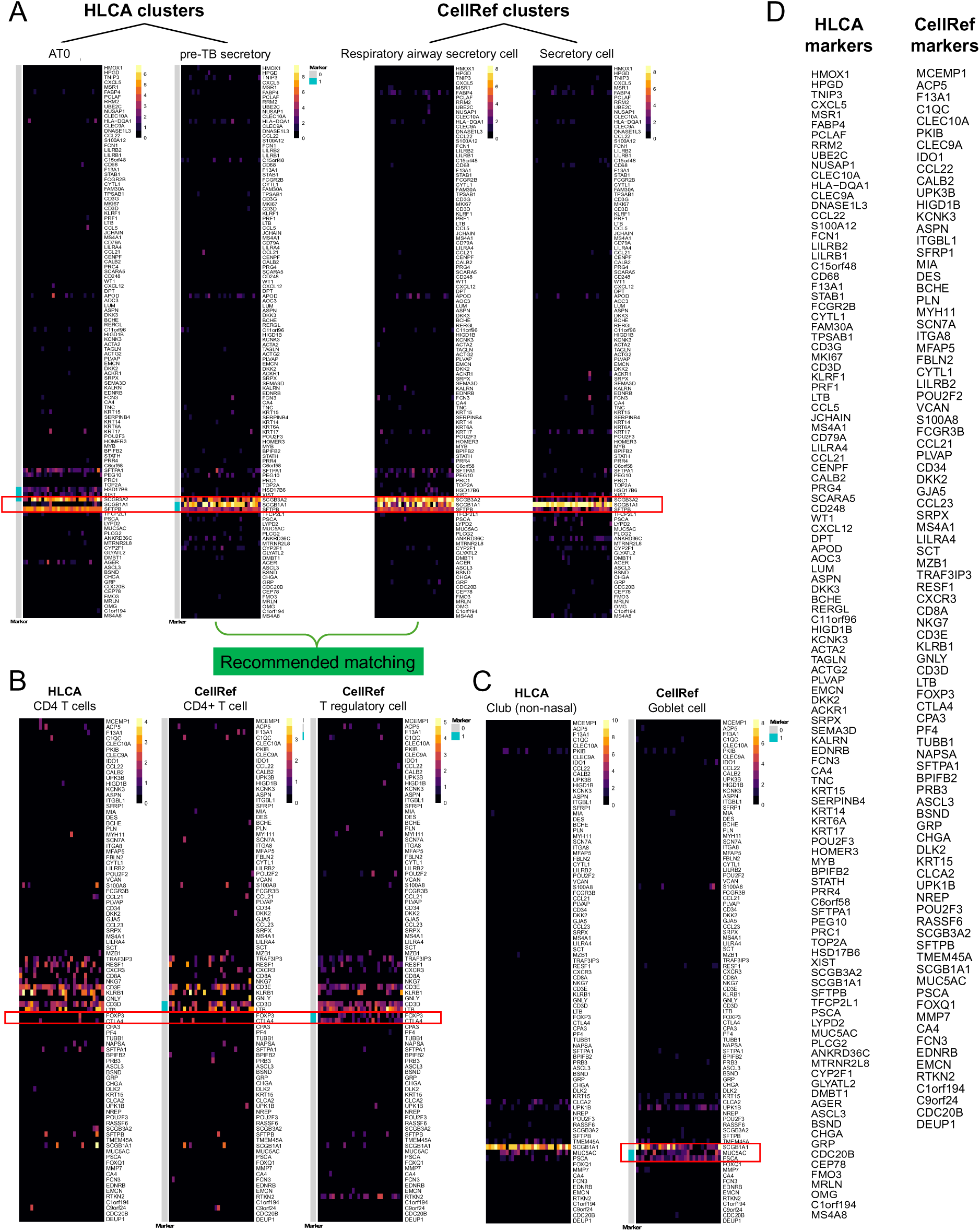
Use of NS-Forest marker genes to disambiguate potential matches. Each heatmap shows the gene expression pattern of 30 randomly selected cells (columns) from the plotting cell type in the NS-Forest marker gene space (rows). **(A)** The recommended matching of HLCA pre-TB secretory and CellRef respiratory airway secretory cell are *SCGB3A2*^high^*SCGB1A1*^high^*SFTPB*^high^, whereas the HLCA AT0 is *SCGB3A2*^high^*SCGB1A1*^low^*SFTPB*^high^, and the CellRef secretory cell is *SCGB3A2*^low^*SCGB1A1*^high^*SFTPB*^low^ **(B)** Distinction of T regulatory cells from CD4+ T cells by markers *FOXP3* and *CTLA4*. **(C)** The CellRef goblet cell type is likely under-partitioned, containing both club cells expressing *SCGB1A1* and goblet cells expressing *MUC5AC.* **(D)** NS-Forest marker gene spaces of HLCA and CellRef.

In summary, out of the 61 HLCA cell types and 48 CellRef cell types, the meta-analysis recommends 41 matched cell types, 20 HLCA-specific cell types, and 7 CellRef-specific cell types. Taking these results together, it suggests that an integrated version of these two atlases – a meta-atlas – with 68 discrete cell types would serve as a higher-quality and more comprehensive reference for normal human lung cell types. This workflow demonstrates a framework that supports the incremental cell type knowledge growth.

### Incremental cell type knowledge growth in practice

Alongside the continued efforts of the cell atlas consortia to release fully integrated datasets, a greater community effort among many consortia is to operationalize the FAIR Data Principles [35]. For example, the HRA and brain atlasing efforts are making substantial changes to the Cell Ontology (CL) to ensure stable cell types are covered [31, 32] and Provisional Cell Ontology (PCL) was developed specifically to hold cell types that are provisionally defined by single cell transcriptomics atlases [36, 37]. The ontological framework of capturing and disseminating the data-driven cell phenotype knowledge is one way to ensure that the annotated data in previous and future publications are interoperable in the evolving context.

The incremental cell type knowledge growth strategy based on the benchmarking framework provides a practical way of expanding the cell type knowledge based on single cell data. We are in the process of building a data-driven knowledgebase about cell phenotypes. The general steps include rigorous cluster quality control using the silhouette score and F-beta score, computationally matching the high-quality cell type clusters using a selective set of methods based on the benchmarking performance, and an end-to-end extract-transform-load pipeline for a graph knowledgebase. More information can be found in [REF will be available in April].

For example, the lung meta-atlas results in a knowledge graph that consists of cell sets (i.e. clusters in a dataset) as nodes, and their relations with other entities in edges (**Figure 9**).

**Figure 9.**
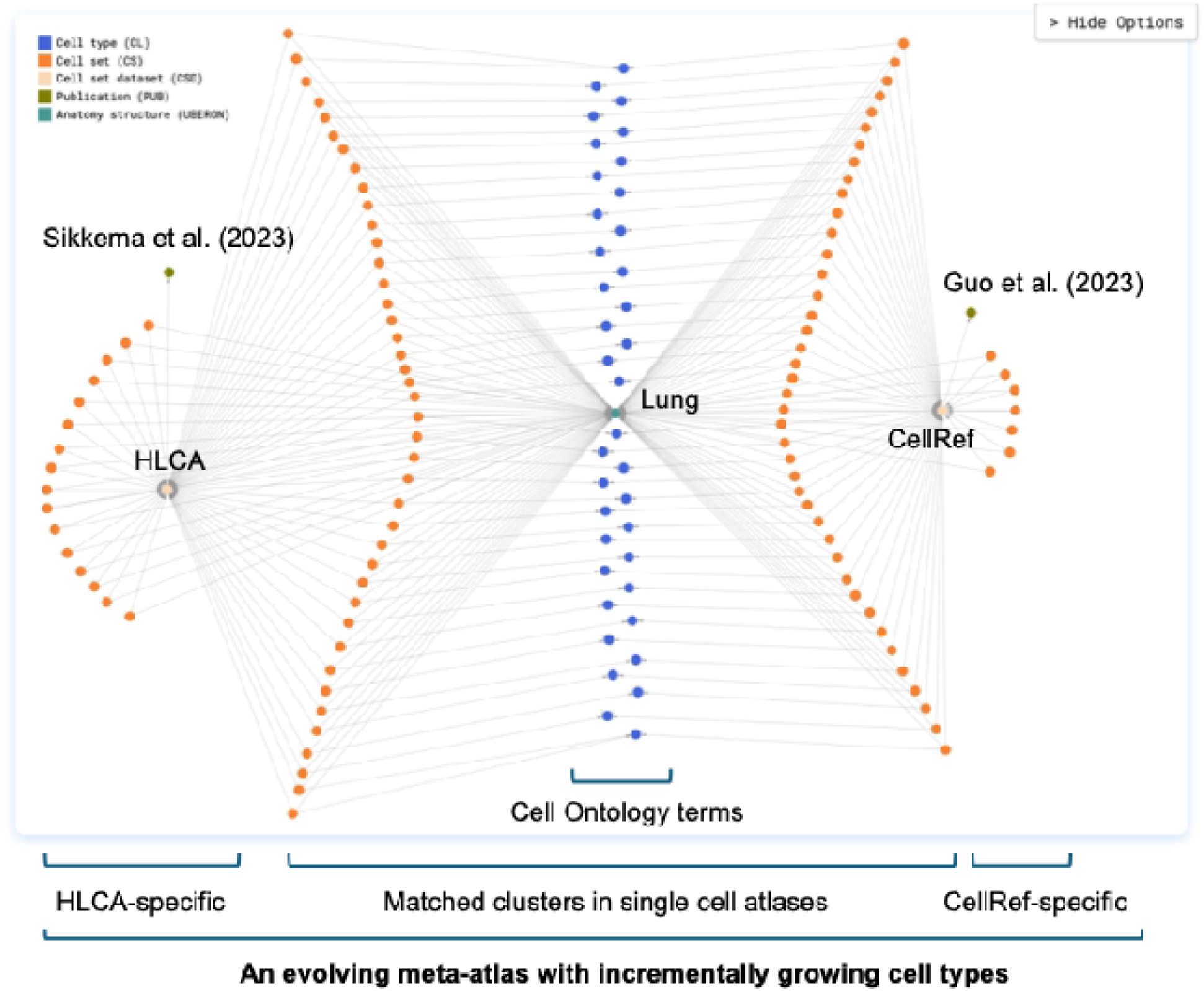
An example of incremental cell type knowledge growth in practice. The knowledge graph captures the knowledge derived from single cell experiments in the cell set, cell set dataset, and publication nodes and connects with existing knowledge in biomedical ontologies in the cell type nodes (Cell Ontology) and anatomy structure node (UBERON), showing a collection of matched cell types and specific cell sets emerged from the cell atlases. Cell set node: a cell cluster in a cell atlas dataset. Cell set dataset node: a cell atlas dataset. Cell type node: a Cell Ontology term linked with Cell Ontology identifier.

Linking cell sets to CL terms is often done manually and is a time-consuming process. Based on the meta-atlas, the computationally matched cell sets would automatically inherit the CL identifiers from an earlier linked dataset, and only the unmatched cell sets need to be assigned with a CL identifier. In this way, the knowledge graph preserves the provenance of each cell set from its original publication, and allows the organic growth of knowledge when new datasets are added, ensuring the interoperability (i.e., uniformly assigned ontology identifiers) and reusability (i.e., from previous publications to future publications) of data with respect to the evolving nature of the cell atlases.

### Method benchmarking on kidney datasets

The same methods were also applied to match two kidney cell atlases in their most current versions. Lake et al. (2023) presented an atlas of healthy and injured cell states in human kidney (Human Kidney Atlas 1.0), where 75 cell types were defined across conditions [17]. More recently, Acera-Mateos et al. (2025) generated a multimodal benchmarking dataset for renal cortex (mBDRC) using kidney tissues from “healthy” donors undergoing nephrectomy [18]. The Acera-Mateos et al. is currently a preprint and has 36 annotated cell types. The Lake et al. dataset was used as the reference in the cell type matching.

Using the same methods, the cell-based matching results are shown in **Supplementary Figure 11,** and the cluster-based matching results are shown in **Figure 10**. As before, the cell-based matching results showed large variations from method to method, and the cluster-based results were easier to interpret, conveying consensus results from FR-Match and CellHint. For the pre-trained models, one limitation is that it depends on the version of the reference that the model is pre-trained on, so that it needs regular updating if the reference dataset is updated. An example is the Azimuth kidney model. According to the documentation, an earlier preprint version of the Lake et al. data was used as the reference, and it is unclear if diseased cells were used or not. Therefore, the Azimuth results had many fewer cell types and different names in **Supplementary Figure 11**. CellTypist and scArches pre-trained models for kidney are not available and so they were trained using the Lake et al. publication dataset as reference in this analysis. For these reasons, the results from the cell-based unsupervised label transfer models (Azimuth, CellTypist, and scArches) showed large variations from each other. The FR-Match cell-to-cluster result showed more concentrated matching in many clusters (i.e., high proportion of query cells from the query cluster matched to the same reference cluster) but also showed uncertainties in some other clusters. On the other hand, using cluster-based matching, 25 agreed matches (21 unique 1-to1 matches and 2 pairs of 1-to-2 matches) were found between FR-Match two-way matches and CellHint harmonization result (**Figure 10**). Considering the pre-print state of the Acera-Mateos et al. (mBDRC) dataset, we might expect that some of those clusters would benefit from further refinement, therefore many mBDRC clusters were unassigned by FR-Match and some others were unassigned by CellHint. Overall, further investigation is needed to disambiguate the 1-to-many matches and resolve the disagreed matches and disagreed unassigned. Having said that, the 21 unique 1-to-1 matches can form the initial basis for a consensus meta-atlas of kidney cell types, while work in forming a stable meta-atlas for human kidney is performed.

**Figure 10.**
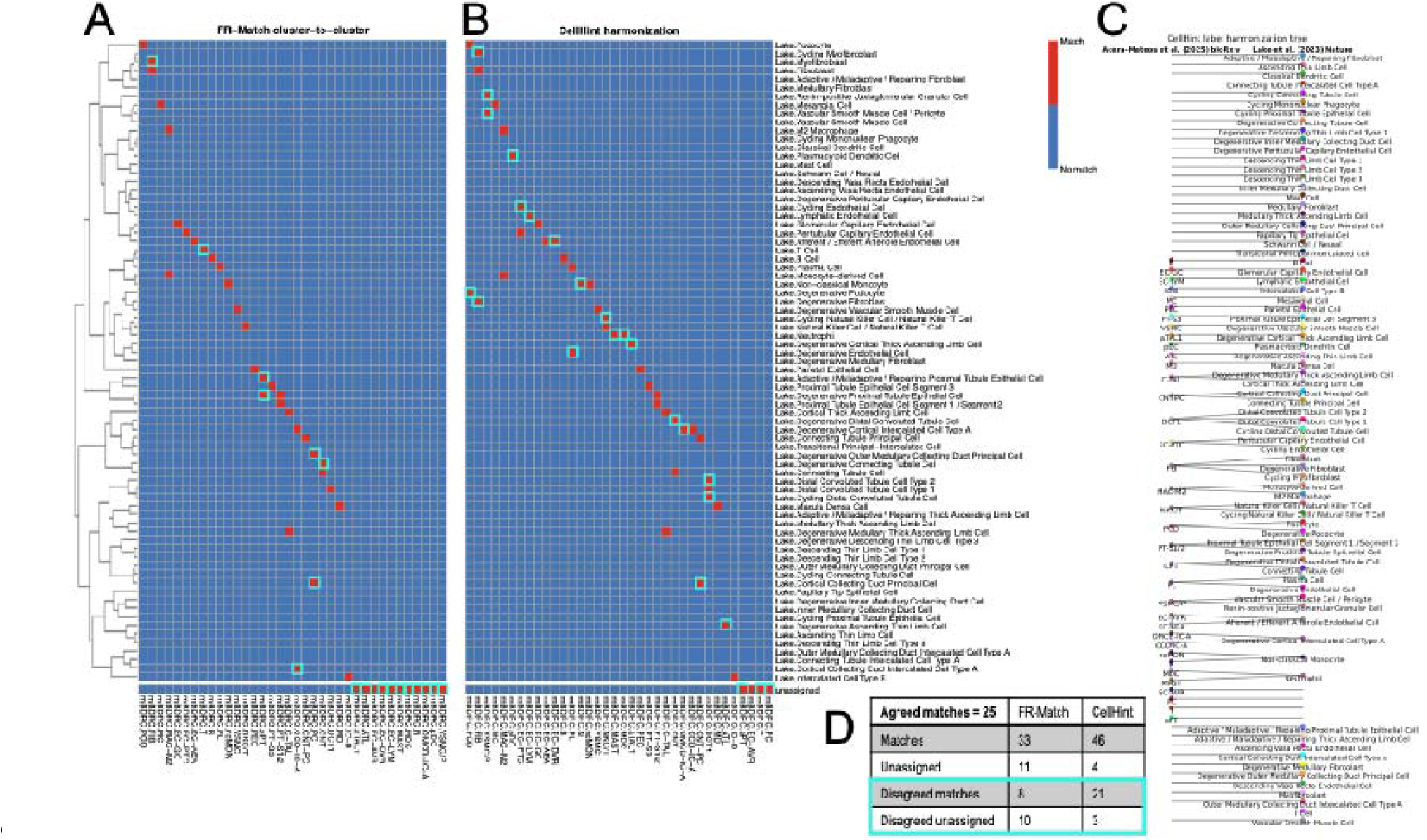
Cluster-based matching of human kidney cell types. **(A)** FR-Match cluster-to-cluster two-way matching results. **(B)** CellHint cell type harmonization results. **(C)** Tree plot outputted from CellHint. Each red block in (B) correspond to an edge in (C). **(D)** Total count table and disagreements between FR-Match and CellHint. Cyan boxed correspond to disagreements.

### Generalizability in more studies

Eight additional community-contributed cell atlases (retina [38, 39], brain [40], breast [41], liver [42], gut [43], non-small cell lung cancer [44], fetal immune system [45], and mouse skeleton [46]) were used to illustrate the generalizability of these benchmarked methods and their performance. Reference and query datasets for each tissue atlas and their summary statistics were reported in **Supplementary Table 8A** [38–50]. The generalizability test evaluates 16 datasets ranging from 9 cell types to 137 cell types, and from ∼24,000 cells/nuclei to more than 1 million cells/nuclei, using a variety of single cell platforms, e.g., multiple versions of 10x, SMART-Seq, Drop-Seq, etc. Wherever possible, all methods were applied, with the only exception for scArches if no count data is available. More details are in **Methods**. Method performances were summarized in **Supplementary Table 8B**. Adjusted Rand Index (ARI) was reported for the cell-based methods, and the number of unique 1-to-1 match and number of unassigned were reported for the cluster-based methods. The higher ARI and the more unique matches are indicative of more concise matching results between the query and reference, corresponding to a cleaner diagonal pattern in the matching plots (**Supplementary Figures 12-19**). While FR-Match has the highest ARI or the most uniquely matched cell types in more than half of the studies, the cluster-based methods, CellHint and FR-Match, are more useful to complement the results from each other, compared to the cell-based methods. Overall, these results showed that the benchmarked methods are generalizable in a range of studies and organ systems, though occasionally a couple methods failed. While some methods performed more “ideally” (since no ground truth) than some other methods, it emerges that a selective set of methods that perform relatively better than others should be used to establish the consensus in meta-atlases.

## Discussion

In this study, cell type matching performance of seven methods – Azimuth, CellTypist, CellHint, FR-Match, scArches, scPred, and singleR – were benchmarked using well-annotated human lung and kidney atlas datasets. Generalizability of selective methods were demonstrated in a variety of organ systems. The results highlight the strengths and limitations of current cell type matching approaches when applied to large-scale atlas datasets. (1) While almost all methods demonstrated high accuracy in the overall results, particularly in well-represented cell types, significant variation was observed in their ability to accurately predict rare cell types.

Discrepancy between the overall performance and cluster-level performance suggests that many of the current approaches are biased towards larger and more abundant cell types. (2) The impact of imbalance in cluster sizes among the cell types is clearly elaborated in this study, as well as Maan et al. (2023) [51]. While many of the current tools provide convenient cell-based predictions, more evidence has emerged that suggests the cell-based approaches may produce misleading results if cluster-level variation is not adequately taken into account. (3) One common limitation to many of the methods is the lack of mechanism to identify novel cell types. With the goal of completing a comprehensive atlas that describes all healthy cell types in the human body, the ability to recognize novel cell types not identified in the initial atlas will be essential as new datasets are generated. In particular, the pre-trained models seem to insufficiently address weakly matched cells and force them to be matched in their algorithms. These limitations highlight the importance of future efforts to further refine the label transfer-based algorithms to better capture rare and novel cell types.

Among the limitations, the impact of size imbalance in scRNA-seq data has been well-studied in Maan et al. (2023) [51], where 2,600 integration experiments were conducted, including perturbation experiments with balanced dataset (“control”), random downsampling (“downsampled”), and complete removal of cell type compartment (“ablated”). These experiments indicates that “sample imbalance has substantial impacts on downstream analyses and the biological interpretation of integration results”, Results presented in this study (e.g., **Figure 2C**) further confirmed some of the observations from the Maan et al. study. The most significant impact of size imbalance in our study is that larger clusters tend to have better cell type matching accuracy than smaller clusters by many computational methods. One possible reason is that the reduced dimensional space of these methods usually consists of highly variable genes that may be dominated by larger clusters contributing more variance and may miss some feature genes representing smaller clusters. One of the methods that had high accuracy in smaller clusters is FR-Match. Instead of highly variable genes, FR-Match uses informative gene space consisting of the supervised NS-Forest feature selection that consists of similar number of feature genes to equally represent each cluster regardless of cluster sizes.

The impact of sample imbalance in the following Fridman-Raftsky test (FR test) had also been explicitly investigated in our earlier study (Supplementary Figure S30-S31 in [19]), therefore, iterative subsampling scheme had been implemented in FR-Match to account for the impact of the sample imbalance. More details are explained in **Methods**.

Earlier in this project, some attempts were made to manually match the CellRef annotated cell types with their HLCA counterparts, and vice versa. Even with domain knowledge, these attempts failed to find one-to-one matches for many cell types. This issue is not unique to lung. In fact, this portion of the study reveals a common struggle in current cell type annotation efforts. Alternative methods based on nature language processing (NLP) and generative artificial intelligence (AI) model, such as OnClass [52] and scGPT [53], have also been proposed to perform cell type annotation for single cell data. However, their performances have yet to be benchmarked.

To circumvent these issues, this study assumes that high-quality predictions are more likely when there is greater consensus among the methods. After obtaining results from the benchmarked cell-based and cluster-based methods, a majority voting strategy was adopted to recommend a consensus matching result. The resulting cross-comparison table deciphers commonalities and complementary results that the constituent methods produce, leading to an evidence-based meta-atlas. Moreover, further exploration into how cell type definitions and marker genes could be standardized across datasets could lead to improved consensus in cell type nomenclature and enhance the construction of cell atlases for the single cell community.

There are several aspects of atlas building that are not directly addressed by this study but are important to consider when evaluating a cell atlas dataset or choosing a computational method. (1) While this study focused on the inter-atlas variability, adequately addressing the intra-atlas variability, coming from data generated by different labs within the same integrated atlas, is key for building an atlas towards community-wide consensus cell types. The intra-atlas variability of cell type annotations can come from multiple sources, for example, different sequencing depth and clustering resolutions that would lead to different coverage of the granularity of cell types. In the HLCA study, the authors have shown that there is significant inconsistency of the intra-atlas annotations across its constituent datasets generated by different labs (Figure 3e in [16]). To validate the approach that the atlas builders have taken to address the intra-atlas variability, the meta-analysis presented in this study could complementarily confirm that the re-annotated cell types in the integrated atlas are robust, suggesting effective removal of the intra-atlas variability in the well-curated cell atlases. (2) Concerns about low-quality or strongly batch-affected data that may degrade the reference atlas is not fully explored. This study focused on well-curated “cell atlas” where stringent quality control, including filtering low-quality cells and removing batch effects, have been applied when constructing the atlas. The selected atlas should be associated with a peer-reviewed publication, and the quality of the identified cell types is widely accepted by the community. That being said, each new dataset should also be evaluated for quality or strong batch effects before adding to the meta-atlas using community-approved best practices. There might be edge cases, e.g., chondrocyte in CellRef with only 6 cells and very low cluster quality metric (silhouette score = −0.936), that needs to be addressed in a case-by-case basis.

Looking forward, it would be expected that each new dataset is an enhancement to the precedent ones, particularly when multimodal assay such as ATAC-sequencing and spatial technologies such as Xenium, CosMx, MERFISH and others are made possible at the single cell and subcellular levels, which will provide additional information to help determine the cell type identities and resolve edge cases. (3) Work presented in this benchmarking study focused on single cell data for stable cell types and cell atlases for adult normal tissues. There is an equally important branch of single cell genomics research for transient cell states and developmental cell atlases, which was not covered in this study. Computational methods benchmarked for transient cell states analysis, i.e., the single cell trajectory inference, can be found in Saelens et al. (2019) [54].

Other than the matching performance, computational cost for this type of analysis is high on atlas datasets. Both lung and kidney analyses presented in this study were done on High Performance Computer (HPC) clusters, which are only “modest” sizes of atlases with ∼0.5M cells or fewer. In our experience, we found that the cluster-based methods are faster than cell-based methods and using an organ-specific pre-trained model is significantly faster than without using an organ-specific pre-trained model, such as scPred and singleR. Since these models are still quite complex, even with some steps pre-trained for the users, the pipeline for each method would still need several customizations (e.g., converting data into required format, e.g., anndata, seurat or SingleCellExperiment object, taking care of feature names, e.g., ensemble gene identifiers or gene symbols, using raw counts as input data or preprocessing needed, and optimizing methodology in terms of number of epochs or iterations, etc.), making the method computational time not directly comparable. Therefore, the end-to-end computational time comparison is not included in this study.

As atlas size quickly approaching multimillion cells, computational cost and algorithm scalability become a major bottleneck. It is one of the reasons why pre-trained models exist, so that only the pre-trained latent space of the reference dataset with reduced dimensionality was saved in the model object to be used directly by the downstream users. For example, the Azimuth reference seurat object only contains highly variable genes instead of all genes. Similarly, as stated in the scArches documentation, “As scArches allows the mapping of query data onto an existing reference embedding, we will only need to download the embedding of the HLCA reference. That saves a lot of time and memory compared to downloading the full count matrix.” Following the cell-based integration strategy like these, it will be more and more challenging for the users to train these models with the rapidly increasing size of the reference datasets, especially when new and old datasets are continuously integrated in a cell-wise growing atlas. The proposed strategy in this study is intended to mitigate such challenge by introducing the concept of incremental growth of cell types. It proposes to build a complete cell atlas in an evolving context by representing the current cell types in a cell type knowledgebase and incrementally populate new cell types to the knowledgebase using validated computational methods benchmarked in this study and others. Based on the above benchmarking results, this study encourages the cluster-level matching in support of the incremental cell type knowledge growth strategy, in this way, the computational need for growing the knowledgebase is more manageable at an incremental pace.

Last, but not least, this study sheds light on how to reliably grow the cell type knowledge in cell atlases when new datasets are ingested into the data corpus. Currently, the common approach is to integrate new datasets into an existing atlas by re-running the whole integration pipeline, including batch correction, latent space training, re-clustering, and expert annotation, etc. This type of approach will likely produce slightly different cell cluster membership every time the atlas is updated, leading to potentially different cell type annotations of the same cell in different versions. In this way, future studies will be hard to reproduce results or leverage findings that are derived using existing and earlier versions of the atlas. A more stable reference would be obtained if using a mechanism that allows organic growth of cell types in an incremental fashion by adding new cell types when there is sufficient evidence from new datasets while preserving the cell cluster membership used in the original atlas. This would require the existing and new methods to have more enhanced and consistent performance for cluster-level matching and the ability to identify novel cell types, distinguishing exact matches from similar cell types.

### Conclusion

The benchmarking analysis presented in this study highlights the challenges of identifying rare and novel cell types using computational cell type matching methods. A meta-atlas for healthy adult human lung with 68 distinct cell types from HLCA and CellRef was formed based on the meta-analysis using benchmarked methods. Additional analysis from a variety of organ systems confirmed our findings. This computational framework supports an incremental knowledge growth strategy for expanding cell types in an evolvable cell meta-atlas.

## Methods

### Glossary

Definitions of the following terms are provided in the context of this manuscript.

**Label transfer:** a computational method that assigns cell type labels or cell type annotations from a well-annotated reference dataset to a new, unannotated dataset. This type of method transfers labels or annotations to *individual cells*.

**Cell atlas:** a comprehensive map that aims to describe all the cell types in an organism, tissue, or biological system, along with their characteristics and relationships.

**Cell atlas dataset:** a well-annotated/curated, high-quality/high-resolution single cell (e.g., scRNA-seq) dataset with consistently processed data (i.e. with best practice workflow), expert-annotated cell types, and standardized metadata. (Short definition: the dataset of a cell atlas). It can be used as a reference for mapping new dataset or other comparative analysis.

**Cell cluster:** a computationally determined group of cells that have similar transcriptional profiles, identified through clustering algorithms applied to e.g., scRNA-seq data. (Short definition: a cluster in a single cell dataset).

**Cell meta-atlas:** an aggregated, standardized (e.g., linking to Cell Ontology), and harmonized collection of cell types from cell atlas datasets that form a unified reference of cell types. (Short definition: “meta-analysis” of cell atlases).

**Cell phenotype:** the manifestation of a cell’s identity and function determined by its expressed genes/proteins/metabolites.

**Cell type:** a group of cells that share a stable, characteristic transcriptional profile that is distinct from other groups of cells. In this manuscript, we often refer the annotated cell clusters in a cell atlas dataset as cell types.

**Cell type matching:** a process or computational method that identifies which cell types (i.e., annotated cell clusters) in one dataset correspond to cell types in another dataset. When it indicates a computational method, this type of method aligns *clusters* across datasets.

**Incremental cell type knowledge growth:** a framework to expand cell types in an evolvable cell meta-atlas to form a cell type knowledgebase by incrementally adding novel cell types from new datasets to the existing cell type collection.

**HRA:** a comprehensive, high-resolution, three-dimensional atlas of major cells in the healthy human body [6].

### Healthy adult lung and kidney datasets

The Human Lung Cell Atlas (HLCA) is an integrated cell atlas consisting of 2.4 million cells from 486 individuals spanning 49 datasets of the human respiratory system in health and disease [16]. A filtered subset of healthy cells forms the HLCA core dataset, which contains more than 0.5 million cells. Hereinafter, we will use “HLCA dataset” to mean the HLCA core healthy dataset. The HLCA atlas was constructed by adopting a supervised data integration strategy that builds a five-level hierarchical cell identity reference framework. At the finest level, the HLCA core defines 61 annotated cell types (“ann_finest_level”) with well-distributed metadata of donors’ demographics, anatomical, and technical variables, suggesting that any batch effects have been effectively normalized. Note that the HLCA reference includes nasal brushing samples in addition to lung structures.

The LungMAP single-cell reference (CellRef) is a reference cell atlas for both normal human and mouse lungs developed using a guided approach for atlas construction [15], where a pre-defined cell type dictionary (i.e., LungMAP CellCards - https://www.lungmap.net/research/cell-cards) was used to guide the annotation of cell identities. The CellRef human dataset contains more than 0.3 million cells/nuclei consolidated from 10 datasets and 104 donors using a computational pipeline designed to account for both biological and technical variations. Hereinafter, we will use “CellRef dataset” to mean the CellRef human dataset. The atlas defines 48 cell types (“celltype_level3_fullname”) in normal human lung.

The Human Kidney Atlas 1.0 (HKA) is “a multimodal single-cell and spatial atlas with integrated transcriptomic, epigenomic and imaging data over three major consortia: the Human Biomolecular Atlas Program (HuBMAP), the Kidney Precision Medicine Project (KPMP) and the Human Cell Atlas (HCA)” [17]. The integrated single-nucleus and single-cell RNA-seq data consists of more than 400,000 high-quality nuclei/cells from normal tissues (18 living donor biopsy cores) and diseased tissues (12 acute kidney injury and 15 chronic kidney disease). The integrated atlas defines 75 consensus cell types (“subclass.full”) across conditions. Only the normal cells/nuclei (107,701) were used in this study.

The multimodal benchmarking dataset for renal cortex (mBDRC) is generated by the CZI Seed Network. From a collection of kidney tissues from 19 “healthy” donors, 36 cell types (“author_cell_type”) were reported by integrating 3’ and 5’ scRNA-seq and multiomics (snRNA-seq and snATAC-seq) joint profiling. “Fresh normal tissue from the unaffected part of surgically removed kidney of patients undergoing total nephrectomy” were used to construct this reference [18]. This dataset is currently available in CELLxGENE as a preprint version.

### Algorithms

Table 2 summarizes the cell-based algorithms benchmarked in this study along with notable parameters and features specified in this study. “Model specification” indicates the name of the pre-trained models used in Azimuth, CellTypist, and scArches, the informative gene space used in FR-Match, the prediction model used in scPred, and the fine-tuning option in singleR. “Reference” specifies the reference dataset that the pre-trained models were trained on. “Code env” states the code environment that the method is available in. “Confidence score” describes the process used to assign a confidence score to each prediction. “Preprocessing” details the notable pre-processing steps taken to prepare the query data.

**Table 2:**
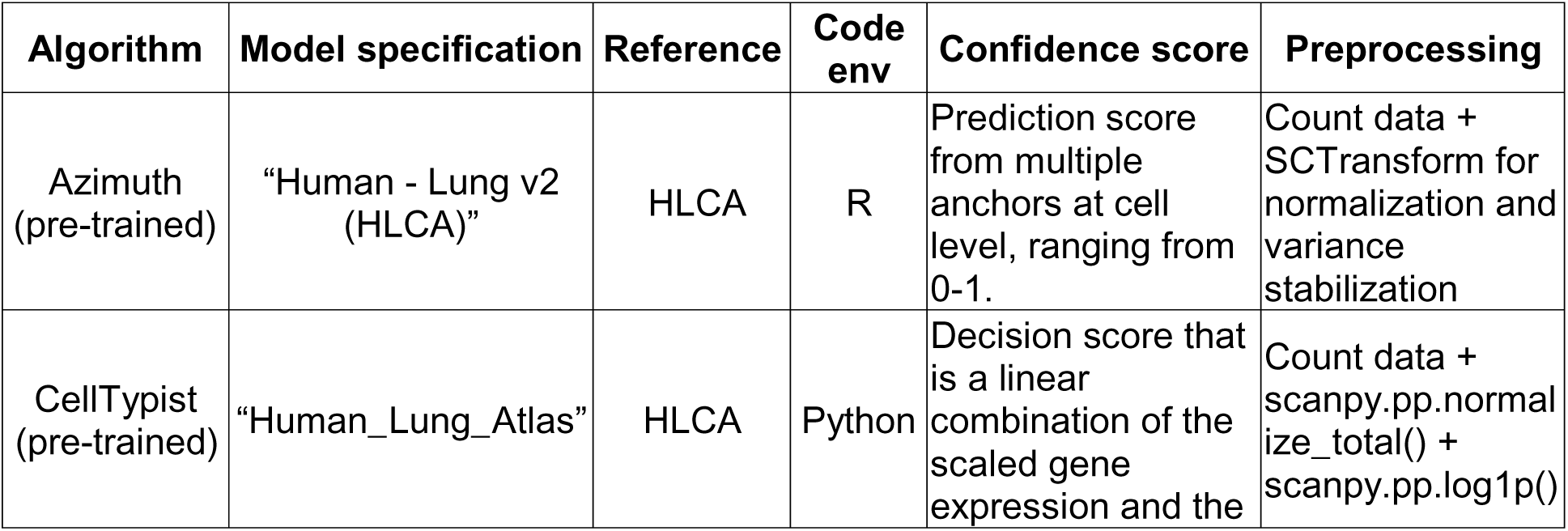
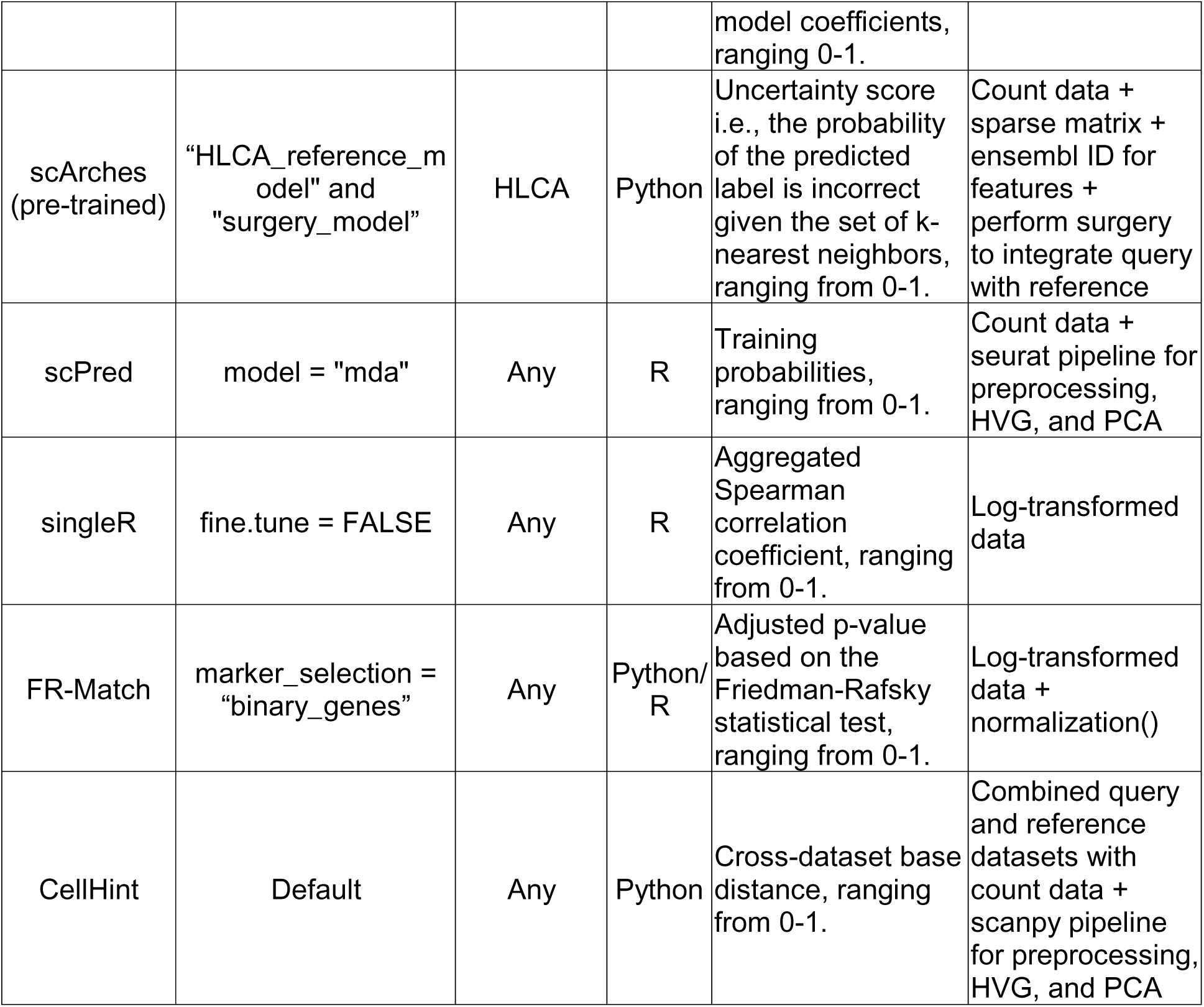
Summary of algorithms used in this study.

### Azimuth

Azimuth employs anchors in lower-dimensional space to transfer cell-type labels from reference to query data clusters. This method first projects the query data onto a lower-dimensional space (UMAP) by applying sparse principal component analysis (sPCA) to the reference dataset.

Subsequently, each query cell is compared to reference-based anchor cells to construct a k-nearest neighbor graph. Finally, Azimuth utilizes weighted voting to classify each query cell to a corresponding reference cluster.

Though an automated web application is available for Azimuth cell annotation, for large-scale query datasets and the purposes of this study, Azimuth was executed locally in R using the “RunAzimuth()” function from the Azimuth library. The pre-trained reference model is readily downloadable from the Azimuth website through Zenodo (https://zenodo.org/records/6342228).

“RunAzimuth()” accepts several forms of input for query data, including path names to AnnData H5AD files and Seurat objects. To perform the HLCA cross-validation, the query data, was subset into 10 folds according to pre-split indices. The HLCA dataset is available as a Seurat object; and each subset was used as input into “RunAzimuth()”. To perform the cross-dataset matching, the CellRef dataset required additional steps to be taken before running Azimuth. As the CellRef dataset is available as a H5AD file, the file path was provided as input for “RunAzimuth()”. Internally, the algorithm uses Seurat’s “LoadH5AD()” function to convert the H5AD file into a Seurat object for annotation. However, due to some differences in file formatting, both the function and H5AD object were manually modified to successfully convert the data into a Seurat object. A more comprehensive tutorial detailing this process is available in the GitHub repository. Once the H5AD file was converted to a Seurat object, the same procedure was applied to subset the query data into 10 folds for annotation.

### CellTypist

CellTypist utilizes a logistic regression framework with stochastic gradient descent (SGD) optimization to train its reference models. To annotate each cell, CellTypist scales each gene according to parameters in the reference model and calculates decision scores for each query cell based on the linear combination of its scaled gene expression and the model coefficients. It assigns the cell type with the largest score to the query cell. CellTypist also provides an additional refinement step, majority voting, that was enabled in this study. Majority voting uses Leiden clustering to over-cluster query cells and re-assigns each subcluster to the majority predicted label for that cluster.

Same as in the Azimuth process, the query data was first subset into 10 folds using the same fold indices. Each subset was then preprocessed using “sc.pp.normalize_total(query_adata, target_sum = 1e4)” and “sc.pp.log1p(query_adata)” from Scanpy. Then, each fold was annotated using the annotate method from CellTypist with “majority_voting=True”.

### CellHint

CellHint is a two-step process to automatically harmonize cell types across datasets: (1) calculating the global distances between cells from combined query and reference datasets, and (2) building a clustering tree based on the distances using a predictive clustering tree (PCT)-based tool. CellHint takes in raw count data in .h5ad format. After combining the query and reference data into one single data object, standard scanpy pipeline is applied with batch-specific highly variable gene (HVG) selection (i.e., “batch_key = ‘dataset’”). Default setting was used for “cellhint.harmonize()”.

### scArches

ScArches uses architectural surgery to incorporate new input nodes from query data with trainable weights into a pre-trained neural network reference model. The method first maps query data to the pre-trained neural network reference model and trains new input nodes from the query data. Then, it uses a weighted k-nearest-neighbor classifier trained on the latent space representation of the reference dataset. Lastly, it calculates the uncertainty of each of the k labels for each query cell and classifies the query cell with the least uncertain label.

To use scArches, the query data was first checked for whether (1) it is a sparse matrix, (2) whether the matrix contains raw counts, and (3) whether the data uses the same features as the input features for its reference model. If the criterion were met, the query data was subset into 10 folds in the same process as Azimuth and CellTypist. Each subset was trained using “surgery_model.train(accelerator=’cpu’, devices=1, max_epochs=surgery_epochs, **early_stopping_kwargs_surgery)” to incorporate the subset into the reference model. The specific parameters used for surgery model training can be found in the tutorial notebook available in the GitHub repository. After surgery, the new surgery model is used to create a latent representation of the reference model to be used in label transfer.

### scPred

ScPred is a standalone R package that was built based on the caret R package of prediction models to classify cells based on a low-dimensional representation of gene expression (e.g. PCA). The low-dimensional representation is obtained from standard seurat workflow. When training the model on the HLCA reference, “mda” was chosen instead of the default “svmRadial” due to a documented issue on GitHub (https://github.com/powellgenomicslab/scPred/issues/17). Instead of the ten-fold cross-validation for HLCA, scPred has its internal reference training and evaluation steps that could assess the training performance on the reference. After training the HLCA reference, “get_probabilities()” was used to obtain a training probabilities matrix based on the training performance. The confidence score for each HLCA cell is the highest training probability and the prediction is the cell type corresponding to the highest probability.

### singleR

SingleR is a standalone R package that annotates cell identities based on correlating gene expression of pure cell types (in a reference dataset or microarray database) with single-cell gene expression. The annotation is performed by calculating the Spearman correlation for each single cell independently across all pure cell types, then through a fine-tuning step to improve accuracy. The score matrix is the correlation-based scores for each cell against every possible reference cell type, prior to any post-hoc fine-tuning. Therefore, for the benchmarking purpose in this study, fine-tuning was tuned off (“fine.tune = FALSE”) in order to be able to work with the scores truly reflecting the correlation strength.

### FR-Match

Originally developed in R, FR-Match is now implemented in Python 3 (https://github.com/BeverlyPeng/frmatch) to streamline the matching workflow in the same code environment as Scanpy’s AnnData object [55] and the NS-Forest algorithm [27, 28] for reduced gene space selection. AnnData is a Python package that manages the annotated single cell experiment data with computational efficient features. NS-Forest is a Python package for machine learning-based selection of cell type marker genes, which is compatible with the AnnData object and the Scanpy workflow.

The inputs to FR-Match are two AnnData objects (one for query and one for reference) and user’s choice of “marker_selection”. The choices for “marker_selection” can be “NSForest_markers”, “binary_genes”, or a user-provided list of marker genes. The reference AnnData object has the NS-Forest results saved in the unstructured metadata slot (.uns). If “NSForest_markers” or “binary_genes” is chosen, the corresponding list of genes will be extracted directly from .uns. Both the query and reference objects will be reduced to the chosen list of genes as the dimensionality reduction step in FR-Match. In this study, “marker_selection=‘binary_genes’” was used.

Before matching, the “normalization()” function is used to align the expression distributions of the two datasets, which performs a weighted per-gene min-max scaling to normalize the gene expression values to the range of [0, 1]. There are two main functions for conducting the matching: FRmatch() for cluster-based matching, and FRmatch_cell2cluster() for cell-based matching. As the other methods, the cell-based matching was conducted through subsetting the query data into 10 folds and matching each fold to the reference respectively. The cluster-based FR-Match does not need to subset the query data as it leverages the query clusters and assigns matching to each of the query clusters as a whole. One of the outputs from FR-Match algorithm is the adjusted p-values based on the non-parametric Friedman-Rafsky statistical test, which serves as an objective confidence score for the matching quality. Based on the adjusted p-value, a matching assignment is determined by adjusted p-value >= 0.05, and “unassigned” is determined by adjusted p-value < 0.05.

For the two-way matching, FRmatch() was run twice with the query and reference datasets switched. The plot_bi_FRmatch() function was used to combine the two sets of results and produce the two-way matching plot. Note that, since FR-Match utilizes the query cluster information, a query cluster with too few cells are removed in the algorithm. Hence, the query lymphatic EC proliferating in HLCA and the query chondrocyte in CellRef were removed in the FR-Match analyses.

### Iterative subsampling scheme in FR-Match

FR-Match is run on each pair of query and reference clusters with equal number of subsampled cells (usually 20 cells) and iteratively sampling cells across the entire cluster with certain number of iterations (usually 1000 iterations). These parameters are tunable using “subsamp_size” and “subsamp_iter”. For cluster-to-cluster matching, it is not necessary to sample every single cell in the cluster, assuming similar cells are grouped in the same cluster. For cell-to-cluster matching, it takes more iterations to sample the entire query cluster for larger clusters and fewer iterations for rare types, therefore, FR-Match could achieve higher accuracy in the rare types and had slightly lower accuracy in the very large clusters that would require much more iterations to go through the entire cluster. To account for the outlier cluster sizes, the parameters “subsamp_iter_custom_k” is implemented to set customized number of iterations for the very large clusters that are not at the same scale as the remaining clusters. E.g., if “subsamp_iter_custom_k = 5”, it will set the number of iterations to be at least “subsamp_iter” or 5 times the cluster size for each query cluster.

### Cell type taxonomy dendrogram

Dendrogram is an informative visualization of a cell type taxonomy. The HLCA taxonomy dendrogram is generated using “scanpy.tl.dendrogram()” [56], which computes a hierarchical clustering for the given cell type annotations. Hierarchical clustering is an unsupervised machine learning technique that groups cell types based on the similarity of their expression profiles in a tree structure, so that similar cell types are grouped together in the same branch. In the HLCA dendrogram, the cell types form major branches of immune, epithelial, endothelial and stromal cells, reflecting the hierarchy of cell classes and types.

### Colorize ROC curves and confidence scores

The colorized ROC curves (**Supplementary Figure 2**), shaded by the confidence score, help visualize the relationship between the prediction confidence with the true and false positive rates. In an ideal case, the colorized plot would display a 1 to 0 score gradient as the curve transitions from the start point (0, 0) to the end point (1,1), which would indicate high confidence associated with correct (true positive) predictions (red-orange portion of the curve) and medium to low confidence associated with incorrect (false positive) predictions (cyan-blue portion of the curve). The heatmaps (**Supplementary Figure 3**) further illustrate the effect of false positives, i.e., cells from specific clusters that were mislabeled with high confidence (the off-diagonal values in the heatmap), on the shape of the ROC curves and quality of the predictions by each method. Note that clusters in the heatmap are ordered by their similarities to each other based on the HLCA taxonomy dendrogram. For example, FR-Match, which has the highest AUC score, has a nearly clean diagonal in its heatmap. There are a few dark squares just off the diagonal, which means that high confidence scores were assigned to closely similar cell types. SingleR, Azimuth, CellTypist, and scArches have noisier heatmaps, resulting in the shallowing of the slopes of their ROC curves. The scPred heatmap has the most dark squares (confidence score close to 1), which led to the unusual shape of its ROC curve, suggesting the confidence scores do not show a linearly monotonic relationship with true positives predictions. The confidence score heatmaps suggest a need to eliminate false positives (i.e., off-diagonal high confidence scores) in the score calculation.

### NS-Forest marker gene selection

NS-Forest is a marker gene selection algorithm based on random forest machine learning. The NS-Forest v4 algorithm is used in this study. The NS-Forest algorithm reports the F-beta score, precision, and recall as metrics to quantify the strength of cell type classification using the selected marker genes. NS-Forest marker genes are the minimum combination among the top features ranked by random forest Gini Impurity index and binary scores that achieves the highest F-beta score in the decision tree evaluation. The algorithm also outputs the top 10 binary genes double-ranked by random forest and binary scores, which represents a more comprehensive list of genes that are highly expressed in a given cell type. The NS-Forest marker genes were visualized in the “barcode” plots; and the binary genes were used in FR-Match in the lung analysis. The NS-Forest “barcode” plots for all recommended matching pairs and atlas-specific cell types are in **Supplementary Figure 20-24**. The recommended matching results with NS-Forest marker genes are in **Supplementary Table 2**.

### High confidence match

For the cell-based methods, the matching results were saved in a table (**Supplementary Table 1**) where rows are the CellRef cells, and columns are the true CellRef label (“cellref_true”) and the predicted HLCA label and corresponding prediction score by each method (“azimuth_pred”, “azimuth_score”, “celltypist_pred”, “celltypist_score”, “frmatch_pred”, “frmatch_score”, “scarches_pred”, “scarches_score”). To summarize the cell-based matching results into cluster-level matching, a contingency table of matching proportions p_ij_ for predicted HLCA label = i and true CellRef label = j were calculated, where

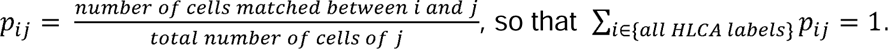

For each method, these values were plotted in **Figure 5A**, where the column sums should be1; however, the row sums of these proportions may be larger than 1, which means that, at the cluster-level, many CellRef clusters were matched to the same HLCA cluster. To normalize the row sums, we define 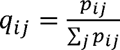, which can be interpreted as the cluster-level precision, ranging from 0 to 1. In the most ideal situation where the CellRef cluster and the HLCA cluster is a one-to-one match, p_ij_ = q_ij_ = 1. However, this is very hard to achieve. Practically, the high confidence matches are determined by requiring p_ij_ > 0.5 and q_ij_ > 0.3, which means the majority of the CellRef cells of label j were matched to HLCA label i, and the cluster-level precision is at least 30%. The values for p_ij_ and q_ij_ for the recommended matches are in **Supplementary Table 3**; and all high confidence matches for each method are in Supplementary Table 4-7.

### Analyses of more studies

The query and reference datasets identified in Supplementary Table 8 were processed separately. Count data were used for scArches; log-transformed data were used for the other methods. When processing the retina and fetal immune datasets, significant overlapping cells were noted and removed from the query data to avoid the “easy” situation that the same cells were matched in the query and reference datasets. Integrated cell type annotations were provided by the brain atlas, but the integrated dataset was not available. Two sub-studies from the brain atlas were matched on their integrated label instead of their original annotations.

Similarly, two sub-studies from the mouse skeleton atlas were matched on their integrated label. After data processing and identifying the annotation levels to match, the same script templates of each method were used for all studies.

## Declarations

### Ethics approval and consent to participate

Not applicable.

## Consent for publication

Not applicable.

## Data availability

The HLCA dataset was download from CZ CELLxGENE: https://cellxgene.cziscience.com/collections/6f6d381a-7701-4781-935c-db10d30de293, namely “An integrated cell atlas of the human lung in health and disease (core)”.

The CellRef dataset was downloaded from LungMAP: https://www.lungmap.net/omics/?experiment_id=LMEX0000004396, namely “LungMAP_HumanLung_CellRef.v1.1.h5ad”.

The Lake et al. dataset was download from CZ CELLxGENE: https://cellxgene.cziscience.com/collections/bcb61471-2a44-4d00-a0af-ff085512674c, namely “Integrated Single-nucleus and Single-cell RNA-seq of the Adult Human Kidney”.

The Acera-Mateos et al. (mBDRC) dataset was download from CZ CELLxGENE: https://cellxgene.cziscience.com/collections/4cbb929b-b03b-4aa8-a943-00f61dc22641, namely “multimodal benchmarking dataset for renal cortex characterization”.

Data download sites of the additional studies are in Supplementary Table 8.

## Code availability

Codes developed in this study are documented in the tutorial notebooks, as a benchmark and protocols to compare and optimize current and future cell type matching tools, available via GitHub (https://github.com/jjoycehu/manuscript).

Azimuth in R: https://github.com/satijalab/azimuth

CellTypist in Python: https://github.com/Teichlab/celltypist

scArches in Python: https://github.com/theislab/scarches

FR-Match in R: https://github.com/JCVenterInstitute/FRmatch

FR-Match in Python: https://github.com/BeverlyPeng/frmatch

CellHint in Python: https://github.com/Teichlab/cellhint

## Authors’ contributions

YZ, RHS and KB conceived the project. YZ and RHS designed the study. JH and BP implemented the workflows and conducted the analyses. JH, BP, AVP, BX, and VAD contributed to the data analyses. JH, RHS and YZ interpreted the results. JH and YZ managed the datasets and results. BP built the software package. AB, BWH, and KB coordinate cell type annotation usage in the Human Reference Atlas resource. YZ and CD contributed to the project management. JH, BP, KB, RHS and YZ wrote the manuscript. All authors agreed on the contents of the manuscript.

## Acknowledgements

Not applicable.

## References

1. Abdulla S, Aevermann B, Assis P, Badajoz S, Bell SM, Bezzi E, Cakir B, Chaffer J, Chambers S, Cherry JM, Chi T, Chien J, Dorman L, Garcia-Nieto P, Gloria N, Hastie M, Hegeman D, Hilton J, Huang T, Infeld A, Istrate AM, Jelic I, Katsuya K, Kim YJ, Liang K, Lin M, Lombardo M, Marshall B, Martin B, McDade F, Megill C, Patel N, Predeus A, Raymor B, Robatmili B, Rogers D, Rutherford E, Sadgat D, Shin A, Small C, Smith T, Sridharan P, Tarashansky A, Tavares N, Thomas H, Tolopko A, Urisko M, Yan J, Yeretssian G, Zamanian J, Mani A, Cool J, Carr A. CZ CELLxGENE Discover: a single-cell data platform for scalable exploration, analysis and modeling of aggregated data. Nucleic Acids Res. 2025;53(D1):D886–d900. Epub 2024/11/28. doi: 10.1093/nar/gkae1142. PubMed PMID: 39607691; PMCID: PMC11701654.

2. Jain S, Pei L, Spraggins JM, Angelo M, Carson JP, Gehlenborg N, Ginty F, Gonçalves JP, Hagood JS, Hickey JW, Kelleher NL, Laurent LC, Lin S, Lin Y, Liu H, Naba A, Nakayasu ES, Qian WJ, Radtke A, Robson P, Stockwell BR, Van de Plas R, Vlachos IS, Zhou M, Börner K, Snyder MP. Advances and prospects for the Human BioMolecular Atlas Program (HuBMAP). Nat Cell Biol. 2023;25(8):1089–100. Epub 2023/07/20. doi: 10.1038/s41556-023-01194-w. PubMed PMID: 37468756; PMCID: PMC10681365.

3. Tarhan L, Bistline J, Chang J, Galloway B, Hanna E, Weitz E. Single Cell Portal: an interactive home for single-cell genomics data. bioRxiv. 2023.Epub 2023/07/28. doi: 10.1101/2023.07.13.548886. PubMed PMID: 37502904; PMCID: PMC10370058.

4. Regev A, Teichmann SA, Lander ES, Amit I, Benoist C, Birney E, Bodenmiller B, Campbell P, Carninci P, Clatworthy M, Clevers H, Deplancke B, Dunham I, Eberwine J, Eils R, Enard W, Farmer A, Fugger L, Göttgens B, Hacohen N, Haniffa M, Hemberg M, Kim S, Klenerman P, Kriegstein A, Lein E, Linnarsson S, Lundberg E, Lundeberg J, Majumder P, Marioni JC, Merad M, Mhlanga M, Nawijn M, Netea M, Nolan G, Pe’er D, Phillipakis A, Ponting CP, Quake S, Reik W, Rozenblatt-Rosen O, Sanes J, Satija R, Schumacher TN, Shalek A, Shapiro E, Sharma P, Shin JW, Stegle O, Stratton M, Stubbington MJT, Theis FJ, Uhlen M, van Oudenaarden A, Wagner A, Watt F, Weissman J, Wold B, Xavier R, Yosef N. The Human Cell Atlas. Elife. 2017;6. Epub 2017/12/06. doi: 10.7554/eLife.27041. PubMed PMID: 29206104; PMCID: PMC5762154.

5. Börner K, Teichmann SA, Quardokus EM, Gee JC, Browne K, Osumi-Sutherland D, Herr BW, 2nd, Bueckle A, Paul H, Haniffa M, Jardine L, Bernard A, Ding SL, Miller JA, Lin S, Halushka MK, Boppana A, Longacre TA, Hickey J, Lin Y, Valerius MT, He Y, Pryhuber G, Sun X, Jorgensen M, Radtke AJ, Wasserfall C, Ginty F, Ho J, Sunshine J, Beuschel RT, Brusko M, Lee S, Malhotra R, Jain S, Weber G. Anatomical structures, cell types and biomarkers of the Human Reference Atlas. Nat Cell Biol. 2021;23(11):1117–28. Epub 2021/11/10. doi: 10.1038/s41556-021-00788-6. PubMed PMID: 34750582; PMCID: PMC10079270.

6. Börner K, Blood PD, Silverstein JC, Ruffalo M, Satija R, Teichmann SA, Pryhuber GJ, Misra RS, Purkerson JM, Fan J, Hickey JW, Molla G, Xu C, Zhang Y, Weber GM, Jain Y, Qaurooni D, Kong Y, Bueckle A, Herr BW, 2nd. Human BioMolecular Atlas Program (HuBMAP): 3D Human Reference Atlas construction and usage. Nat Methods. 2025. Epub 2025/03/14. doi: 10.1038/s41592-024-02563-5. PubMed PMID: 40082611.

7. Hemberg M, Marini F, Ghazanfar S, Al Ajami A, Abassi N, Anchang B, Benayoun BA, Cao Y, Chen K, Cuesta-Astroz Y, DeBruine Z, Dendrou CA, De Vlaminck I, Imkeller K, Korsunsky I, Lederer AR, Li JJ, Meysman P, Miller CL, Mullan KA, Ohler U, Panwar P, Patikas N, Schuck J, Siu JHY, Triche TJ, Jr., Tsankov A, van der Laan SW, Yajima M, Yang J, Zanini F, Jelic I. Insights, opportunities, and challenges provided by large cell atlases. Genome Biol. 2025;26(1):358. Epub 2025/10/21. doi: 10.1186/s13059-025-03771-8. PubMed PMID: 41116172; PMCID: PMC12536537 Competing interests: The authors declare no competing interests.

8. Korsunsky I, Millard N, Fan J, Slowikowski K, Zhang F, Wei K, Baglaenko Y, Brenner M, Loh PR, Raychaudhuri S. Fast, sensitive and accurate integration of single-cell data with Harmony. Nat Methods. 2019;16(12):1289–96. Epub 2019/11/20. doi: 10.1038/s41592-019-0619-0. PubMed PMID: 31740819; PMCID: PMC6884693.

9. Polański K, Young MD, Miao Z, Meyer KB, Teichmann SA, Park JE. BBKNN: fast batch alignment of single cell transcriptomes. Bioinformatics. 2020;36(3):964–5. Epub 2019/08/11. doi: 10.1093/bioinformatics/btz625. PubMed PMID: 31400197; PMCID: PMC9883685.

10. Lopez R, Regier J, Cole MB, Jordan MI, Yosef N. Deep generative modeling for single-cell transcriptomics. Nat Methods. 2018;15(12):1053–8. Epub 2018/12/07. doi: 10.1038/s41592-018-0229-2. PubMed PMID: 30504886; PMCID: PMC6289068.

11. Stuart T, Butler A, Hoffman P, Hafemeister C, Papalexi E, Mauck WM, 3rd, Hao Y, Stoeckius M, Smibert P, Satija R. Comprehensive Integration of Single-Cell Data. Cell. 2019;177(7):1888–902.e21. Epub 2019/06/11. doi: 10.1016/j.cell.2019.05.031. PubMed PMID: 31178118; PMCID: PMC6687398.

12. Welch JD, Kozareva V, Ferreira A, Vanderburg C, Martin C, Macosko EZ. Single-Cell Multi-omic Integration Compares and Contrasts Features of Brain Cell Identity. Cell. 2019;177(7):1873–87.e17. Epub 2019/06/11. doi: 10.1016/j.cell.2019.05.006. PubMed PMID: 31178122; PMCID: PMC6716797.

13. Xu C, Lopez R, Mehlman E, Regier J, Jordan MI, Yosef N. Probabilistic harmonization and annotation of single-cell transcriptomics data with deep generative models. Mol Syst Biol. 2021;17(1):e9620. Epub 2021/01/26. doi: 10.15252/msb.20209620. PubMed PMID: 33491336; PMCID: PMC7829634.

14. Luecken MD, Büttner M, Chaichoompu K, Danese A, Interlandi M, Mueller MF, Strobl DC, Zappia L, Dugas M, Colomé-Tatché M, Theis FJ. Benchmarking atlas-level data integration in single-cell genomics. Nat Methods. 2022;19(1):41–50. Epub 2021/12/25. doi: 10.1038/s41592-021-01336-8. PubMed PMID: 34949812; PMCID: PMC8748196 Dermagnostix GmbH and Cellarity. The remaining authors declare no competing interests.

15. Guo M, Morley MP, Jiang C, Wu Y, Li G, Du Y, Zhao S, Wagner A, Cakar AC, Kouril M, Jin K, Gaddis N, Kitzmiller JA, Stewart K, Basil MC, Lin SM, Ying Y, Babu A, Wikenheiser-Brokamp KA, Mun KS, Naren AP, Clair G, Adkins JN, Pryhuber GS, Misra RS, Aronow BJ, Tickle TL, Salomonis N, Sun X, Morrisey EE, Whitsett JA, Xu Y. Guided construction of single cell reference for human and mouse lung. Nat Commun. 2023;14(1):4566. Epub 2023/07/30. doi: 10.1038/s41467-023-40173-5. PubMed PMID: 37516747; PMCID: PMC10387117.

16. Sikkema L, Ramírez-Suástegui C, Strobl DC, Gillett TE, Zappia L, Madissoon E, Markov NS, Zaragosi LE, Ji Y, Ansari M, Arguel MJ, Apperloo L, Banchero M, Bécavin C, Berg M, Chichelnitskiy E, Chung MI, Collin A, Gay ACA, Gote-Schniering J, Hooshiar Kashani B, Inecik K, Jain M, Kapellos TS, Kole TM, Leroy S, Mayr CH, Oliver AJ, von Papen M, Peter L, Taylor CJ, Walzthoeni T, Xu C, Bui LT, De Donno C, Dony L, Faiz A, Guo M, Gutierrez AJ, Heumos L, Huang N, Ibarra IL, Jackson ND, Kadur Lakshminarasimha Murthy P, Lotfollahi M, Tabib T, Talavera-López C, Travaglini KJ, Wilbrey-Clark A, Worlock KB, Yoshida M, van den Berge M, Bossé Y, Desai TJ, Eickelberg O, Kaminski N, Krasnow MA, Lafyatis R, Nikolic MZ, Powell JE, Rajagopal J, Rojas M, Rozenblatt-Rosen O, Seibold MA, Sheppard D, Shepherd DP, Sin DD, Timens W, Tsankov AM, Whitsett J, Xu Y, Banovich NE, Barbry P, Duong TE, Falk CS, Meyer KB, Kropski JA, Pe’er D, Schiller HB, Tata PR, Schultze JL, Teichmann SA, Misharin AV, Nawijn MC, Luecken MD, Theis FJ. An integrated cell atlas of the lung in health and disease. Nat Med. 2023;29(6):1563–77. Epub 2023/06/09. doi: 10.1038/s41591-023-02327-2. PubMed PMID: 37291214; PMCID: PMC10287567.

17. Lake BB, Menon R, Winfree S, Hu Q, Melo Ferreira R, Kalhor K, Barwinska D, Otto EA, Ferkowicz M, Diep D, Plongthongkum N, Knoten A, Urata S, Mariani LH, Naik AS, Eddy S, Zhang B, Wu Y, Salamon D, Williams JC, Wang X, Balderrama KS, Hoover PJ, Murray E, Marshall JL, Noel T, Vijayan A, Hartman A, Chen F, Waikar SS, Rosas SE, Wilson FP, Palevsky PM, Kiryluk K, Sedor JR, Toto RD, Parikh CR, Kim EH, Satija R, Greka A, Macosko EZ, Kharchenko PV, Gaut JP, Hodgin JB, Eadon MT, Dagher PC, El-Achkar TM, Zhang K, Kretzler M, Jain S. An atlas of healthy and injured cell states and niches in the human kidney. Nature. 2023;619(7970):585–94. Epub 2023/07/20. doi: 10.1038/s41586-023-05769-3. PubMed PMID: 37468583; PMCID: PMC10356613 Biomage. A.V. is a consultant for Astute and NxStage. C.R.P. is a member of the advisory board of and owns equity in RenalytixAI, and serves as a consultant for Genfit and Novartis. M.K. has grants from JDRF, Astra-Zeneca, NovoNordisc, Eli Lilly, Gilead, Goldfinch Bio, Janssen, Boehringer-Ingelheim, Moderna, European Union Innovative Medicine Initiative, Chan Zuckerberg Initiative, Certa, Chinook, amfAR, Angion Pharmaceuticals, RenalytixAI, Travere Therapeutics, Regeneron, IONIS Pharmaceuticals, Astellas, Poxel and a patent (PCT/EP2014/073413; ‘Biomarkers and methods for progression prediction for chronic kidney disease’) licensed. F.C. and E.Z.M. are paid consultants for Atlas Bio. F.P.W. receives research support from Astrazeneca, Boeringher-Ingelheim, Vifor Pharma and Whoop. P.M.P. is a consultant for Janssen. S.R. has research funding from AstraZeneca and Bayer Healthcare. S.S.W. is a consultant for GSK, GEHC, JNJ, Strataca, Roth Capital Partners, Venbio, and an expert witness on litigation for Davita and Pfizer. J.R.S. consults for Maze and Goldfinch and receives royalties from Sanfi Genzyme. K.Z. is a co-founder, equity holder and serves on the scientific advisory board of Singlera Genomics. A.S.N. is on the external advisory board for CareDX. L.H.M. is a consultant for Reata Pharmaceuticals, Travere Therapeutics and Calliditas. S.J. is a paid Blue SKy mentor for Meharry Medical College, Nashville and receives royalties from Elsevier. J.L.M. is an employee and shareholder of Solid Biosciences. The other authors declare no competing interests.

18. Acera-Mateos M, Adiconis X, Li JK, Marchese D, Caratù G, Hon CC, Tiwari P, Kojima M, Vieth B, Murphy MA, Simmons SK, Lefevre T, Claes I, O’Connor CL, Menon R, Otto EA, Ando Y, Vandereyken K, Kretzler M, Bitzer M, Fraenkel E, Voet T, Enard W, Carninci P, Heyn H, Levin JZ, Mereu E. Systematic evaluation of single-cell multimodal data integration for comprehensive human reference atlas. bioRxiv. 2025. Epub 2025/03/17. doi: 10.1101/2025.03.06.637075. PubMed PMID: 40093094; PMCID: PMC11908249.

19. Zhang Y, Aevermann BD, Bakken TE, Miller JA, Hodge RD, Lein ES, Scheuermann RH. FR-Match: robust matching of cell type clusters from single cell RNA sequencing data using the Friedman-Rafsky non-parametric test. Brief Bioinform. 2021;22(4). Epub 2020/11/30. doi: 10.1093/bib/bbaa339. PubMed PMID: 33249453; PMCID: PMC8294536.

20. Zhang Y, Aevermann B, Gala R, Scheuermann RH. Cell type matching in single-cell RNA-sequencing data using FR-Match. Sci Rep. 2022;12(1):9996. Epub 2022/06/16. doi: 10.1038/s41598-022-14192-z. PubMed PMID: 35705694; PMCID: PMC9200772.

21. Xu C, Prete M, Webb S, Jardine L, Stewart BJ, Hoo R, He P, Meyer KB, Teichmann SA. Automatic cell-type harmonization and integration across Human Cell Atlas datasets. Cell. 2023;186(26):5876-91.e20. Epub 2023/12/23. doi: 10.1016/j.cell.2023.11.026. PubMed PMID: 38134877.

22. Hao Y, Hao S, Andersen-Nissen E, Mauck WM, 3rd, Zheng S, Butler A, Lee MJ, Wilk AJ, Darby C, Zager M, Hoffman P, Stoeckius M, Papalexi E, Mimitou EP, Jain J, Srivastava A, Stuart T, Fleming LM, Yeung B, Rogers AJ, McElrath JM, Blish CA, Gottardo R, Smibert P, Satija R. Integrated analysis of multimodal single-cell data. Cell. 2021;184(13):3573–87.e29. Epub 2021/06/02. doi: 10.1016/j.cell.2021.04.048. PubMed PMID: 34062119; PMCID: PMC8238499.

23. Domínguez Conde C, Xu C, Jarvis LB, Rainbow DB, Wells SB, Gomes T, Howlett SK, Suchanek O, Polanski K, King HW, Mamanova L, Huang N, Szabo PA, Richardson L, Bolt L, Fasouli ES, Mahbubani KT, Prete M, Tuck L, Richoz N, Tuong ZK, Campos L, Mousa HS, Needham EJ, Pritchard S, Li T, Elmentaite R, Park J, Rahmani E, Chen D, Menon DK, Bayraktar OA, James LK, Meyer KB, Yosef N, Clatworthy MR, Sims PA, Farber DL, Saeb-Parsy K, Jones JL, Teichmann SA. Cross-tissue immune cell analysis reveals tissue-specific features in humans. Science. 2022;376(6594):eabl5197. Epub 2022/05/14. doi: 10.1126/science.abl5197. PubMed PMID: 35549406; PMCID: PMC7612735.

24. Lotfollahi M, Naghipourfar M, Luecken MD, Khajavi M, Büttner M, Wagenstetter M, Avsec Ž, Gayoso A, Yosef N, Interlandi M, Rybakov S, Misharin AV, Theis FJ. Mapping single-cell data to reference atlases by transfer learning. Nat Biotechnol. 2022;40(1):121–30. Epub 2021/09/01. doi: 10.1038/s41587-021-01001-7. PubMed PMID: 34462589; PMCID: PMC8763644 equity in Celsius Therapeutics and Rheos Medicines. The remaining authors declare no competing interests.

25. Alquicira-Hernandez J, Sathe A, Ji HP, Nguyen Q, Powell JE. scPred: accurate supervised method for cell-type classification from single-cell RNA-seq data. Genome Biol. 2019;20(1):264. Epub 2019/12/13. doi: 10.1186/s13059-019-1862-5. PubMed PMID: 31829268; PMCID: PMC6907144.

26. Aran D, Looney AP, Liu L, Wu E, Fong V, Hsu A, Chak S, Naikawadi RP, Wolters PJ, Abate AR, Butte AJ, Bhattacharya M. Reference-based analysis of lung single-cell sequencing reveals a transitional profibrotic macrophage. Nat Immunol. 2019;20(2):163–72. Epub 2019/01/16. doi: 10.1038/s41590-018-0276-y. PubMed PMID: 30643263; PMCID: PMC6340744.

27. Aevermann B, Zhang Y, Novotny M, Keshk M, Bakken T, Miller J, Hodge R, Lelieveldt B, Lein E, Scheuermann RH. A machine learning method for the discovery of minimum marker gene combinations for cell type identification from single-cell RNA sequencing. Genome Res. 2021;31(10):1767–80. Epub 2021/06/06. doi: 10.1101/gr.275569.121. PubMed PMID: 34088715; PMCID: PMC8494219.

28. Liu A, Peng B, Pankajam Ajith V, Duong TE, Pryhuber G, Scheuermann RH, Zhang Y. Discovery of optimal cell type classification marker genes from single cell RNA sequencing data. BMC Methods. 2024;1(1):15. doi: 10.1186/s44330-024-00015-2.

29. Bard J, Rhee SY, Ashburner M. An ontology for cell types. Genome biology. 2005;6(2):R21.

30. Diehl AD, Meehan TF, Bradford YM, Brush MH, Dahdul WM, Dougall DS, He Y, Osumi-Sutherland D, Ruttenberg A, Sarntivijai S. The Cell Ontology 2016: enhanced content, modularization, and ontology interoperability. Journal of biomedical semantics. 2016;7(1):44.

31. Osumi-Sutherland D, Xu C, Keays M, Levine AP, Kharchenko PV, Regev A, Lein E, Teichmann SA. Cell type ontologies of the Human Cell Atlas. Nature cell biology. 2021;23(11):1129–35.

32. Tan SZK, Puig-Barbe A, Goutte-Gattat D, Eastwood C, Aevermann B, Avola A, Balhoff JP, Bayindir IU, Belfiore J, Caron AR. The Cell Ontology in the age of single-cell omics. arXiv preprint arXiv:250610037. 2025.

33. Martinu T, Todd JL, Gelman AE, Guerra S, Palmer SM. Club Cell Secretory Protein in Lung Disease: Emerging Concepts and Potential Therapeutics. Annu Rev Med. 2023;74:427–41. Epub 2022/12/01. doi: 10.1146/annurev-med-042921-123443. PubMed PMID: 36450281; PMCID: PMC10472444.

34. Thai P, Chen Y, Dolganov G, Wu R. Differential regulation of MUC5AC/Muc5ac and hCLCA-1/mGob-5 expression in airway epithelium. Am J Respir Cell Mol Biol. 2005;33(6):523–30. Epub 2005/09/10. doi: 10.1165/rcmb.2004-0220RC. PubMed PMID: 16151054; PMCID: PMC2715330.

35. Wilkinson MD, Dumontier M, Aalbersberg IJ, Appleton G, Axton M, Baak A, Blomberg N, Boiten JW, da Silva Santos LB, Bourne PE, Bouwman J, Brookes AJ, Clark T, Crosas M, Dillo I, Dumon O, Edmunds S, Evelo CT, Finkers R, Gonzalez-Beltran A, Gray AJ, Groth P, Goble C, Grethe JS, Heringa J, t Hoen PA, Hooft R, Kuhn T, Kok R, Kok J, Lusher SJ, Martone ME, Mons A, Packer AL, Persson B, Rocca-Serra P, Roos M, van Schaik R, Sansone SA, Schultes E, Sengstag T, Slater T, Strawn G, Swertz MA, Thompson M, van der Lei J, van Mulligen E, Velterop J, Waagmeester A, Wittenburg P, Wolstencroft K, Zhao J, Mons B. The FAIR Guiding Principles for scientific data management and stewardship. Sci Data. 2016;3:160018. Epub 2016/03/16. doi: 10.1038/sdata.2016.18. PubMed PMID: 26978244; PMCID: PMC4792175 Honorary Academic Editor and consultant.

36. Tan SZK, Kir H, Aevermann BD, Gillespie T, Harris N, Hawrylycz MJ, Jorstad NL, Lein ES, Matentzoglu N, Miller JA. Brain Data Standards-A method for building data-driven cell-type ontologies. Scientific Data. 2023;10(1):50.

37. Bakken T, Cowell L, Aevermann BD, Novotny M, Hodge R, Miller JA, Lee A, Chang I, McCorrison J, Pulendran B. Cell type discovery and representation in the era of high-content single cell phenotyping. BMC bioinformatics. 2017;18(Suppl 17):559.

38. Swamy VS, Fufa TD, Hufnagel RB, McGaughey DM. Building the mega single-cell transcriptome ocular meta-atlas. Gigascience. 2021;10(10). Epub 2021/10/16. doi: 10.1093/gigascience/giab061. PubMed PMID: 34651173; PMCID: PMC8514335.

39. Swamy VS, Batz ZA, McGaughey DM. PLAE Web App Enables Powerful Searching and Multiple Visualizations Across One Million Unified Single-Cell Ocular Transcriptomes. Transl Vis Sci Technol. 2023;12(9):18. Epub 2023/09/25. doi: 10.1167/tvst.12.9.18. PubMed PMID: 37747415; PMCID: PMC10578359.

40. Chen X, Huang Y, Huang L, Huang Z, Hao ZZ, Xu L, Xu N, Li Z, Mou Y, Ye M, You R, Zhang X, Liu S, Miao Z. A brain cell atlas integrating single-cell transcriptomes across human brain regions. Nat Med. 2024;30(9):2679–91. Epub 2024/08/03. doi: 10.1038/s41591-024-03150-z. PubMed PMID: 39095595; PMCID: PMC11405287.

41. Reed AD, Pensa S, Steif A, Stenning J, Kunz DJ, Porter LJ, Hua K, He P, Twigger AJ, Siu AJQ, Kania K, Barrow-McGee R, Goulding I, Gomm JJ, Speirs V, Jones JL, Marioni JC, Khaled WT. A single-cell atlas enables mapping of homeostatic cellular shifts in the adult human breast. Nat Genet. 2024;56(4):652–62. Epub 2024/03/29. doi: 10.1038/s41588-024-01688-9. PubMed PMID: 38548988; PMCID: PMC11018528 declare no competing interests.

42. Wu Y, Fan Y, Miao Y, Li Y, Du G, Chen Z, Diao J, Chen YA, Ye M, You R, Chen A, Chen Y, Li W, Guo W, Dong J, Zhang X, Wang Y, Gu J. uniLIVER: a human liver cell atlas for data-driven cellular state mapping. J Genet Genomics. 2025;52(9):1133–47. Epub 2025/02/02. doi: 10.1016/j.jgg.2025.01.017. PubMed PMID: 39892777.

43. Oliver AJ, Huang N, Bartolome-Casado R, Li R, Koplev S, Nilsen HR, Moy M, Cakir B, Polanski K, Gudiño V, Melón-Ardanaz E, Sumanaweera D, Dimitrov D, Milchsack LM, FitzPatrick MEB, Provine NM, Boccacino JM, Dann E, Predeus AV, To K, Prete M, Chapman JA, Masi AC, Stephenson E, Engelbert J, Lobentanzer S, Perera S, Richardson L, Kapuge R, Wilbrey-Clark A, Semprich CI, Ellams S, Tudor C, Joseph P, Garrido-Trigo A, Corraliza AM, Oliver TRW, Hook CE, James KR, Mahbubani KT, Saeb-Parsy K, Zilbauer M, Saez-Rodriguez J, Høivik ML, Bækkevold ES, Stewart CJ, Berrington JE, Meyer KB, Klenerman P, Salas A, Haniffa M, Jahnsen FL, Elmentaite R, Teichmann SA. Single-cell integration reveals metaplasia in inflammatory gut diseases. Nature. 2024;635(8039):699–707. Epub 2024/11/21. doi: 10.1038/s41586-024-07571-1. PubMed PMID: 39567783; PMCID: PMC11578898 Labs, OMass Therapeutics, a co-founder and equity holder of TransitionBio and EnsoCell Therapeutics, a non-executive director of 10x Genomics and a part-time employee of GlaxoSmithKline. R.E. is an equity holder in EnsoCell. P.K. has consulted for AstraZeneca, UCB, Biomunex and Infinitopes. N.M.P reports consulting fees from Infinitopes. J.S.-R. reports funding from GSK, Pfizer and Sanofi and fees/honoraria from Travere Therapeutics, Stadapharm, Astex, Owkin, Pfizer, Moderna and Grunenthal. A.S. is the recipient of research grants from Roche-Genentech, Abbvie, GSK, Scipher Medicine, Pfizer, Alimentiv, Boehringer Ingelheim and Agomab and has received consulting fees from Genentech, GSK, Pfizer, HotSpot Therapeutics, Alimentiv, Agomab, Goodgut and Orikine. R.E. and S.A.T are inventors on the patent GB2412853.0 filed in the UK, some components of which are related to this work. All other authors declare no competing interests.

44. Salcher S, Sturm G, Horvath L, Untergasser G, Kuempers C, Fotakis G, Panizzolo E, Martowicz A, Trebo M, Pall G, Gamerith G, Sykora M, Augustin F, Schmitz K, Finotello F, Rieder D, Perner S, Sopper S, Wolf D, Pircher A, Trajanoski Z. High-resolution single-cell atlas reveals diversity and plasticity of tissue-resident neutrophils in non-small cell lung cancer. Cancer Cell. 2022;40(12):1503–20.e8. Epub 2022/11/12. doi: 10.1016/j.ccell.2022.10.008. PubMed PMID: 36368318; PMCID: PMC9767679.

45. Suo C, Dann E, Goh I, Jardine L, Kleshchevnikov V, Park JE, Botting RA, Stephenson E, Engelbert J, Tuong ZK, Polanski K, Yayon N, Xu C, Suchanek O, Elmentaite R, Domínguez Conde C, He P, Pritchard S, Miah M, Moldovan C, Steemers AS, Mazin P, Prete M, Horsfall D, Marioni JC, Clatworthy MR, Haniffa M, Teichmann SA. Mapping the developing human immune system across organs. Science. 2022;376(6597):eabo0510. Epub 2022/05/14. doi: 10.1126/science.abo0510. PubMed PMID: 35549310; PMCID: PMC7612819.

46. Herpelinck T, Ory L, Verbraeken T, Nasello G, Barzegari M, Bolander J, Luyten FP, Tylzanowski P, Geris L. A single-cell atlas of the murine limb skeleton integrating the developmental and adult stages. Sci Rep. 2025;15(1):22514. Epub 2025/07/02. doi: 10.1038/s41598-025-05277-6. PubMed PMID: 40596066; PMCID: PMC12215971.

47. Kumar T, Nee K, Wei R, He S, Nguyen QH, Bai S, Blake K, Pein M, Gong Y, Sei E, Hu M, Casasent AK, Thennavan A, Li J, Tran T, Chen K, Nilges B, Kashikar N, Braubach O, Ben Cheikh B, Nikulina N, Chen H, Teshome M, Menegaz B, Javaid H, Nagi C, Montalvan J, Lev T, Mallya S, Tifrea DF, Edwards R, Lin E, Parajuli R, Hanson S, Winocour S, Thompson A, Lim B, Lawson DA, Kessenbrock K, Navin N. A spatially resolved single-cell genomic atlas of the adult human breast. Nature. 2023;620(7972):181–91. Epub 2023/06/29. doi: 10.1038/s41586-023-06252-9. PubMed PMID: 37380767; PMCID: PMC11443819.

48. Elmentaite R, Kumasaka N, Roberts K, Fleming A, Dann E, King HW, Kleshchevnikov V, Dabrowska M, Pritchard S, Bolt L, Vieira SF, Mamanova L, Huang N, Perrone F, Goh Kai’En I, Lisgo SN, Katan M, Leonard S, Oliver TRW, Hook CE, Nayak K, Campos LS, Domínguez Conde C, Stephenson E, Engelbert J, Botting RA, Polanski K, van Dongen S, Patel M, Morgan MD, Marioni JC, Bayraktar OA, Meyer KB, He X, Barker RA, Uhlig HH, Mahbubani KT, Saeb-Parsy K, Zilbauer M, Clatworthy MR, Haniffa M, James KR, Teichmann SA. Cells of the human intestinal tract mapped across space and time. Nature. 2021;597(7875):250–5. Epub 2021/09/10. doi: 10.1038/s41586-021-03852-1. PubMed PMID: 34497389; PMCID: PMC8426186 advisory boards at Roche, Qiagen, Genentech, Biogen, GlaxoSmithKline and ForeSite Labs. The remaining authors declare no competing interests.

49. Edgar RD, Nakib D, Camat D, Chung S, Lumanto P, Atif J, Perciani CT, Ma XZ, Thoeni C, Selvakumaran N, Manuel J, Sayed B, Huysentruyt K, Ricciuto A, McGilvray I, Avitzur Y, Bader GD, MacParland SA. Single-cell atlas of human pediatric liver reveals age-related hepatic gene signatures. Hepatol Commun. 2025;9(11). Epub 2025/10/07. doi: 10.1097/hc9.0000000000000813. PubMed PMID: 41056487; PMCID: PMC12506995.

50. Park JE, Botting RA, Domínguez Conde C, Popescu DM, Lavaert M, Kunz DJ, Goh I, Stephenson E, Ragazzini R, Tuck E, Wilbrey-Clark A, Roberts K, Kedlian VR, Ferdinand JR, He X, Webb S, Maunder D, Vandamme N, Mahbubani KT, Polanski K, Mamanova L, Bolt L, Crossland D, de Rita F, Fuller A, Filby A, Reynolds G, Dixon D, Saeb-Parsy K, Lisgo S, Henderson D, Vento-Tormo R, Bayraktar OA, Barker RA, Meyer KB, Saeys Y, Bonfanti P, Behjati S, Clatworthy MR, Taghon T, Haniffa M, Teichmann SA. A cell atlas of human thymic development defines T cell repertoire formation. Science. 2020;367(6480). Epub 2020/02/23. doi: 10.1126/science.aay3224. PubMed PMID: 32079746; PMCID: PMC7611066.

51. Maan H, Zhang L, Yu C, Geuenich MJ, Campbell KR, Wang B. Characterizing the impacts of dataset imbalance on single-cell data integration. Nat Biotechnol. 2024;42(12):1899-908. Epub 2024/03/02. doi: 10.1038/s41587-023-02097-9. PubMed PMID: 38429430.

52. Wang S, Pisco AO, McGeever A, Brbic M, Zitnik M, Darmanis S, Leskovec J, Karkanias J, Altman RB. Leveraging the Cell Ontology to classify unseen cell types. Nat Commun. 2021;12(1):5556. Epub 2021/09/23. doi: 10.1038/s41467-021-25725-x. PubMed PMID: 34548483; PMCID: PMC8455606 (Personalis, 23andme); consulting or advisory role (United Health, Second Genome, Karius, UK Biobank, Swiss Personalized Health Network, Danish National Genome Center, All of Us Project, and Bridge Bio).

53. Cui H, Wang C, Maan H, Pang K, Luo F, Duan N, Wang B. scGPT: toward building a foundation model for single-cell multi-omics using generative AI. Nat Methods. 2024;21(8):1470–80. Epub 2024/02/27. doi: 10.1038/s41592-024-02201-0. PubMed PMID: 38409223.

54. Saelens W, Cannoodt R, Todorov H, Saeys Y. A comparison of single-cell trajectory inference methods. Nature biotechnology. 2019;37(5):547–54.

55. Virshup I, Rybakov S, Theis FJ, Angerer P, Wolf FA. anndata: Access and store annotated data matrices. Journal of Open Source Software. 2024;9(101):4371.

56. Wolf FA, Angerer P, Theis FJ. SCANPY: large-scale single-cell gene expression data analysis. Genome Biol. 2018;19(1):15. Epub 2018/02/08. doi: 10.1186/s13059-017-1382-0. PubMed PMID: 29409532; PMCID: PMC5802054.

