## Supplementary figures and images for "Benchmarking single cell transcriptome matching methods for incremental growth of cell atlases"

### Supplementary Figure 1

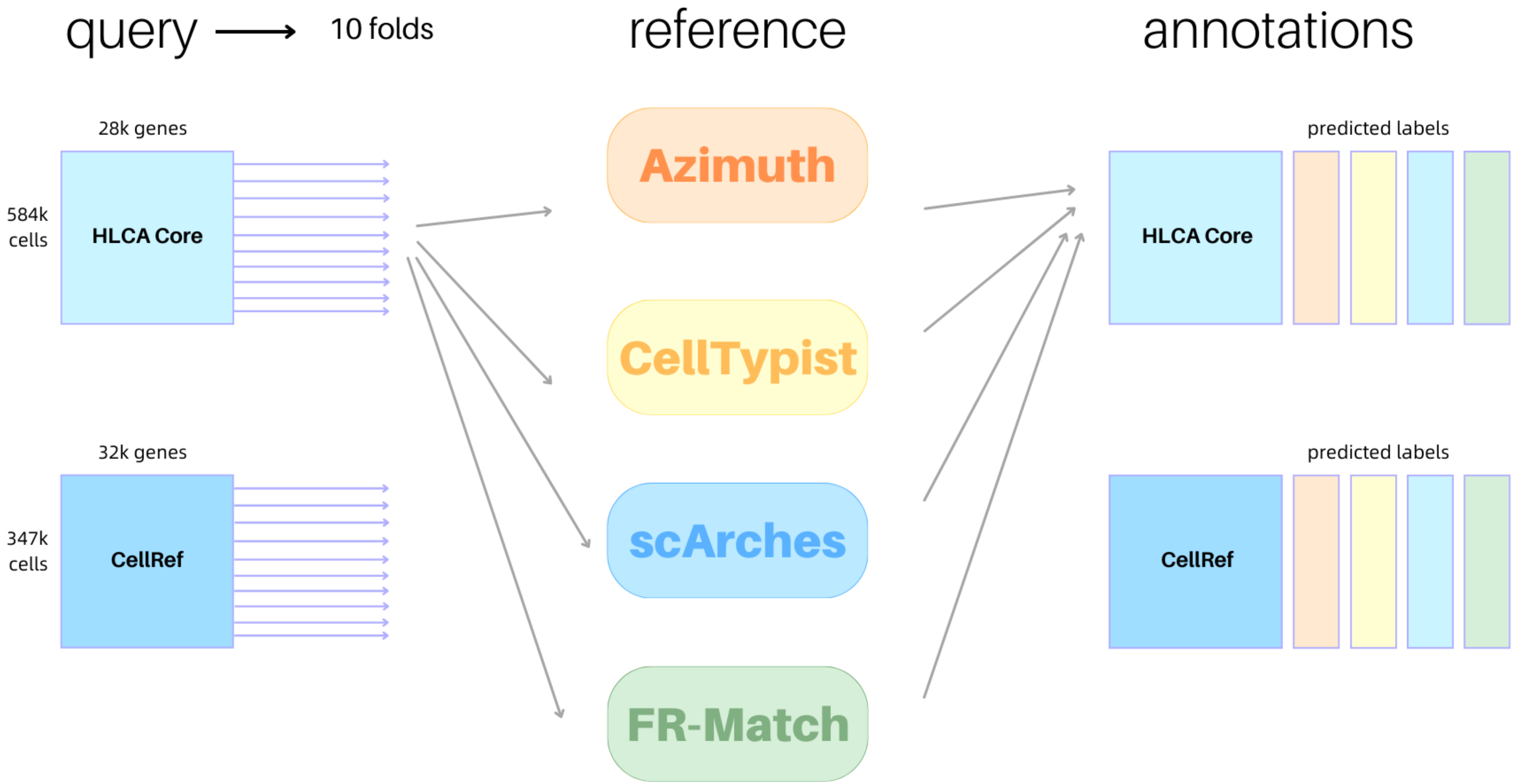

### Supplementary Figure 2

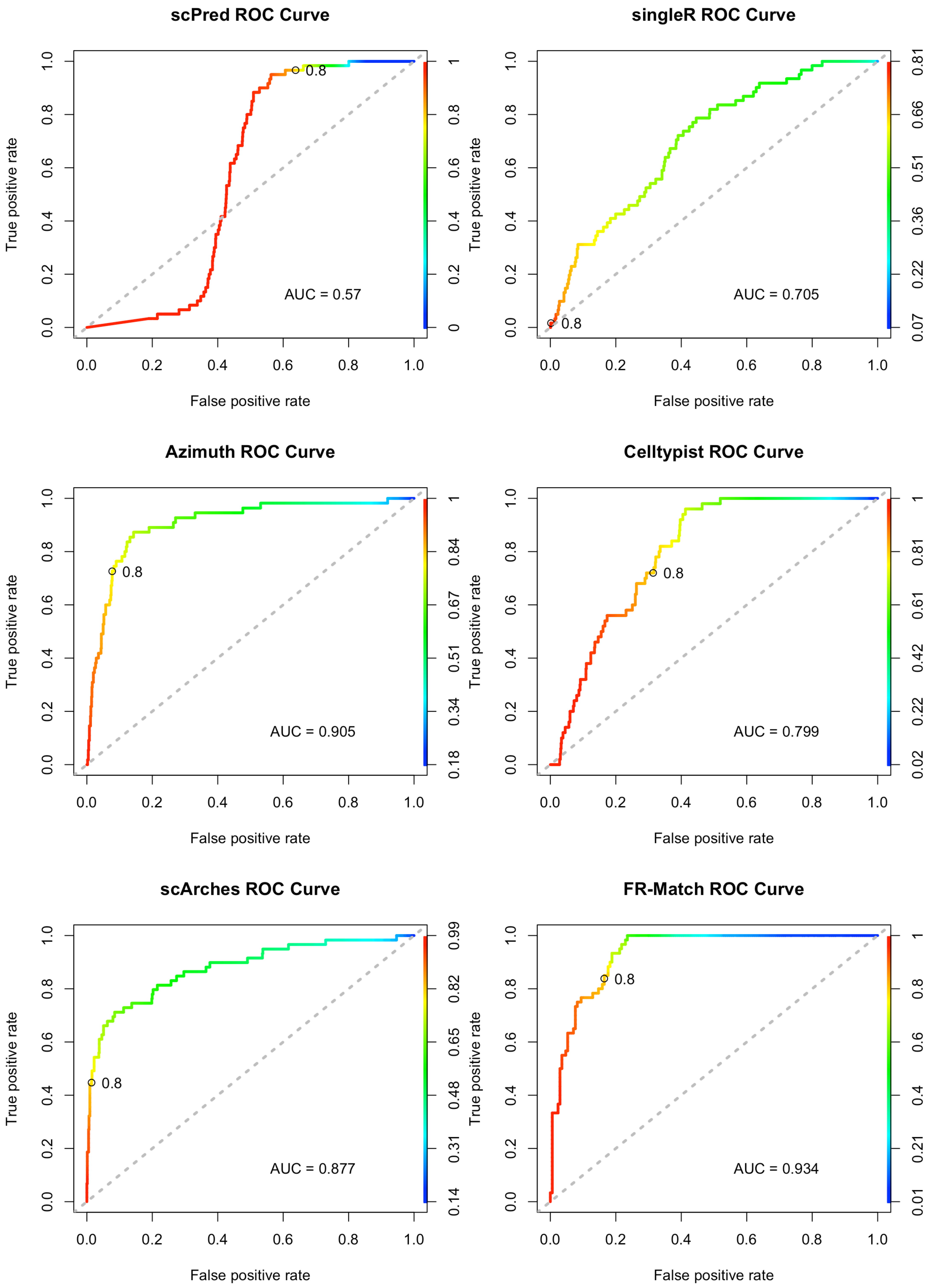

### Supplementary Figure 3

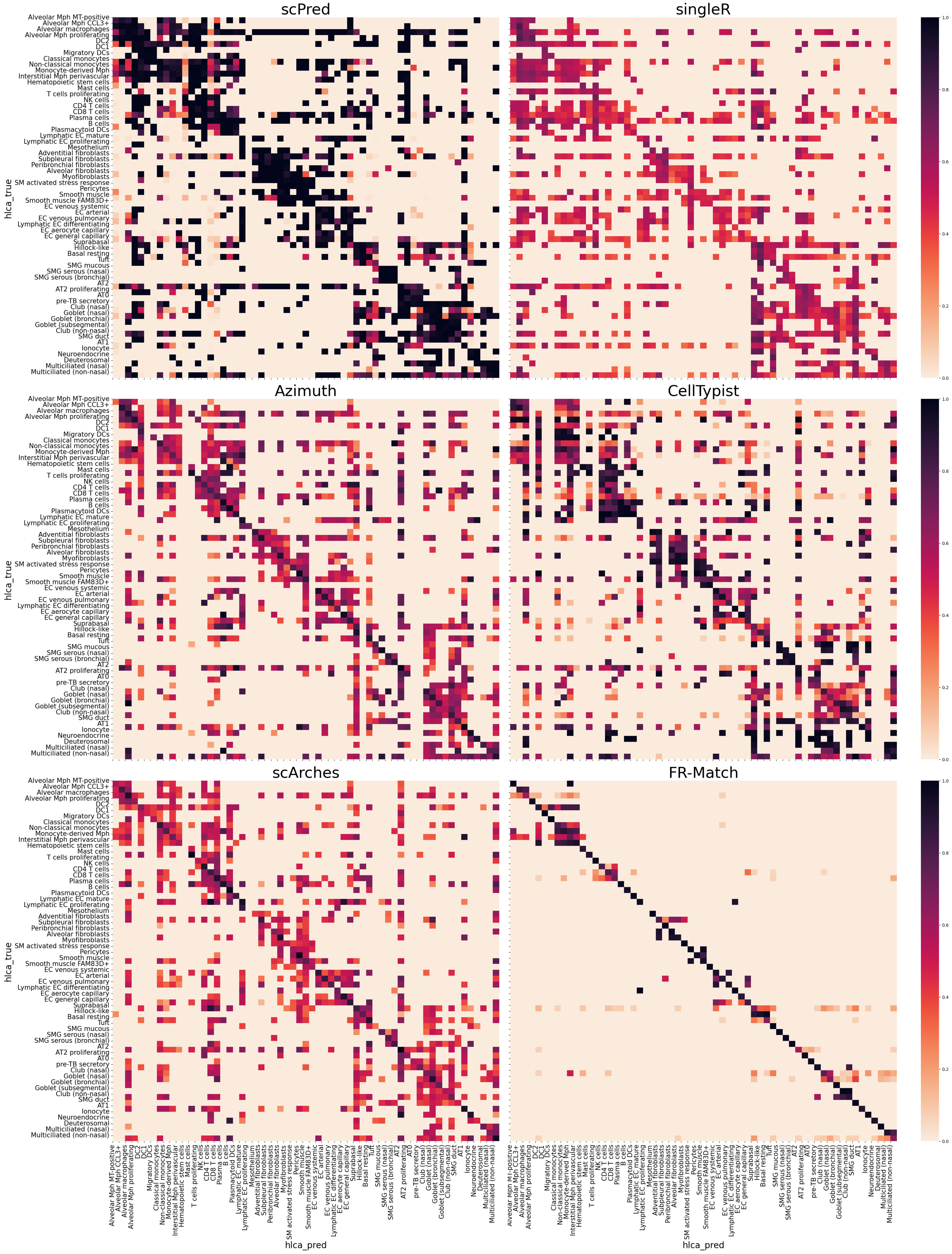

### Supplementary Figure 4

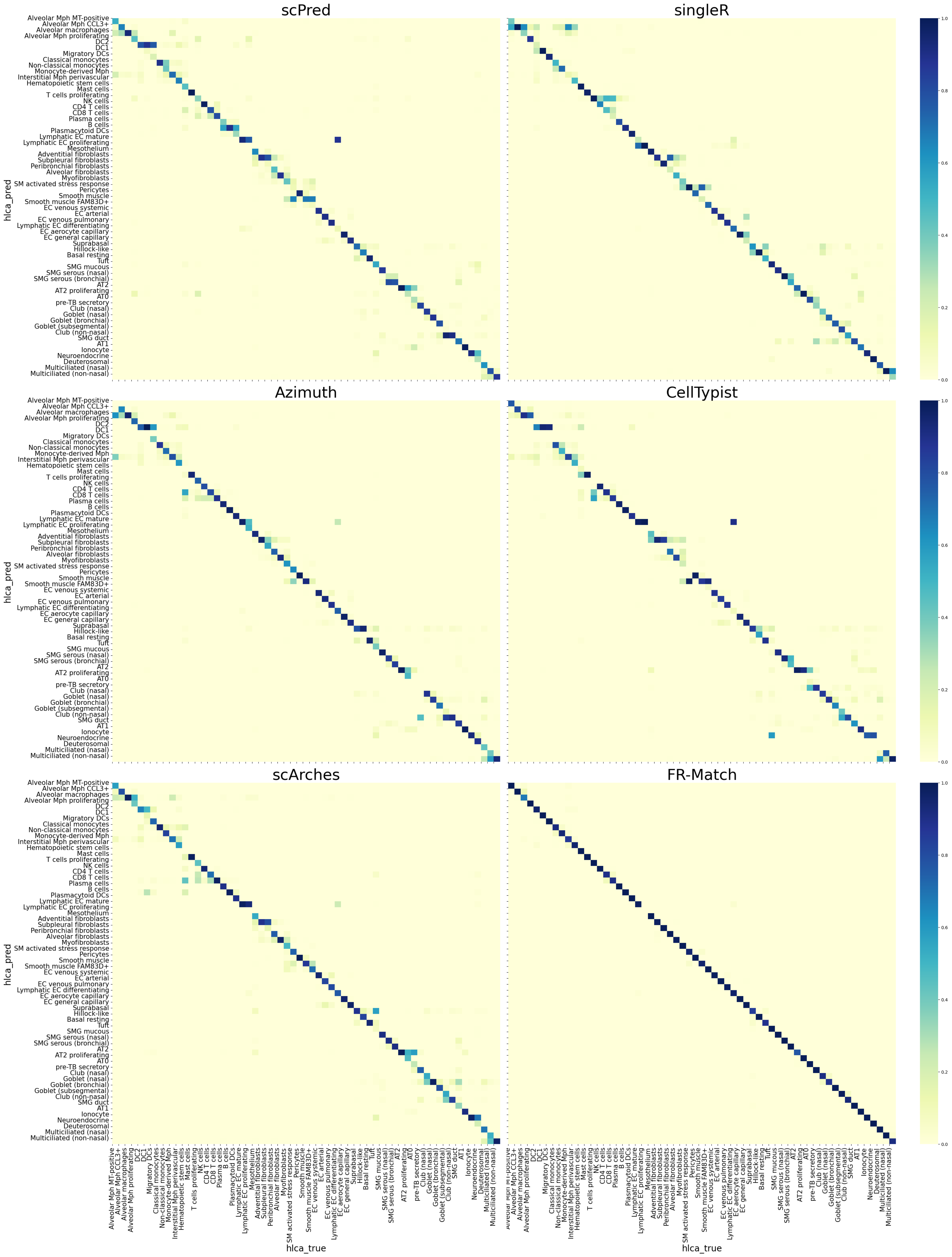

### Supplementary Figure 5

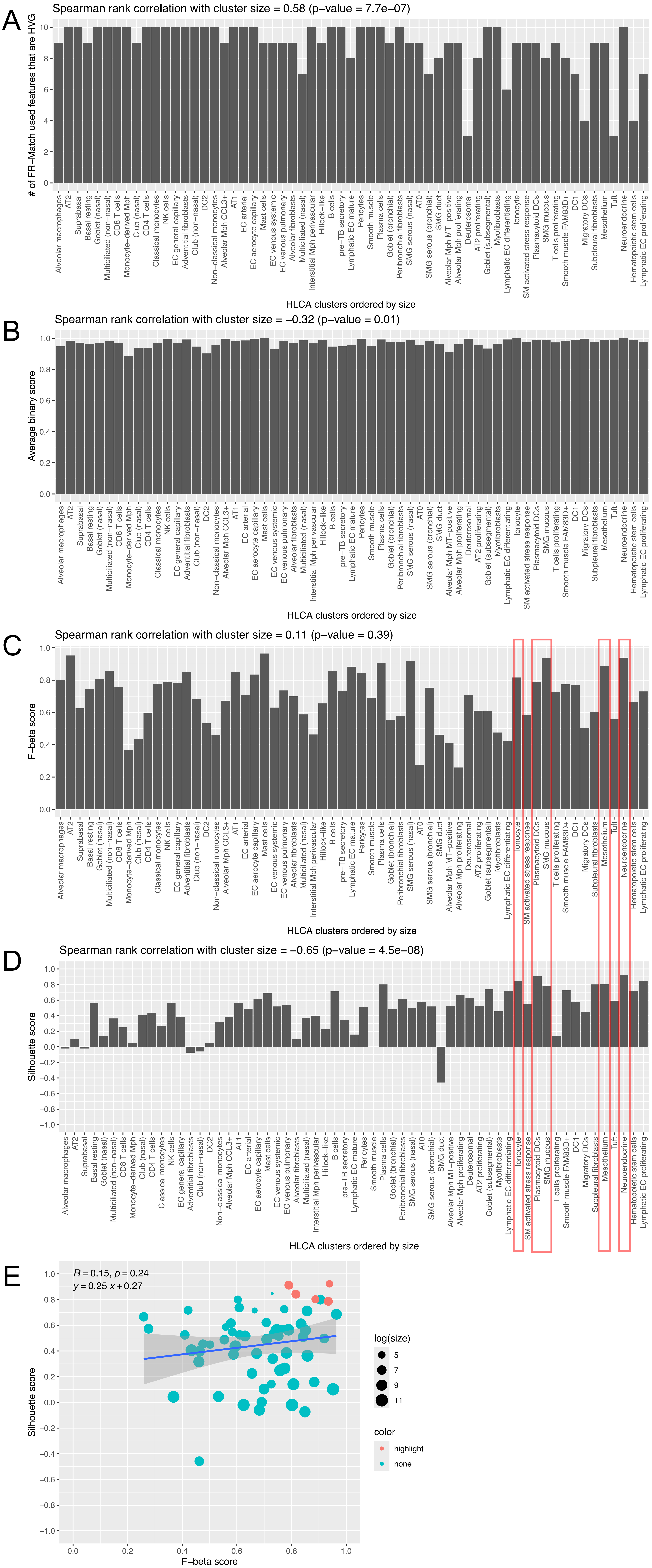

### Supplementary Figure 6

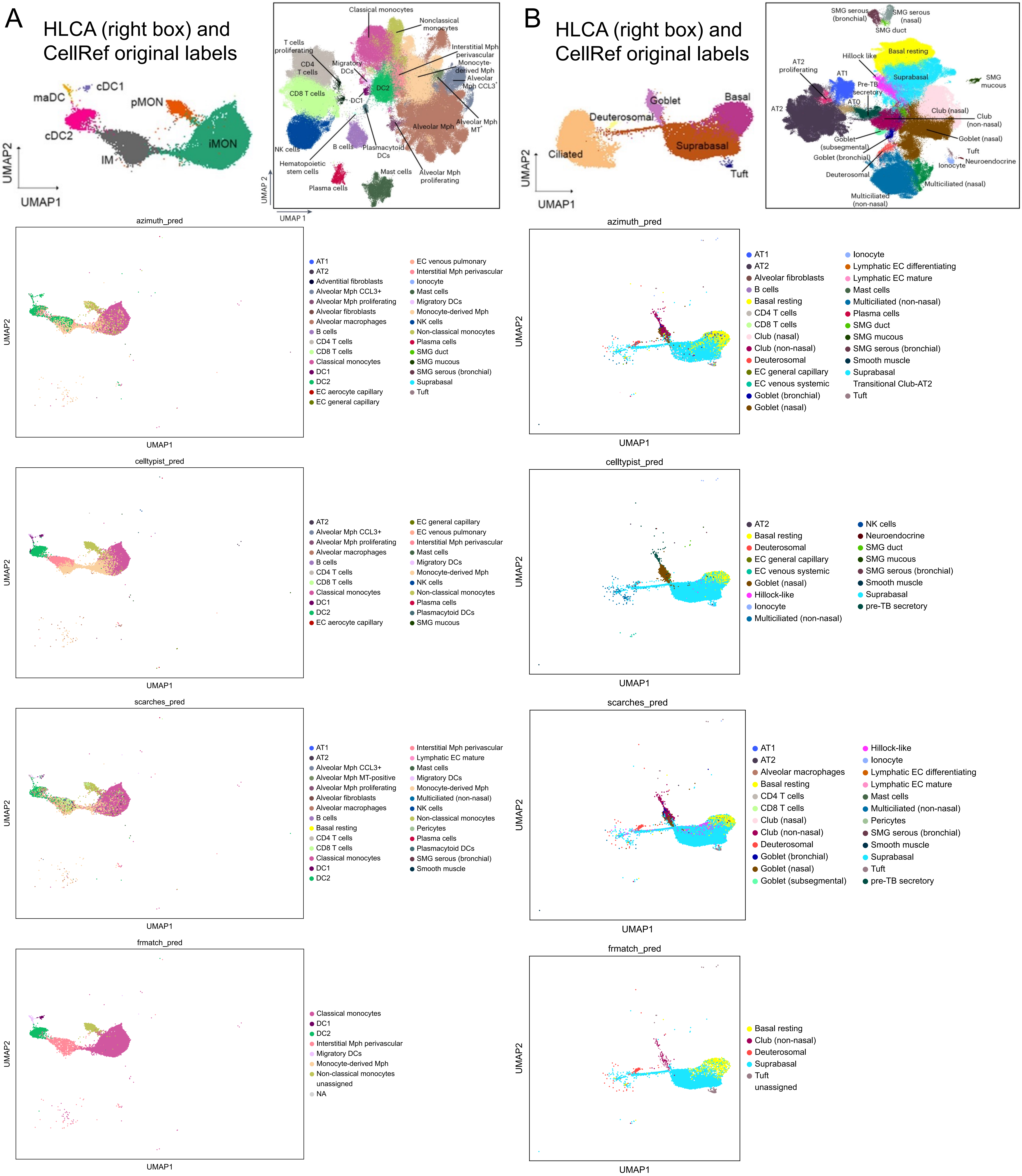

### Supplementary Figure 7

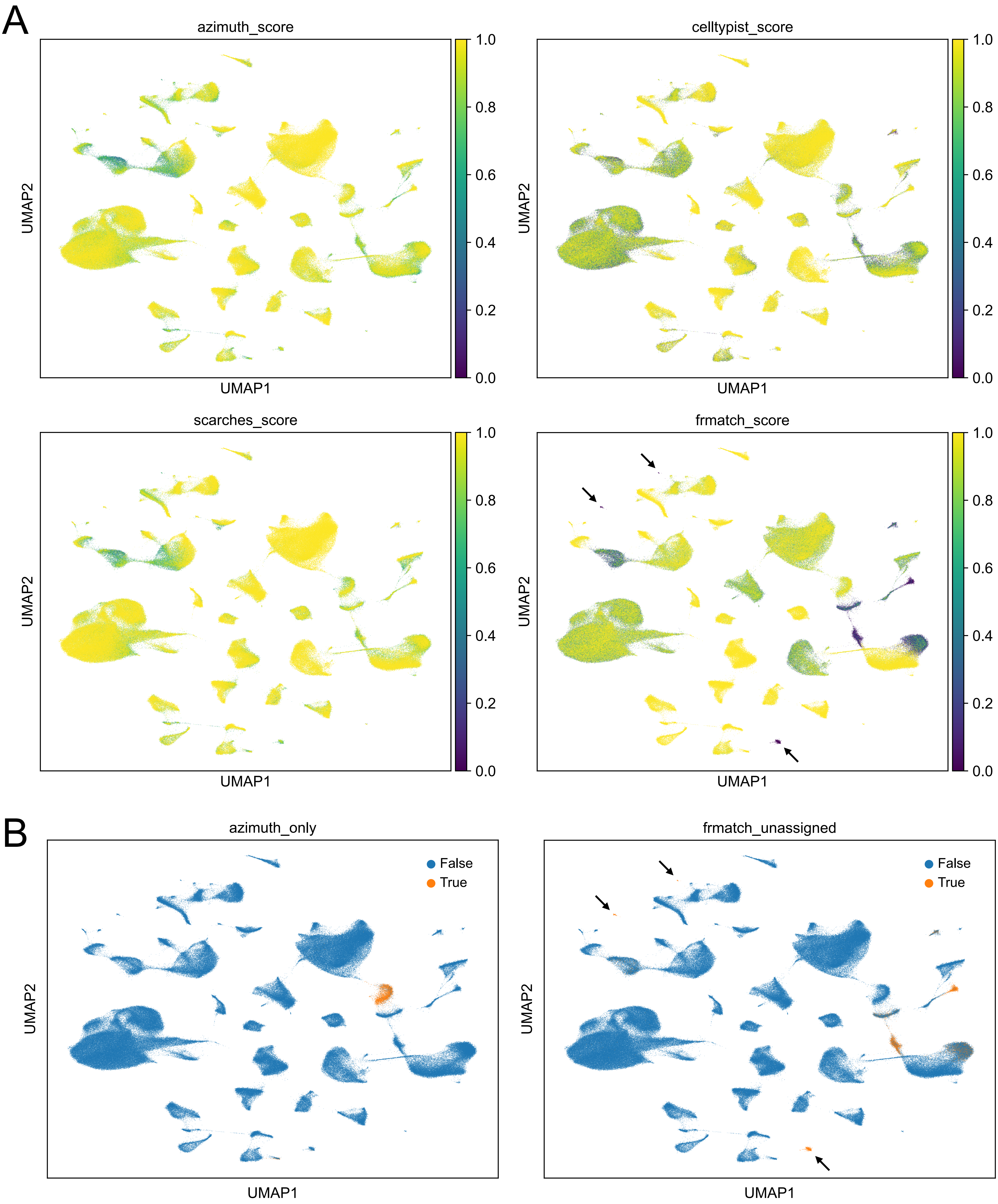

### Supplementary Figure 8

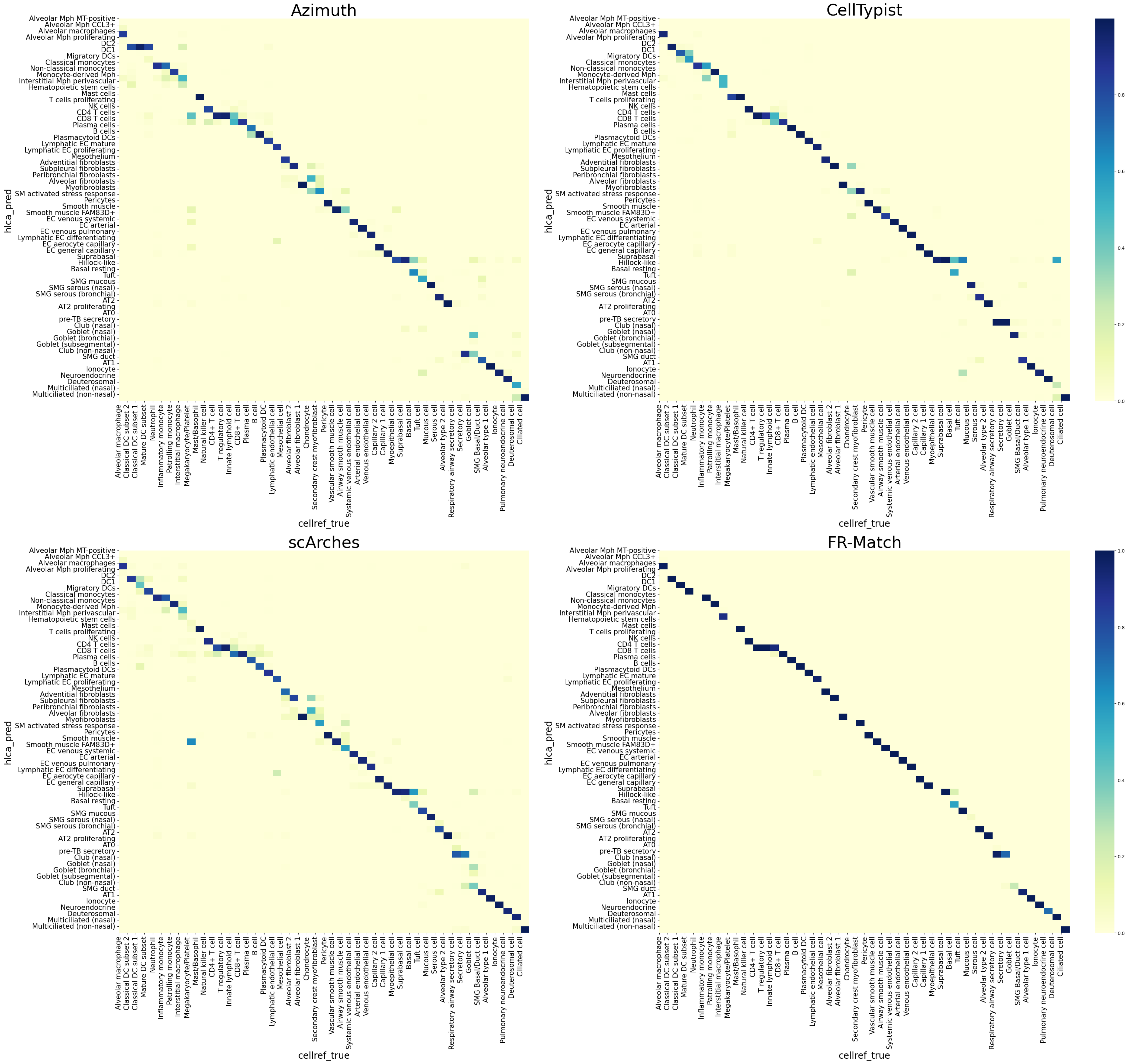

### Supplementary Figure 9

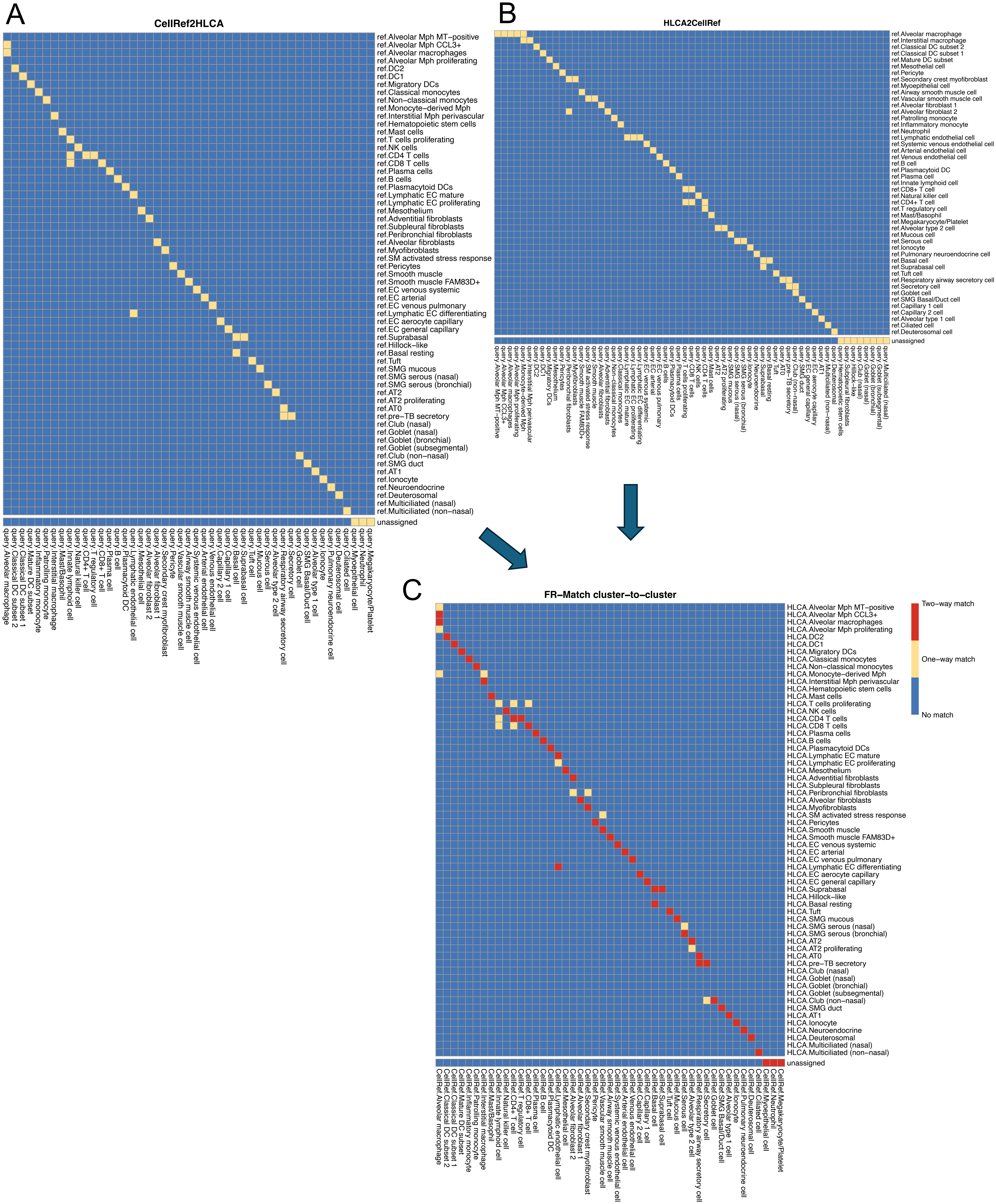

### Supplementary Figure 10

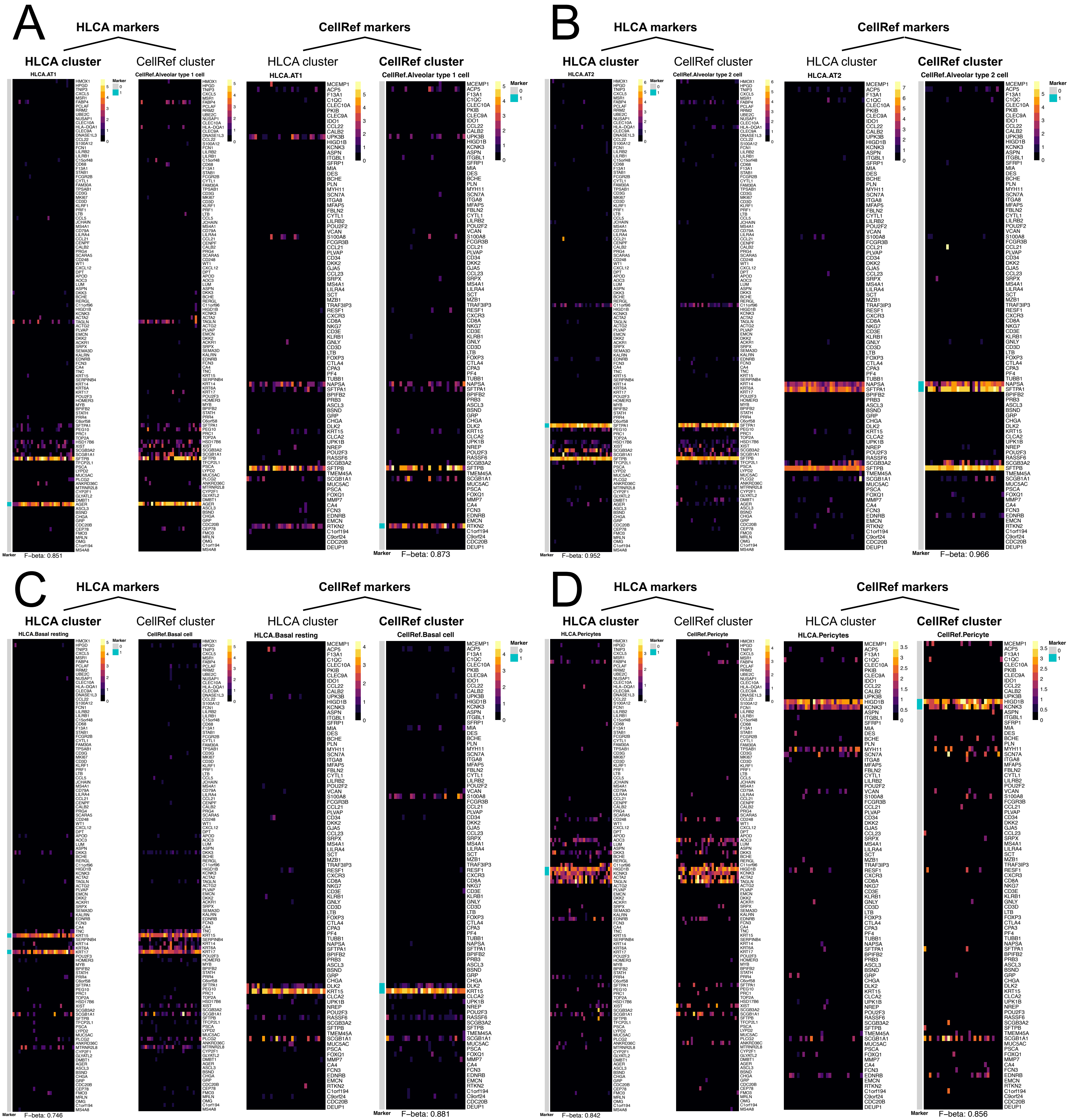

### Supplementary Figure 11

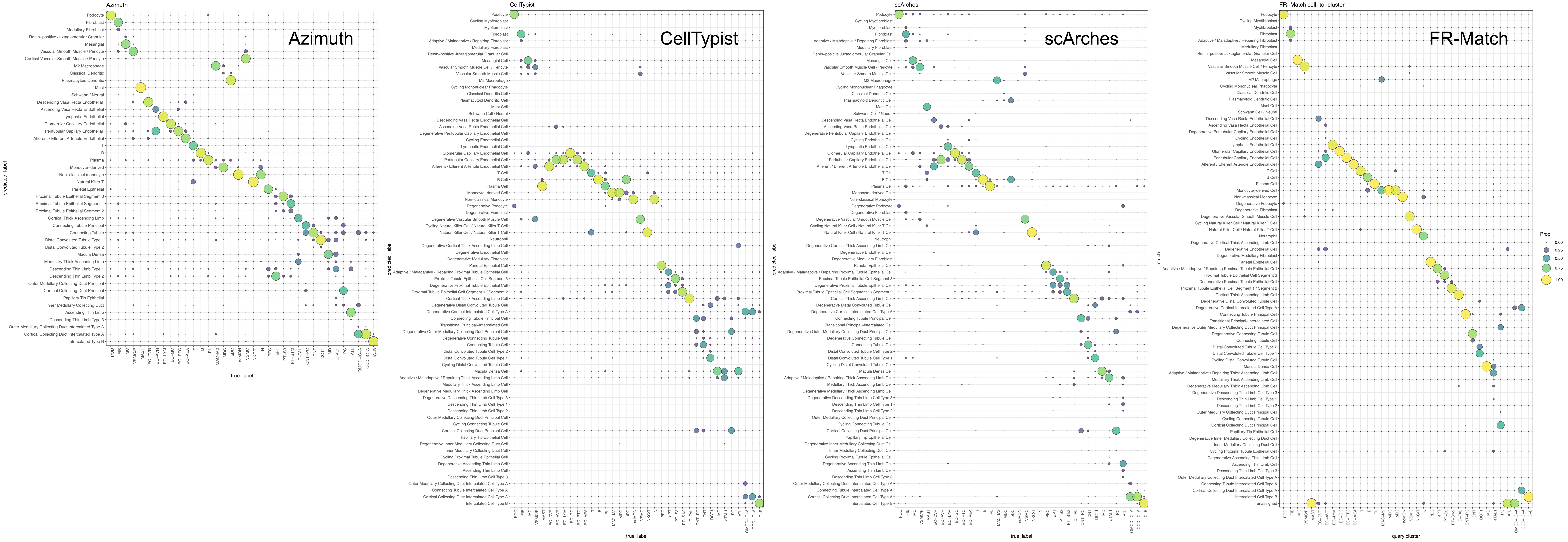

### Supplementary Figure 12

#### Retina

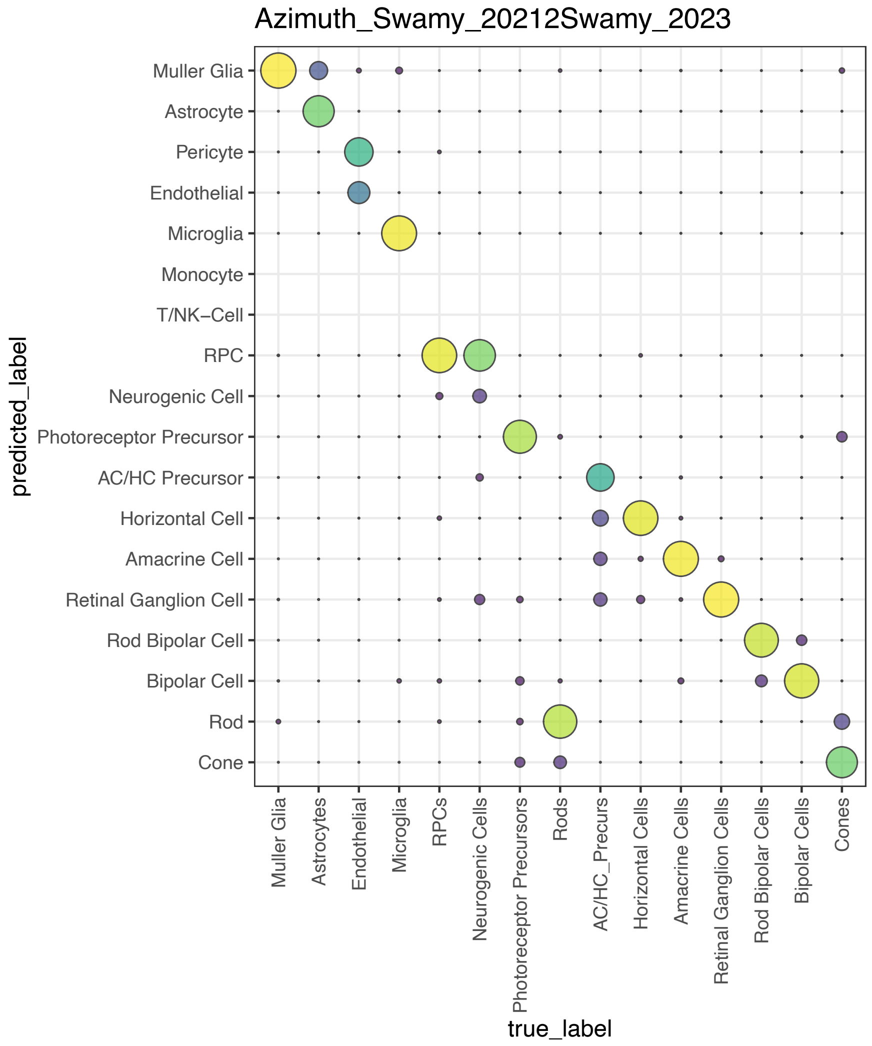


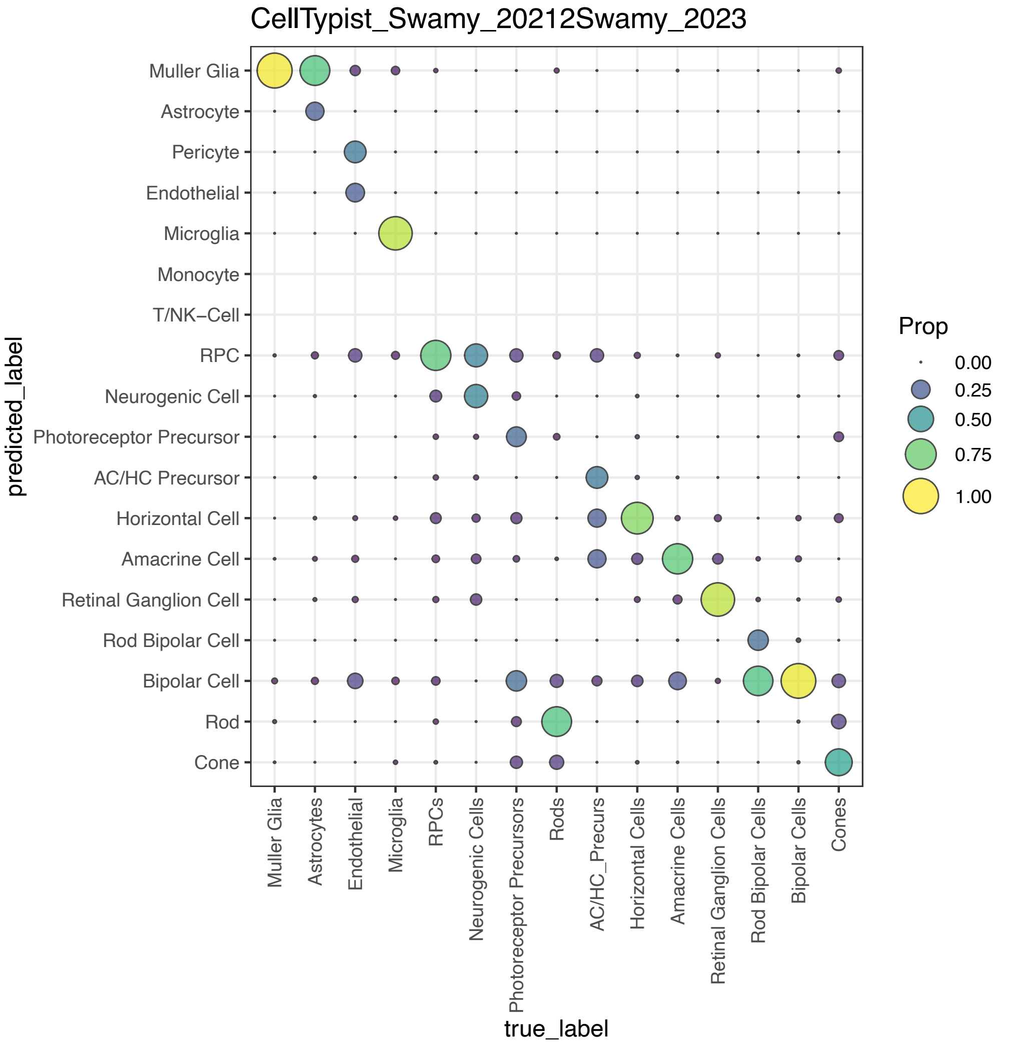


*No count data*

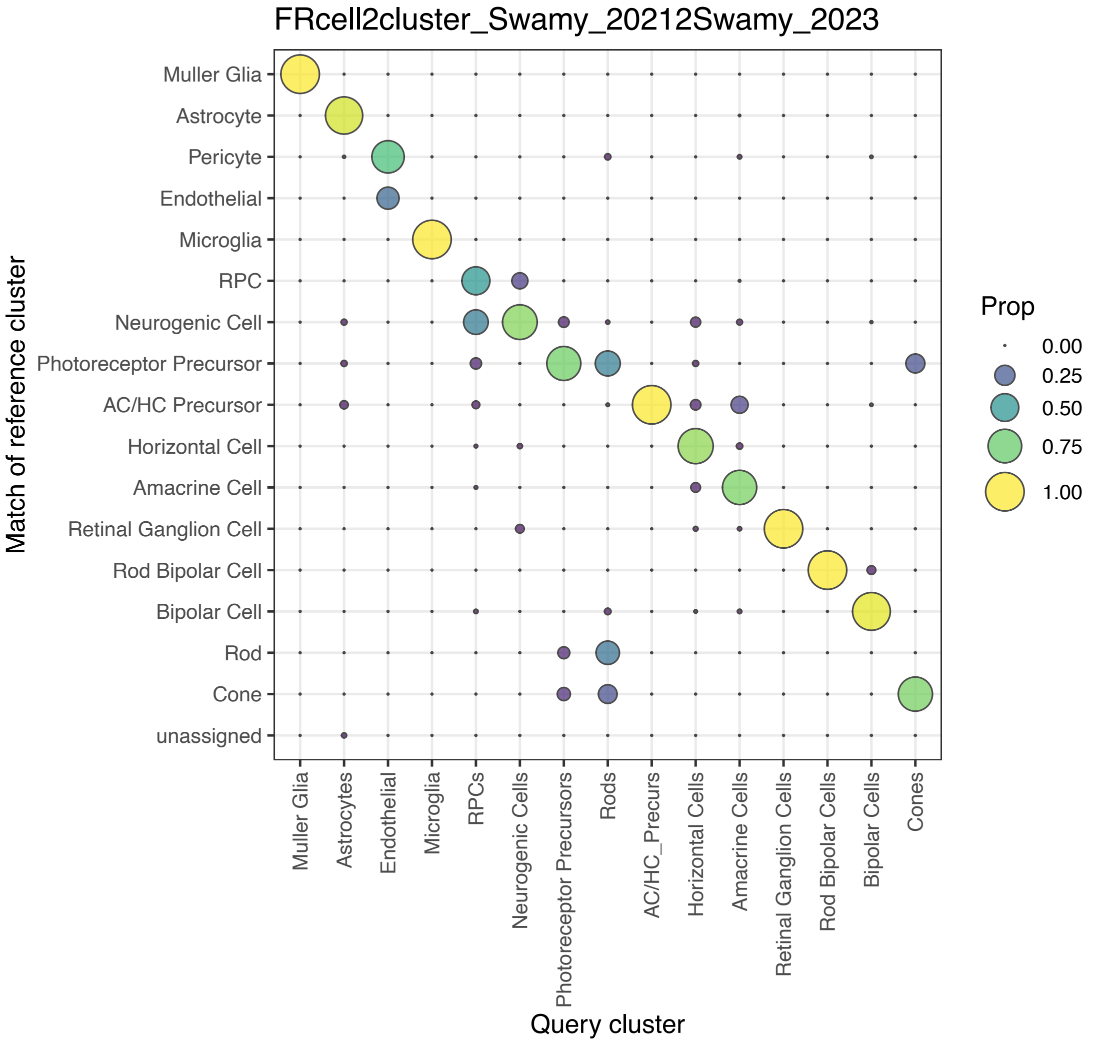


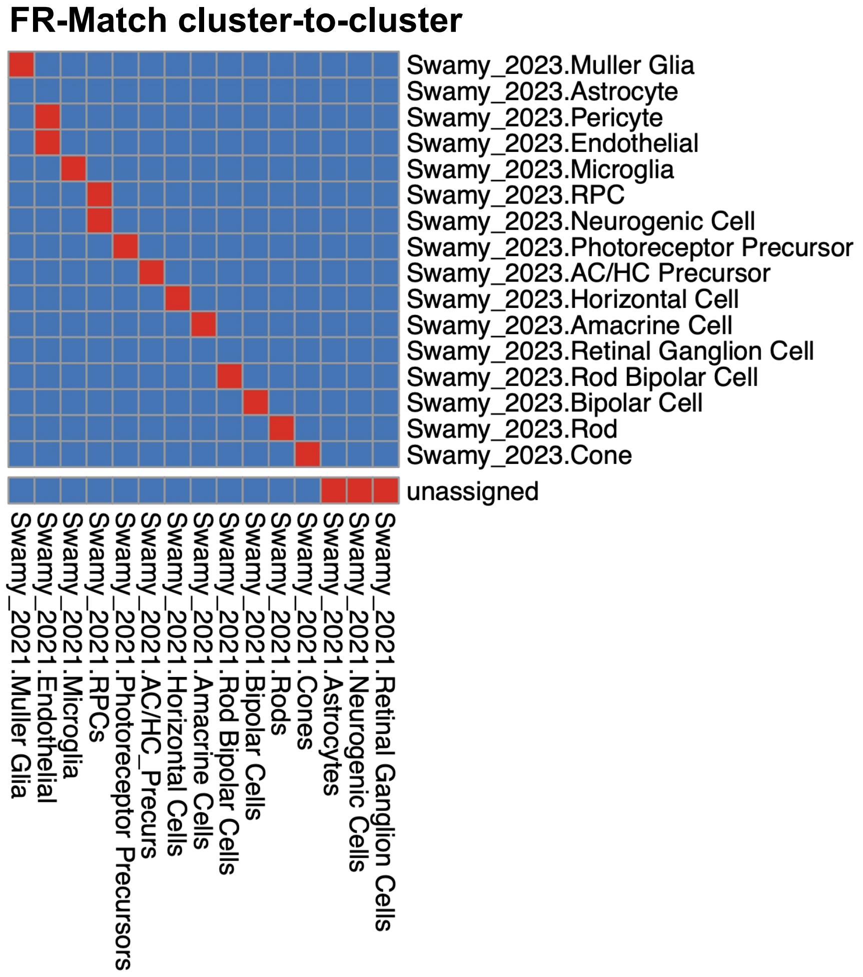


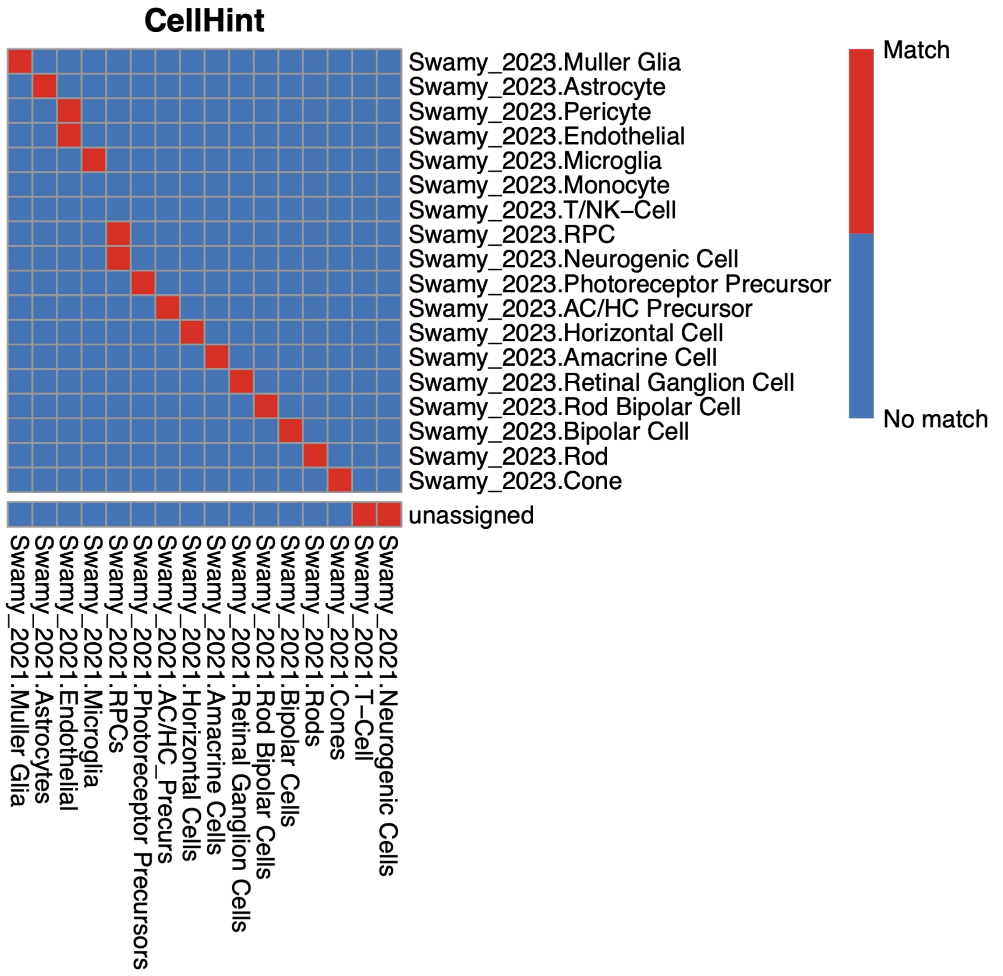

### Supplementary Figure 13

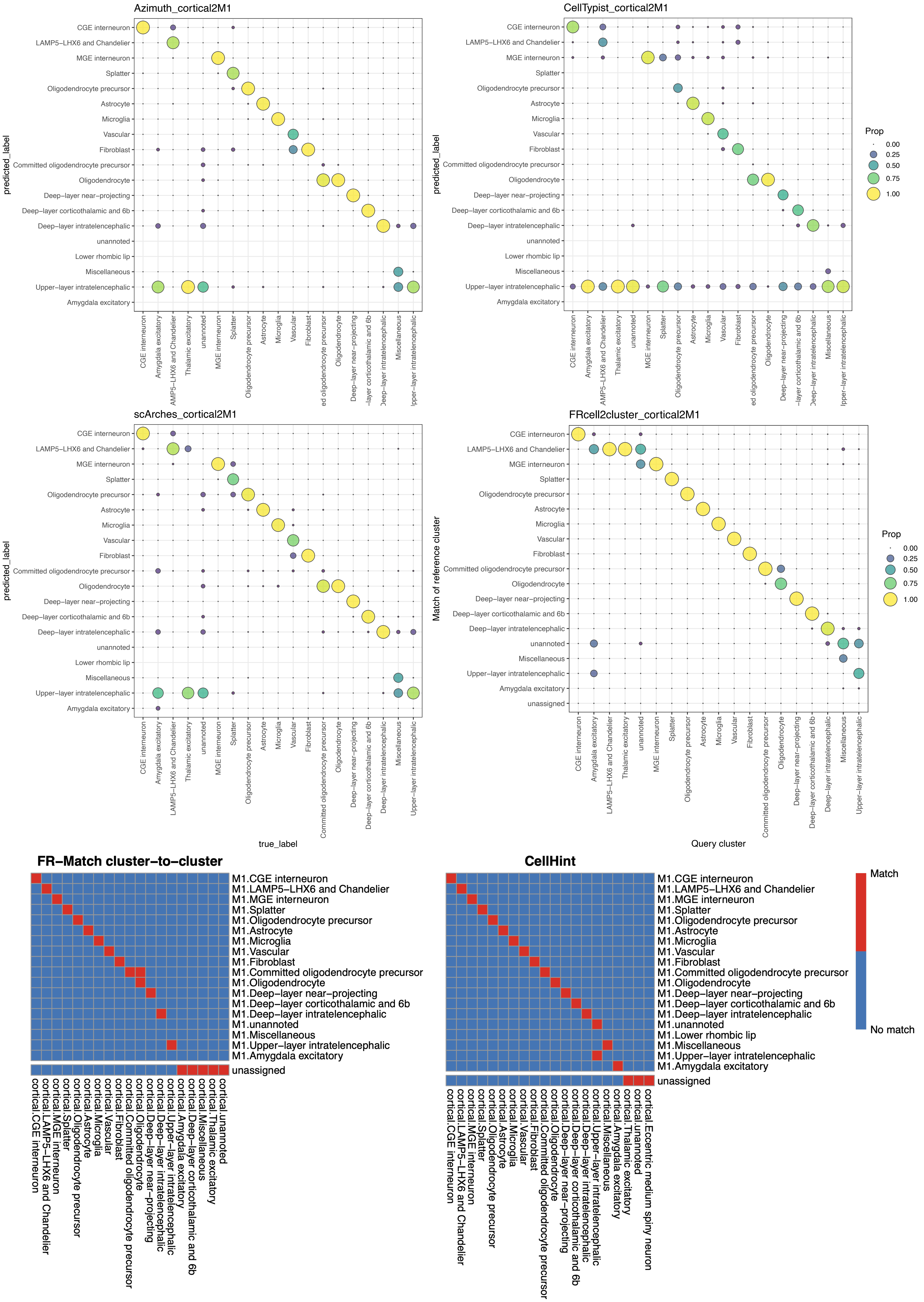

### Supplementary Figure 14

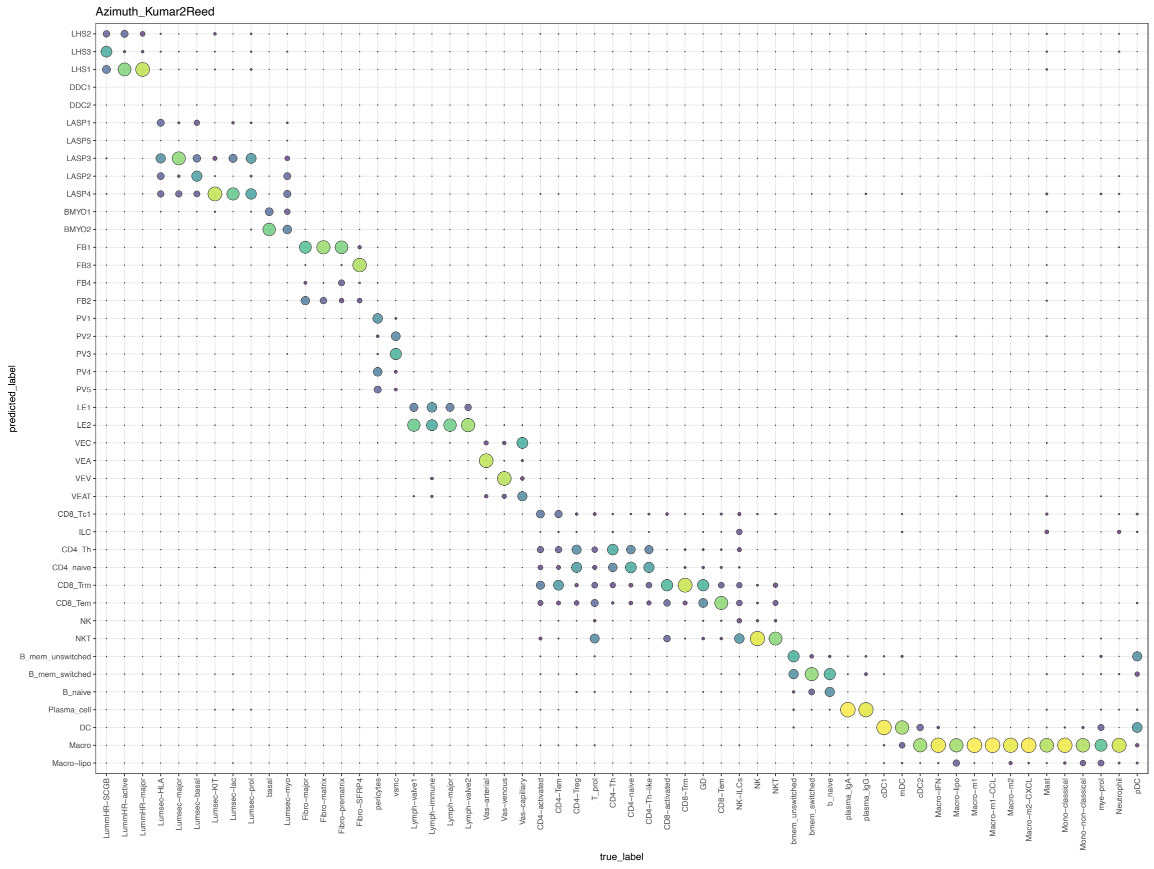

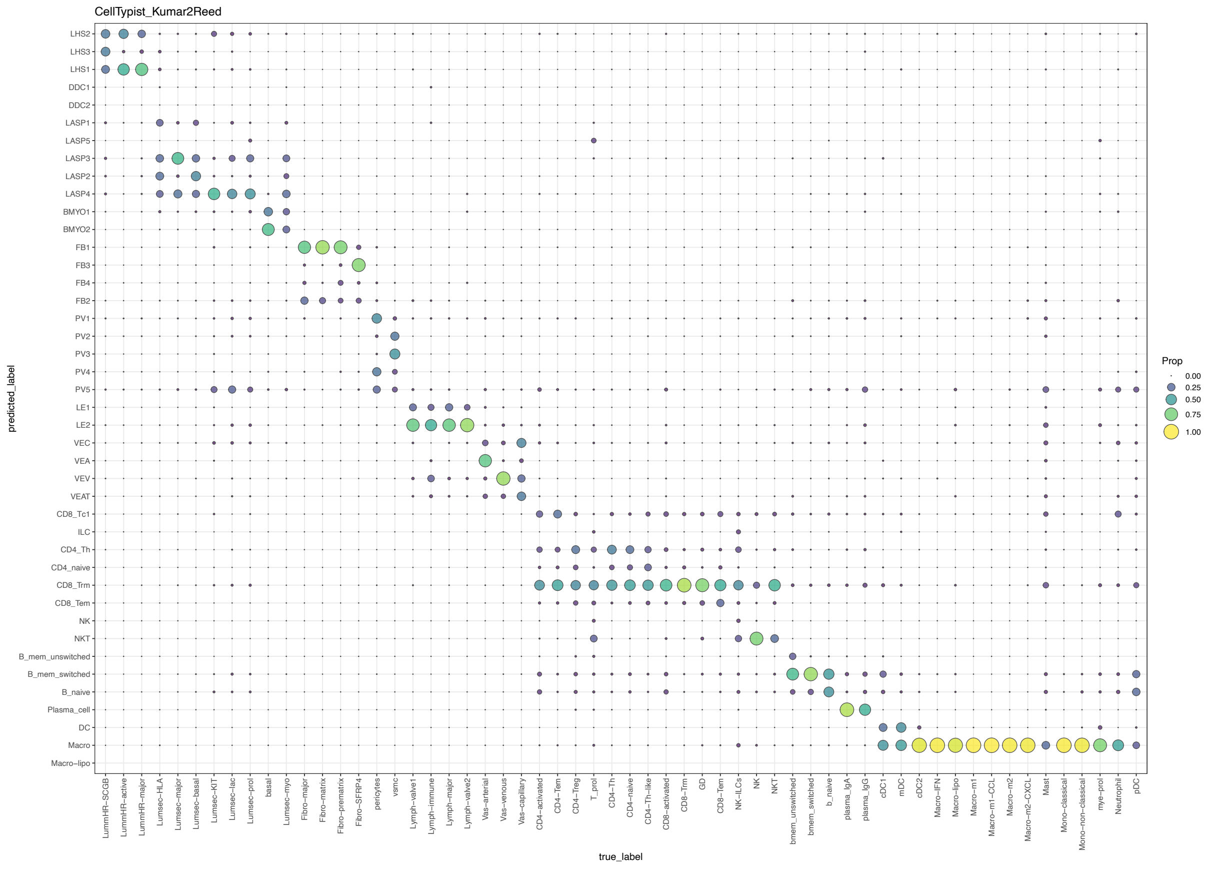

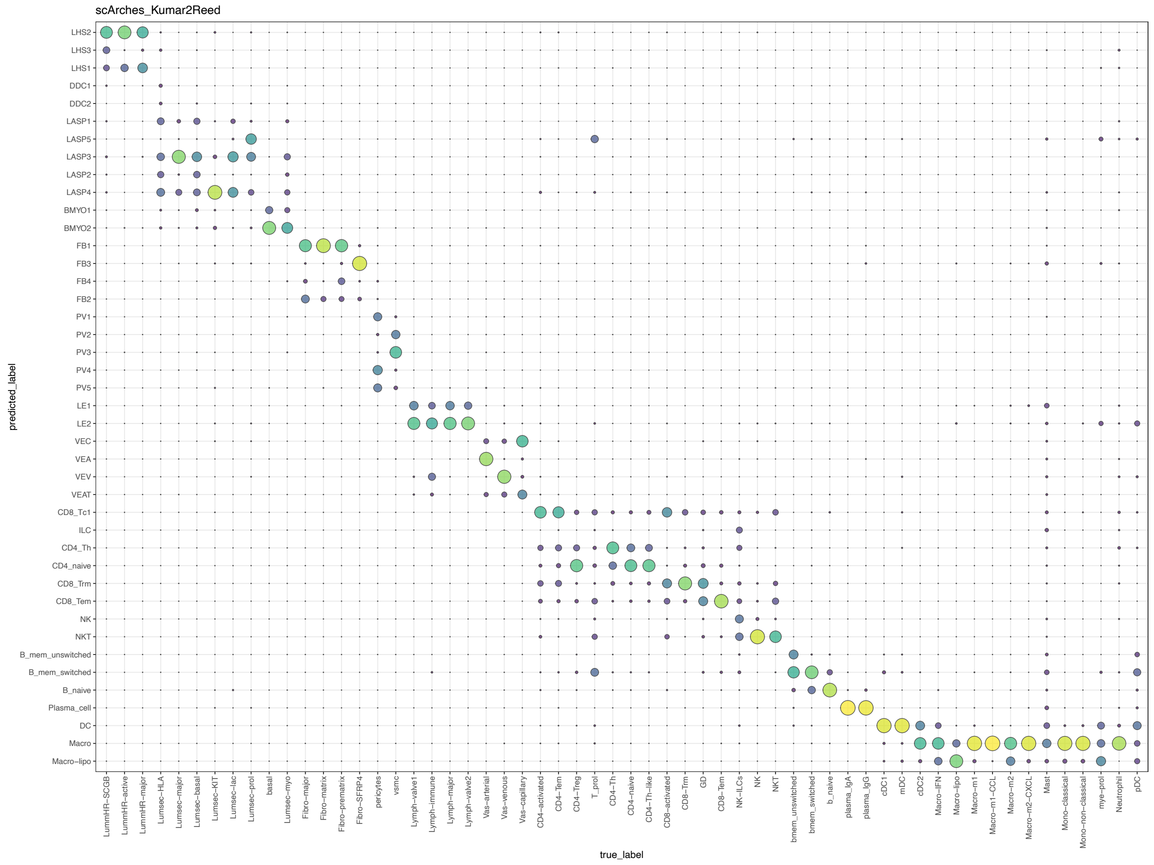

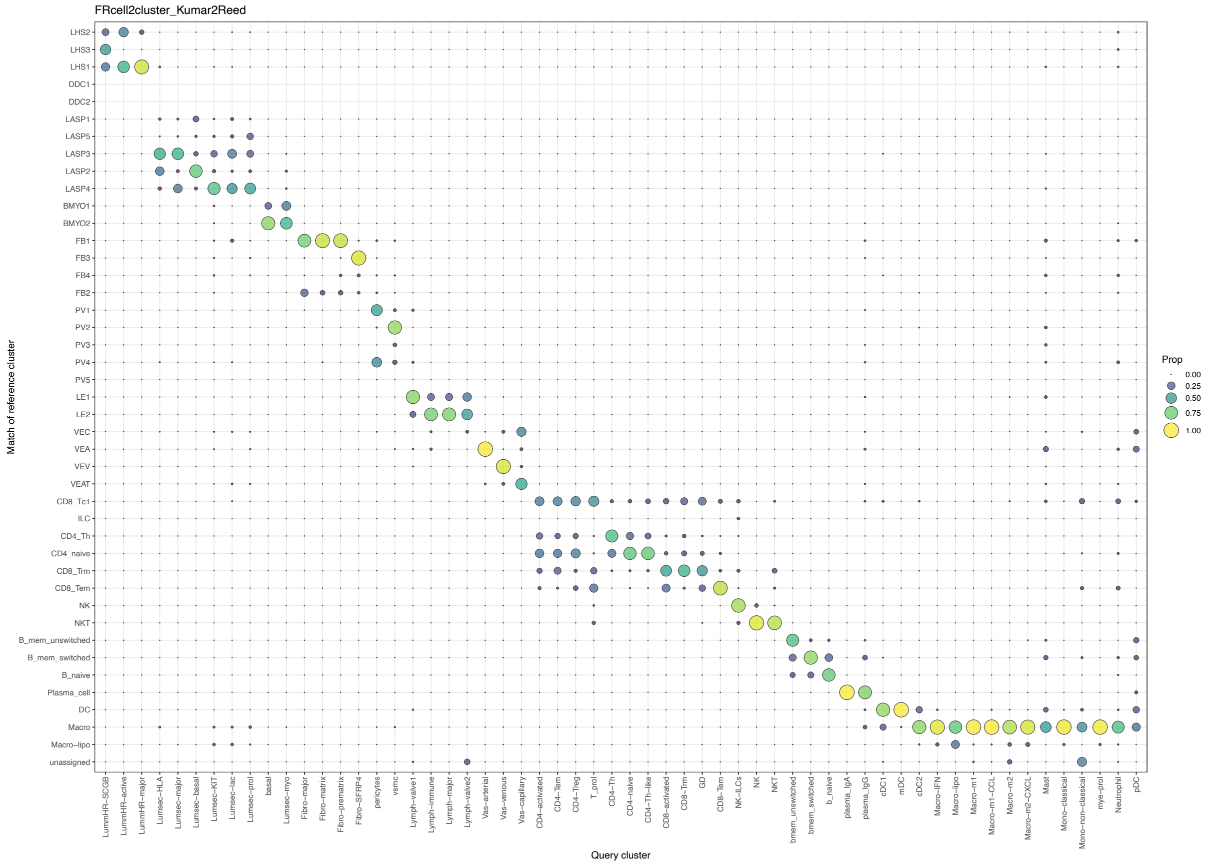

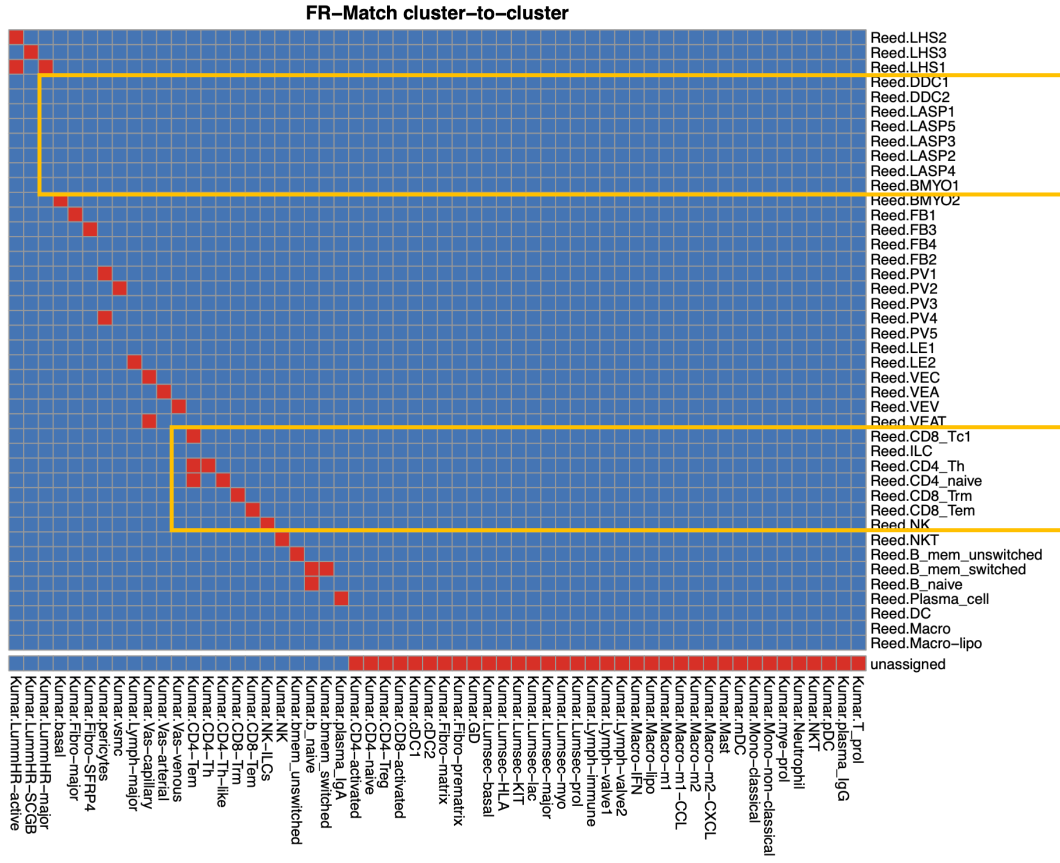

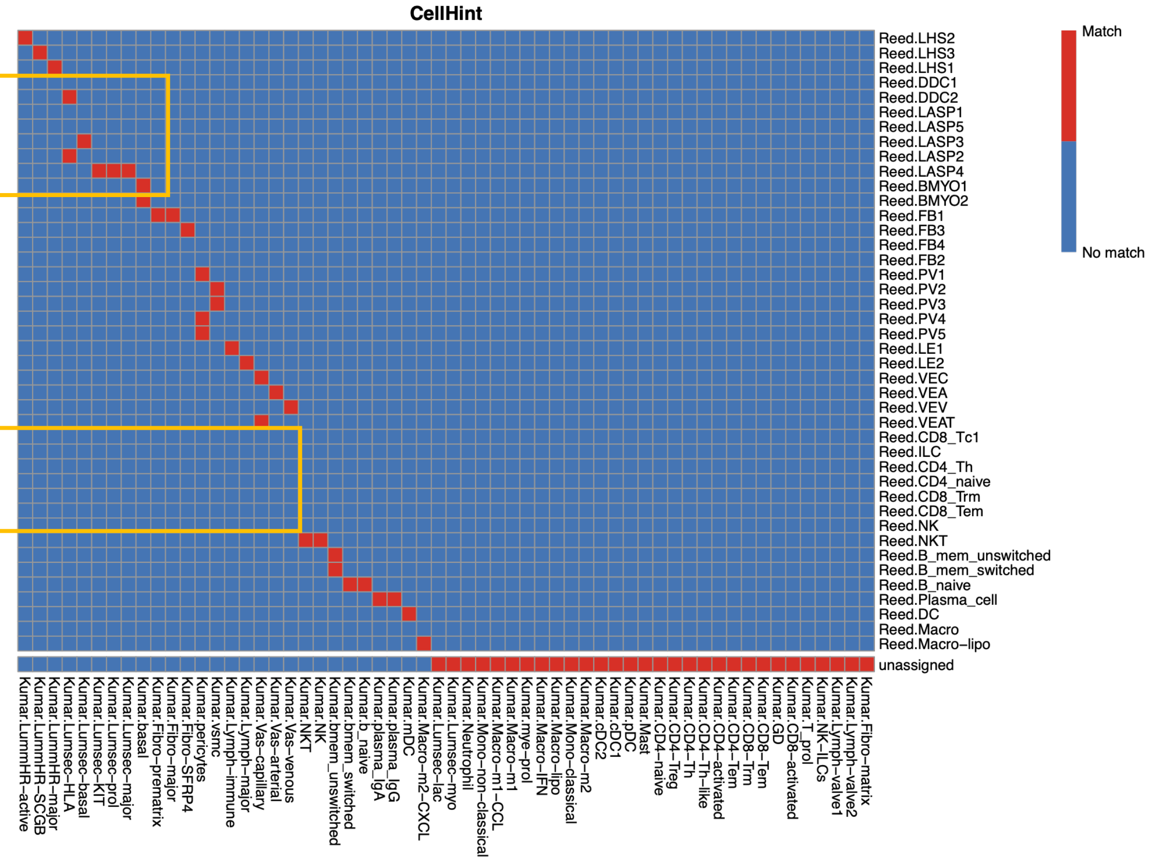

### Supplementary Figure 15

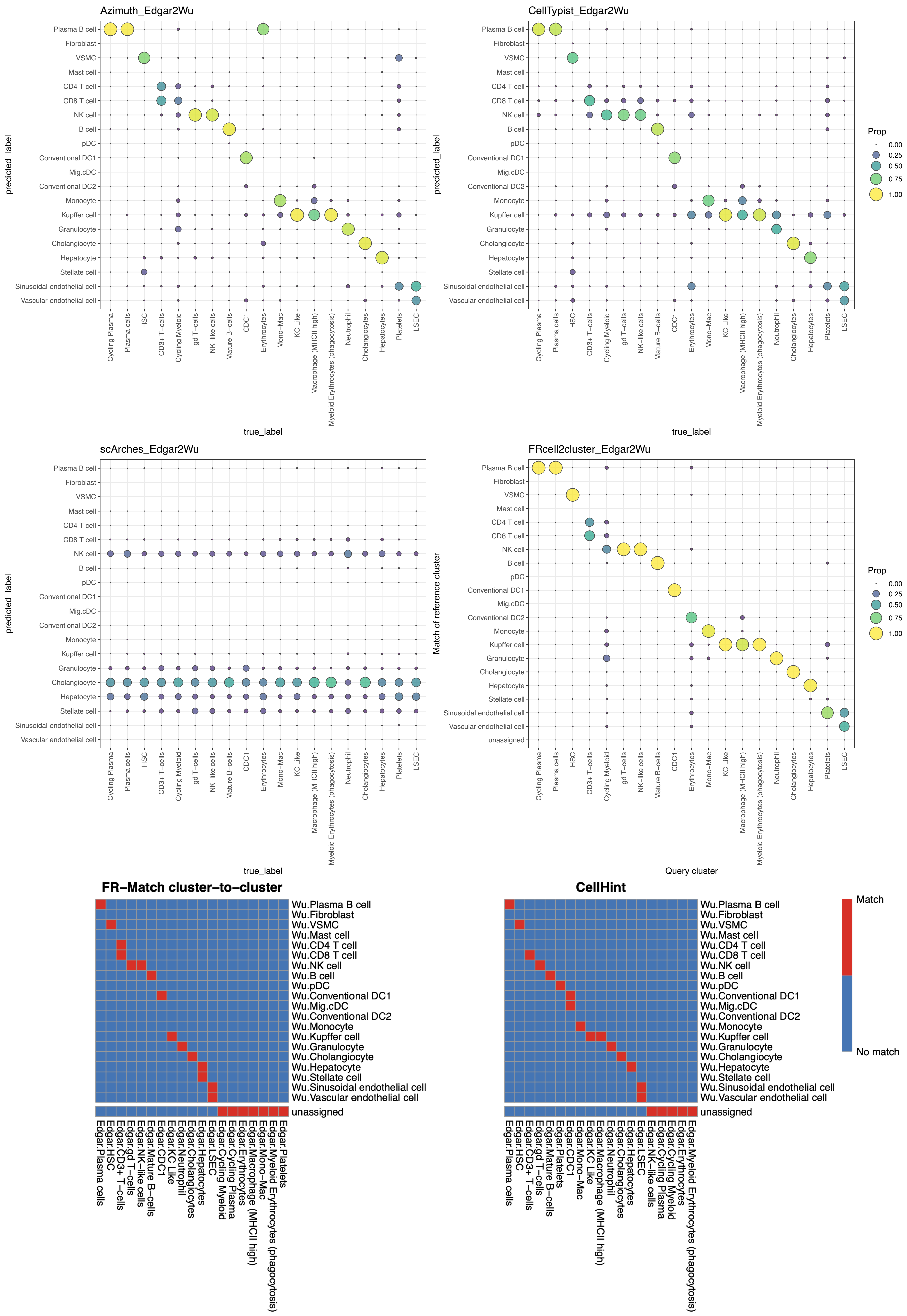

### Supplementary Figure 16

#### Gut

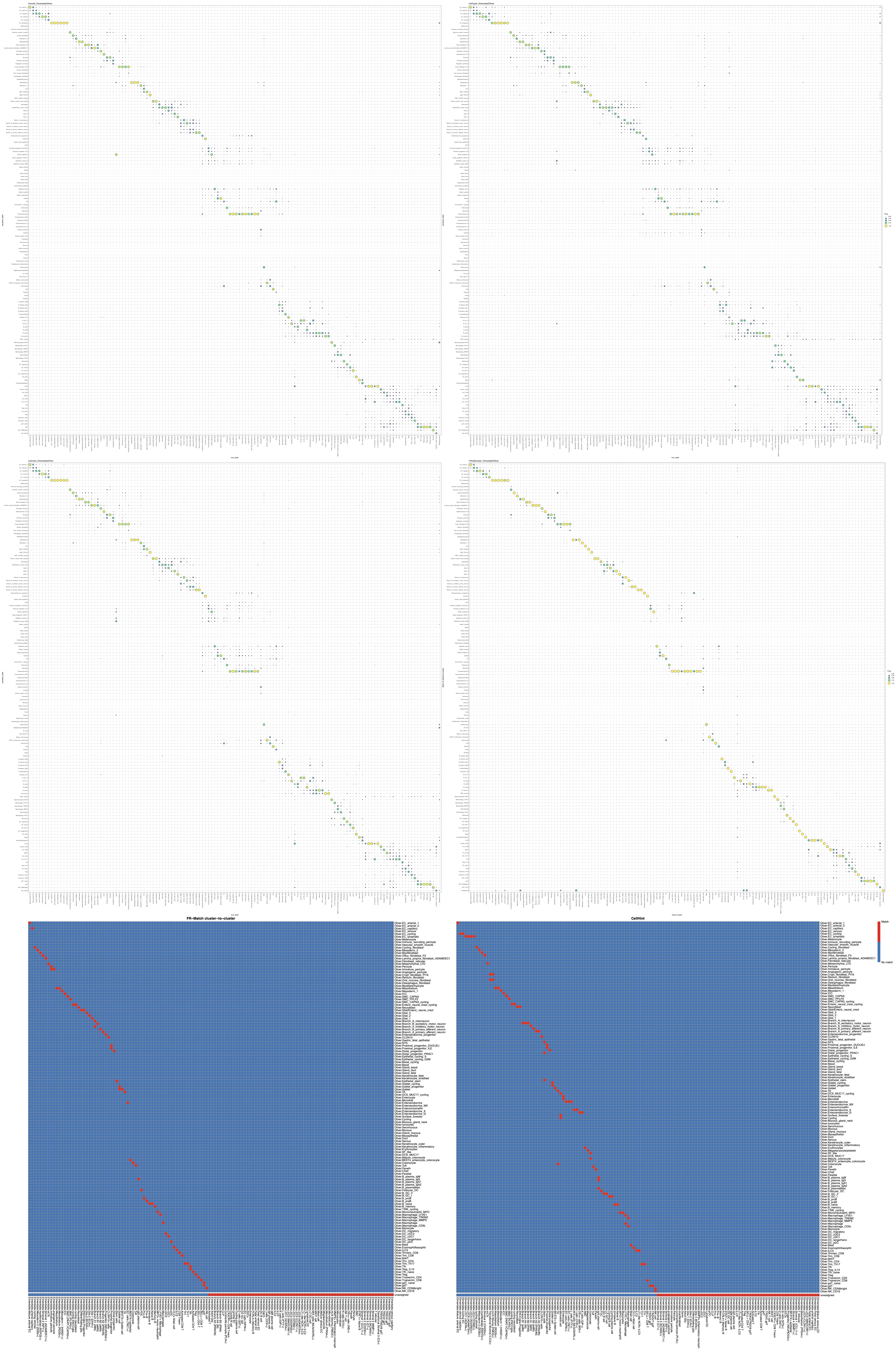

### Supplementary Figure 17

#### Lung (non-small cell lung cancer)

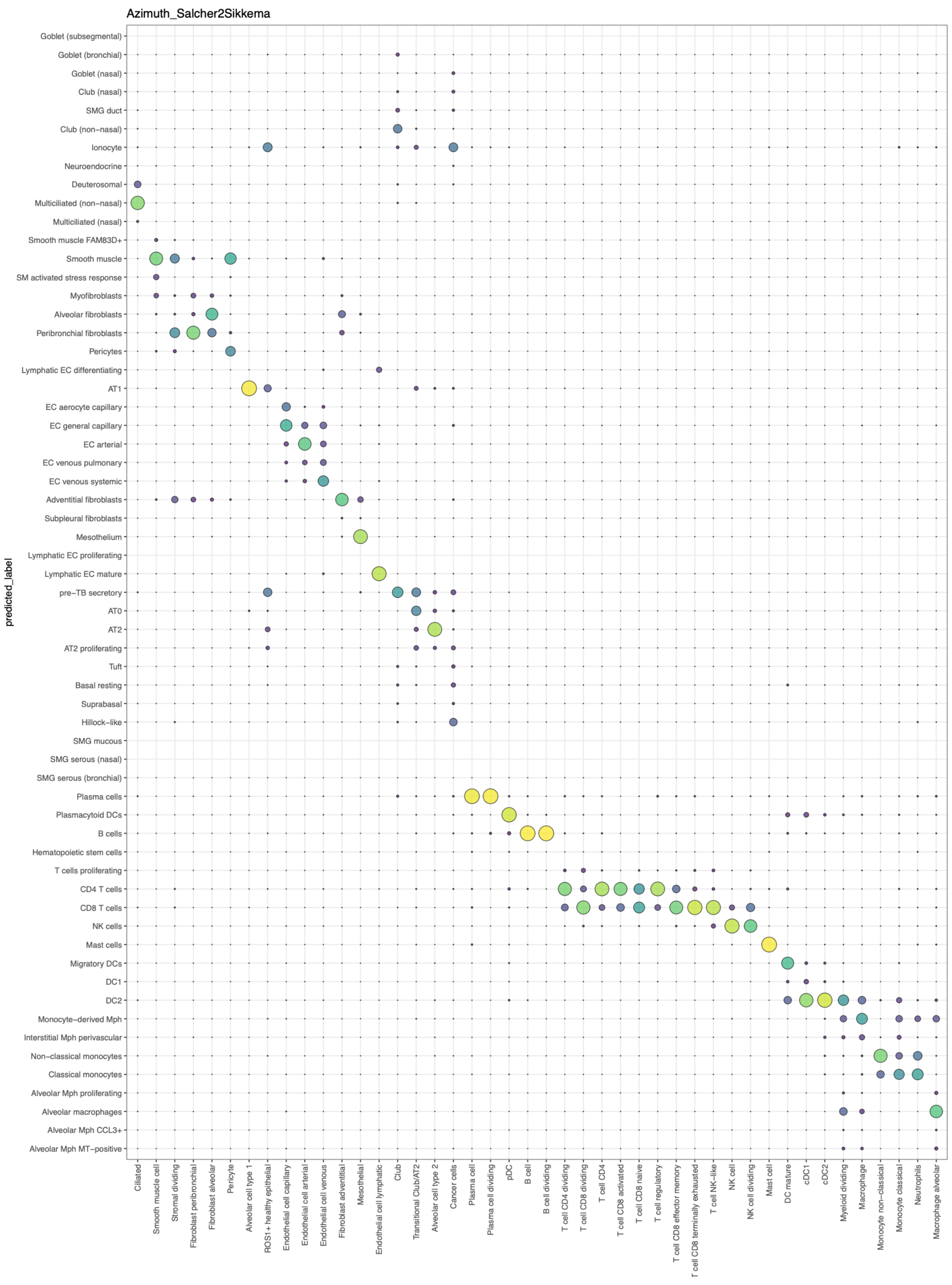


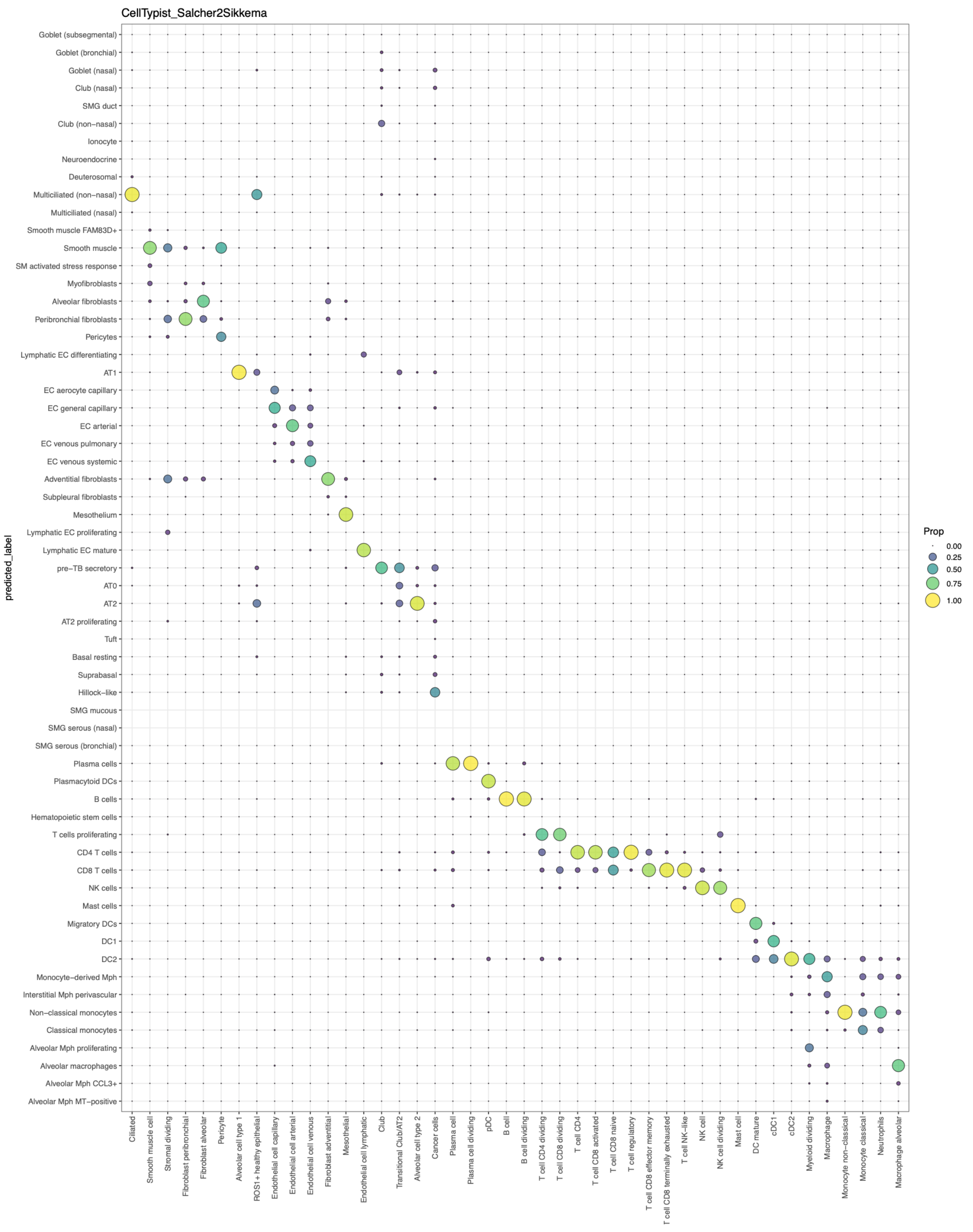


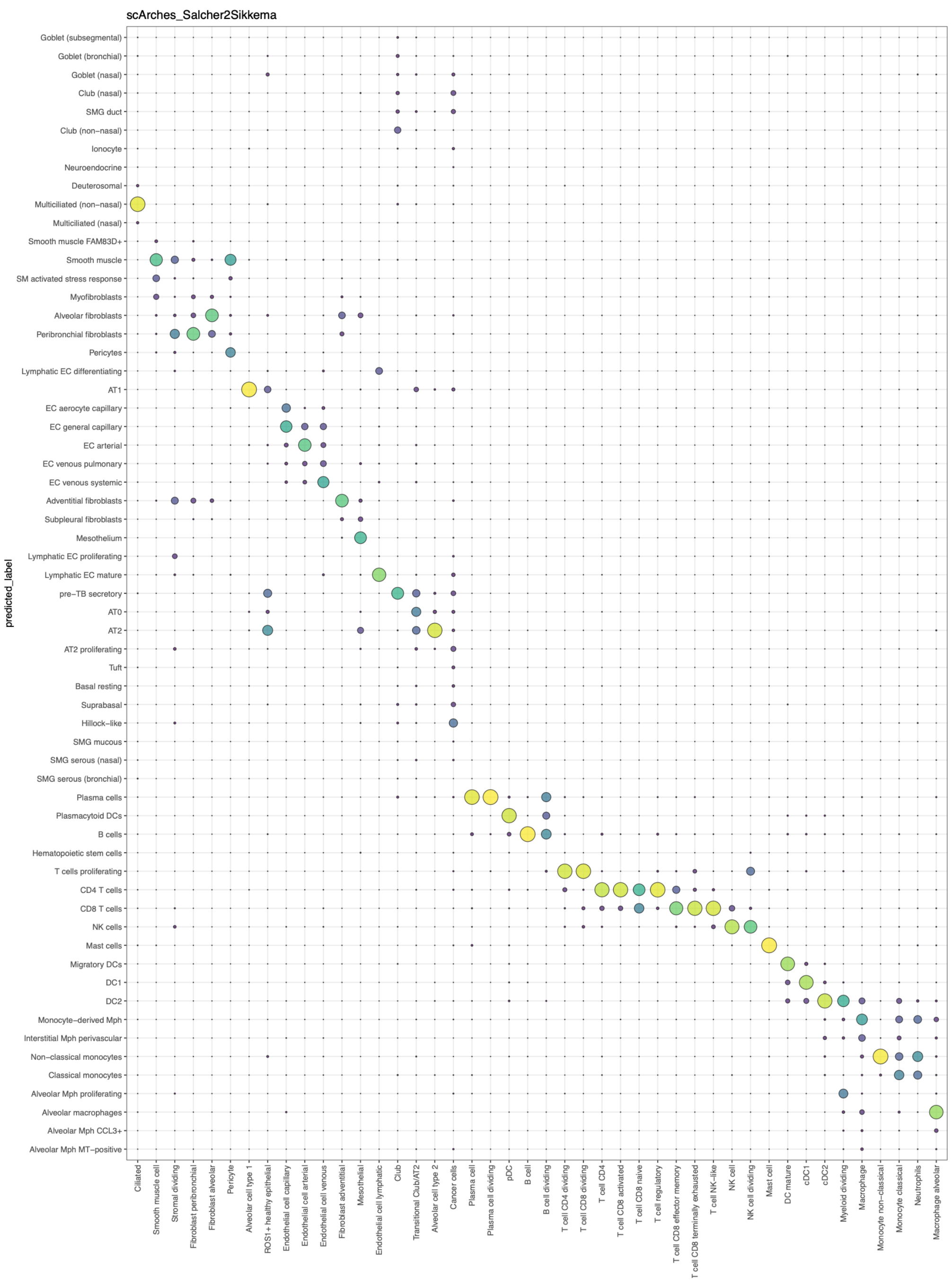


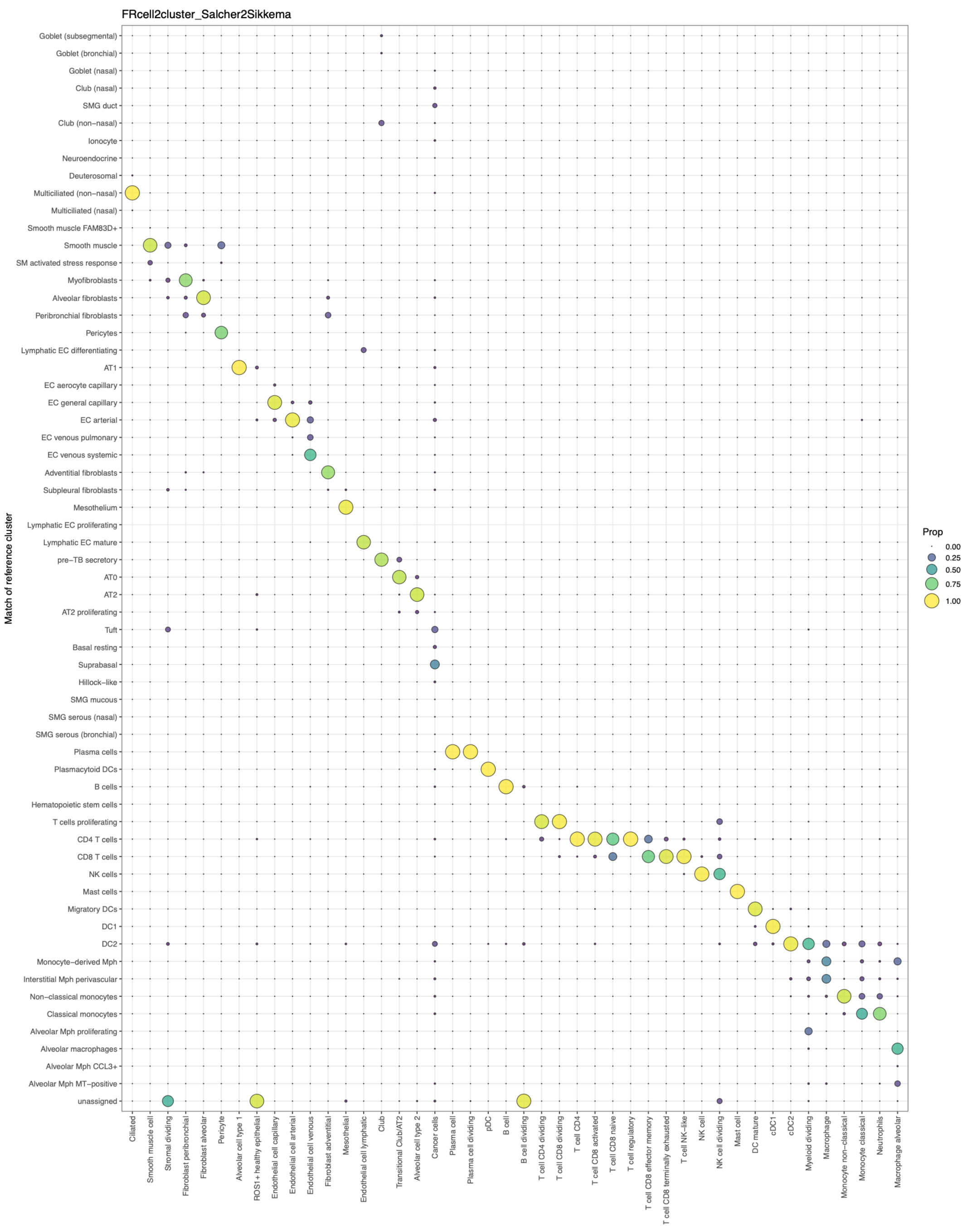


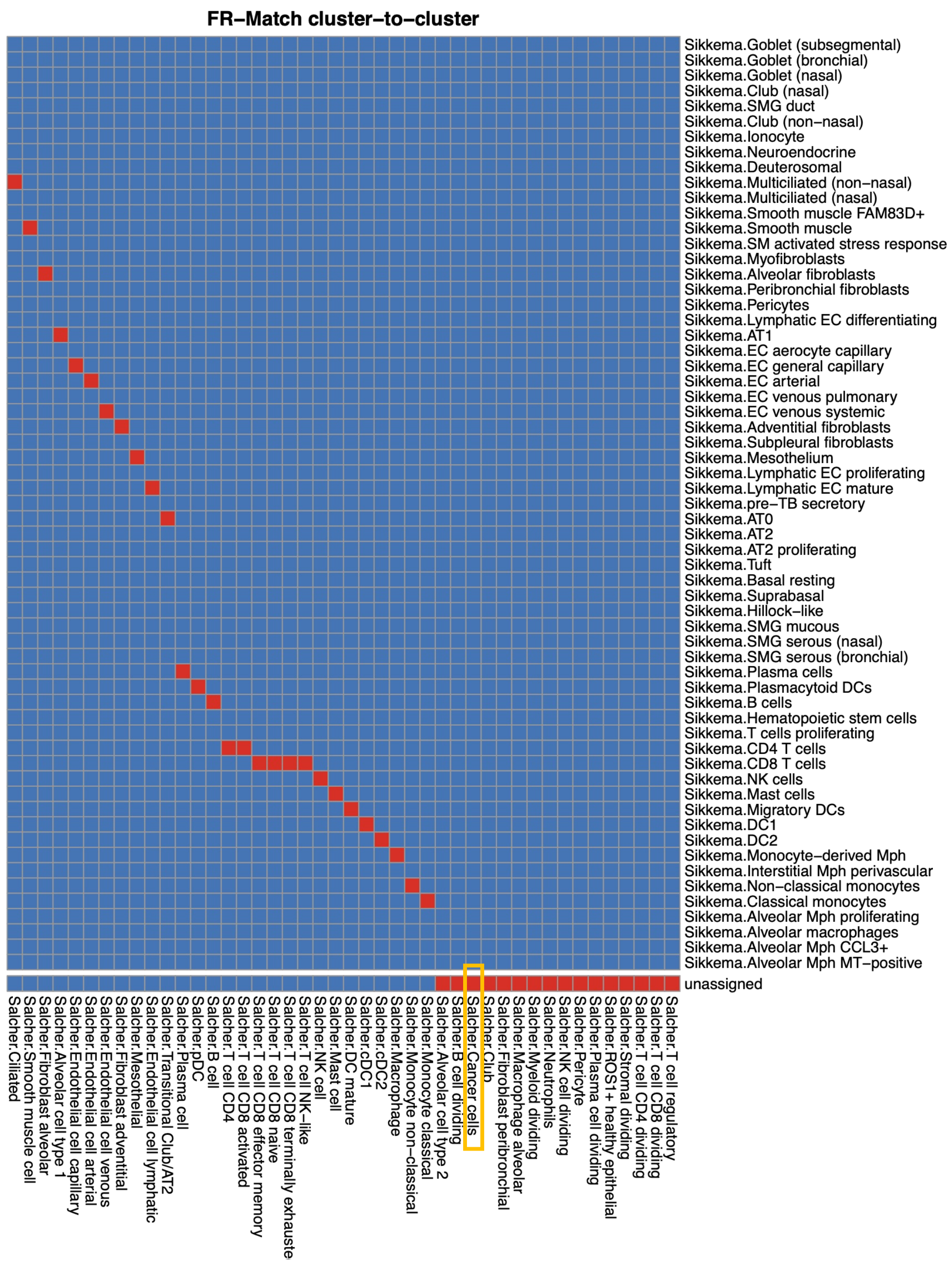


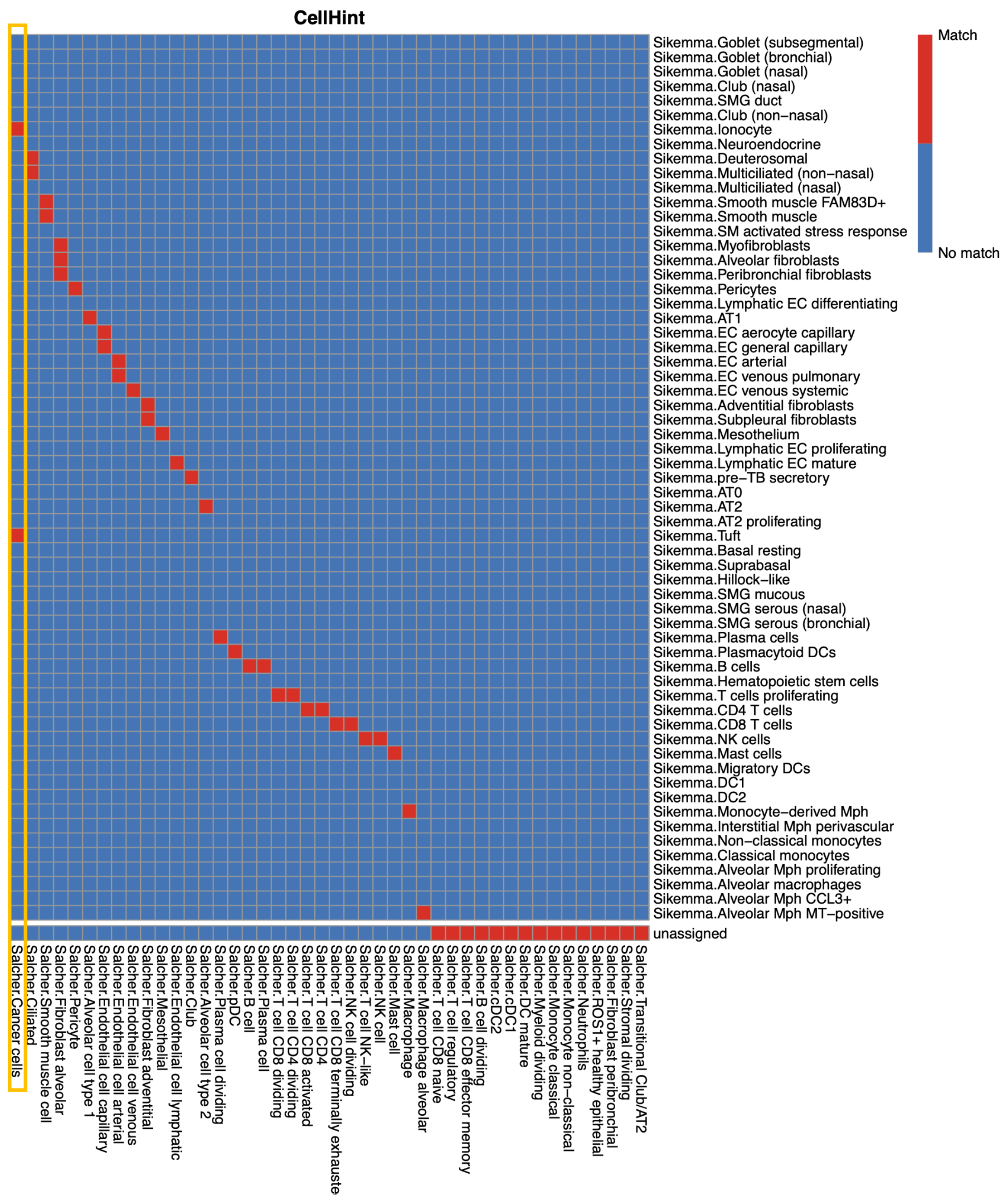

### Supplementary Figure 18

#### Fetal immune

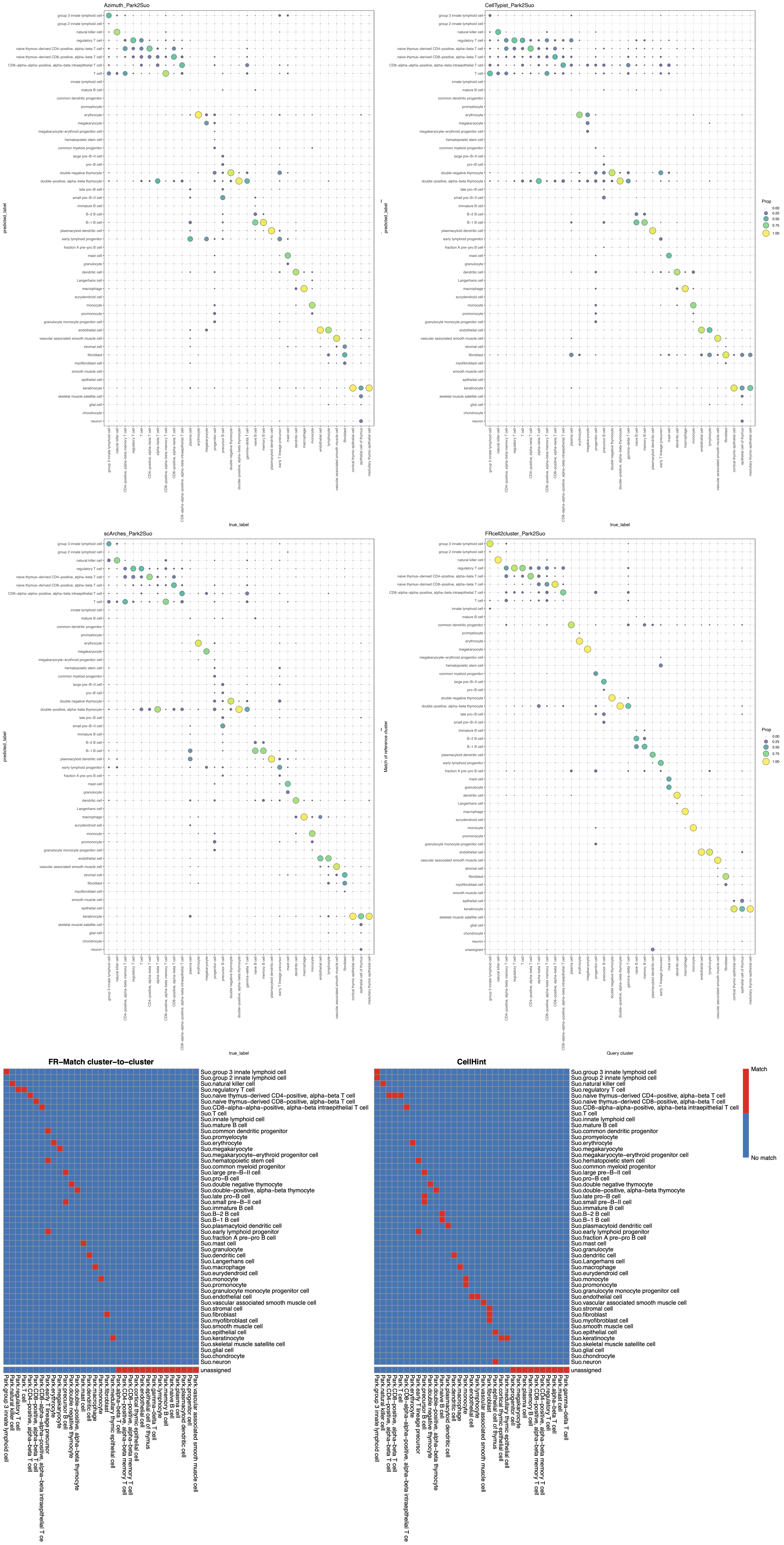

### Supplementary Figure 19

#### Mouse skeleton

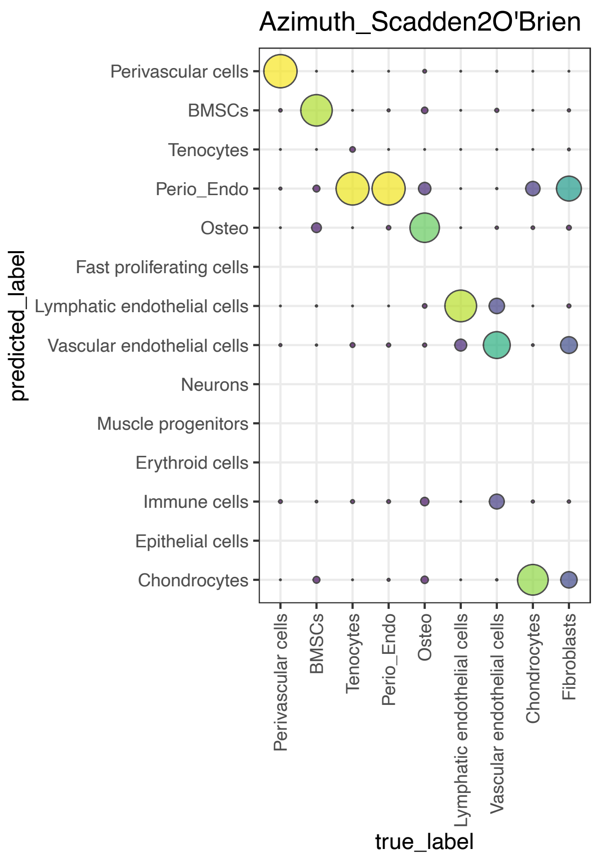


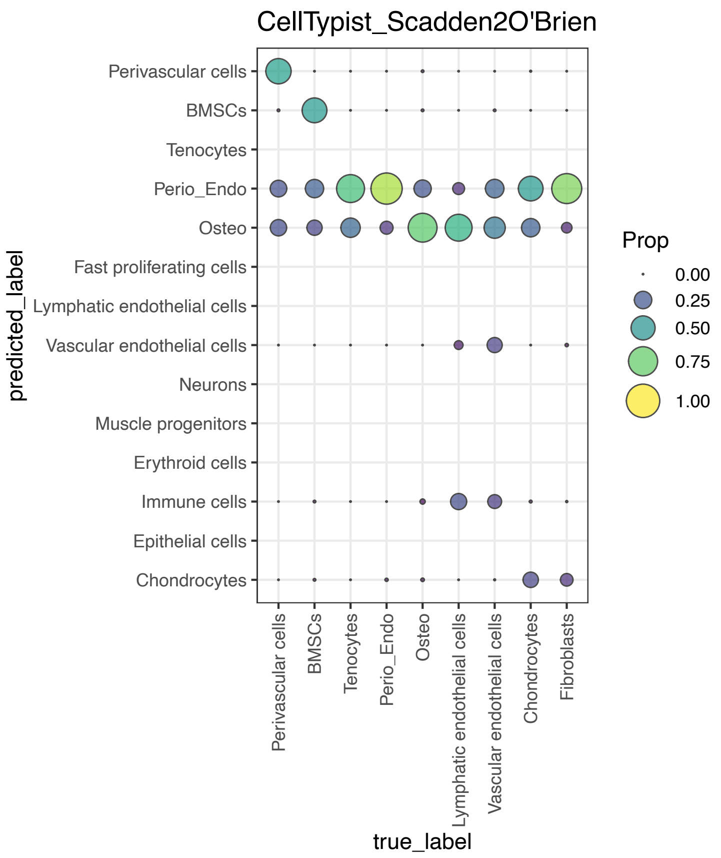


*No count data*

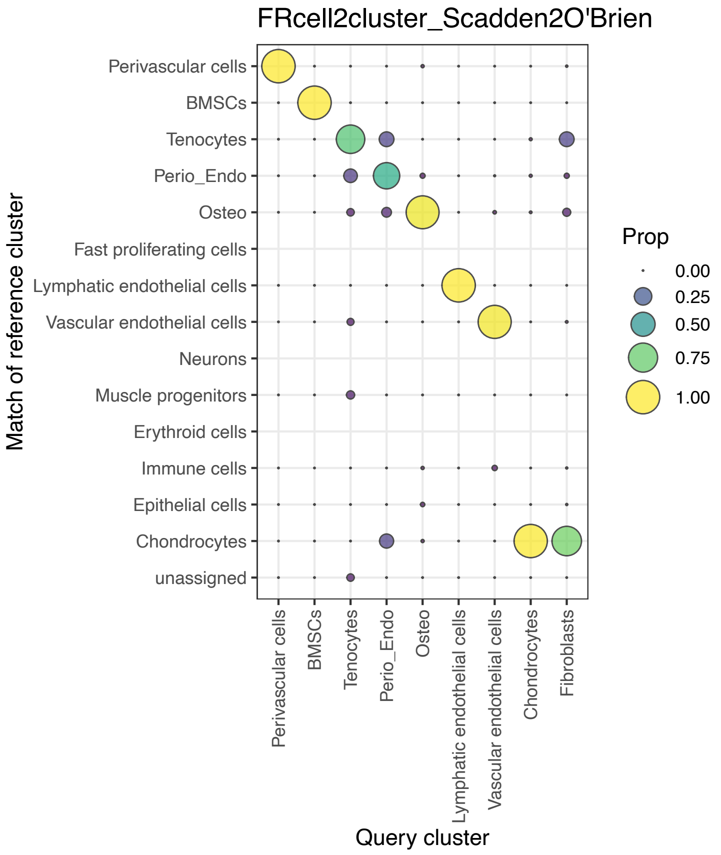


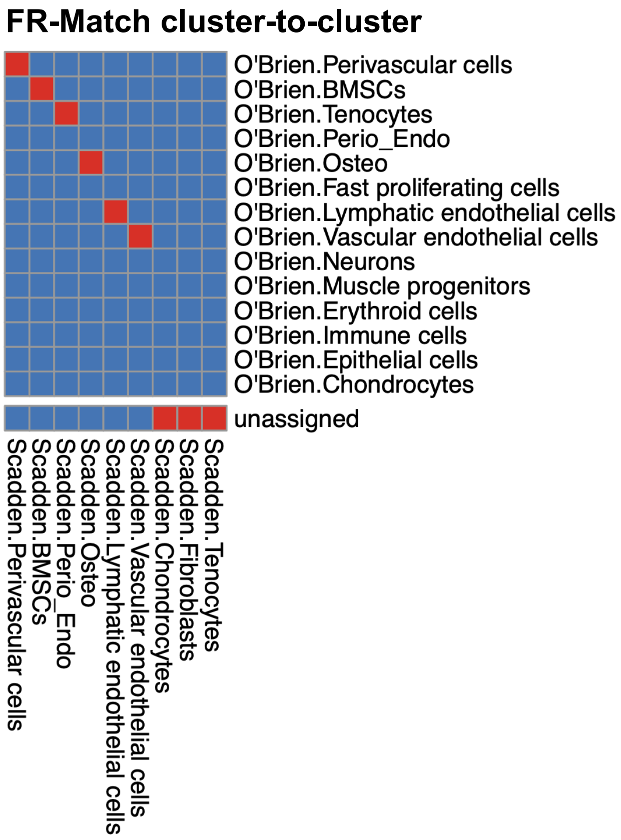


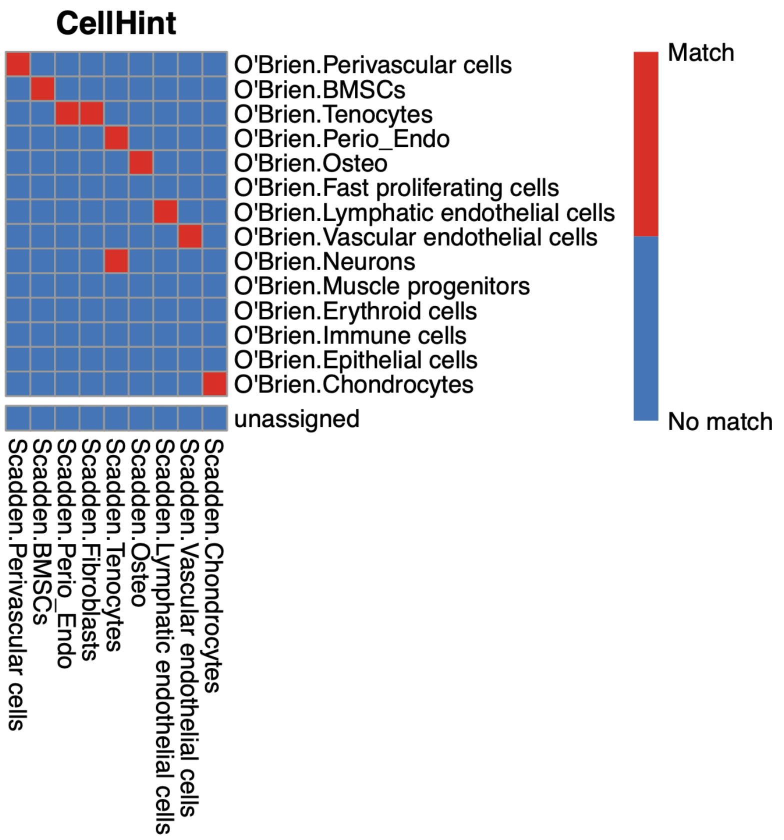

### Supplementary Figure 20

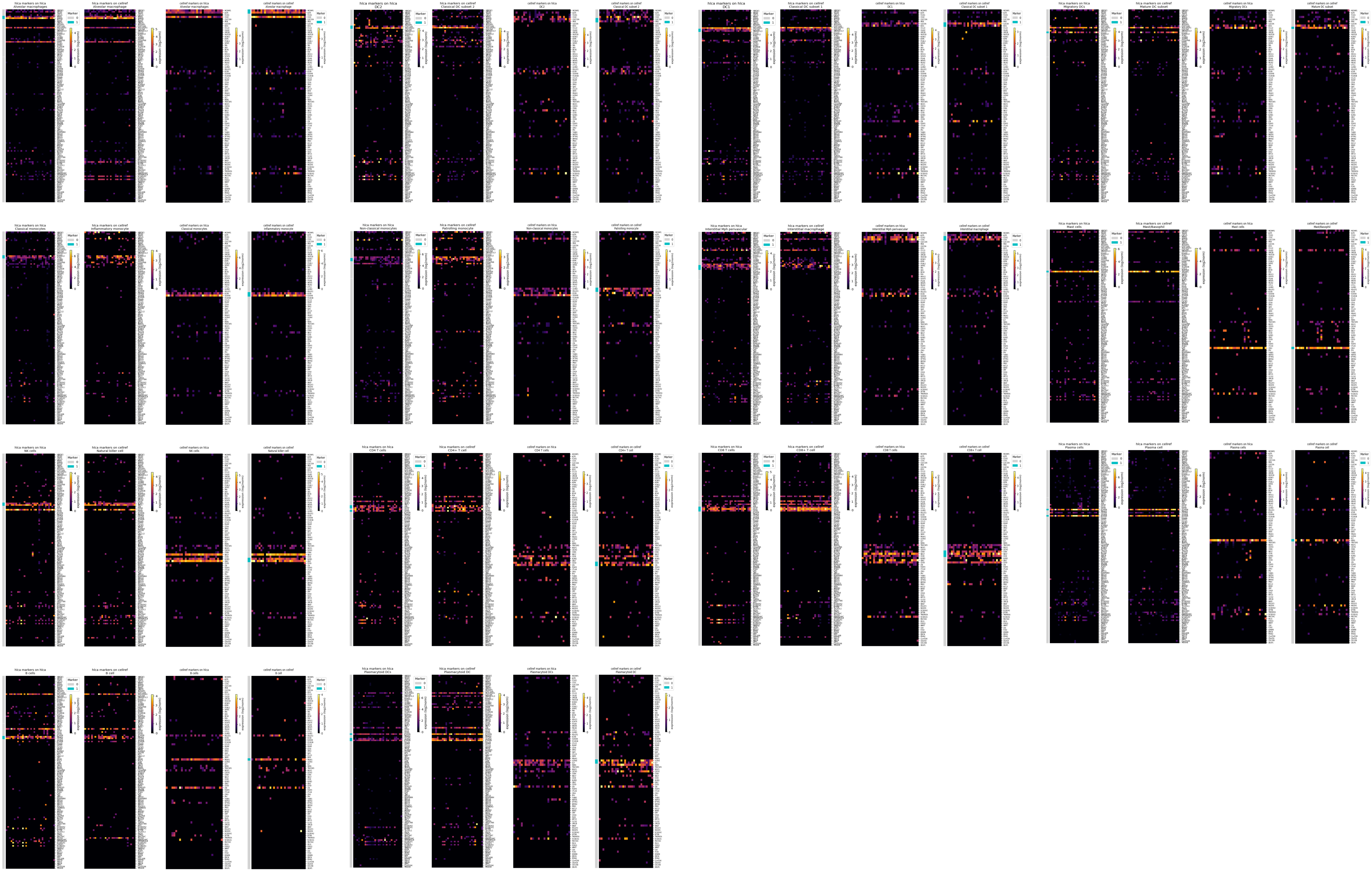

### Supplementary Figure 21

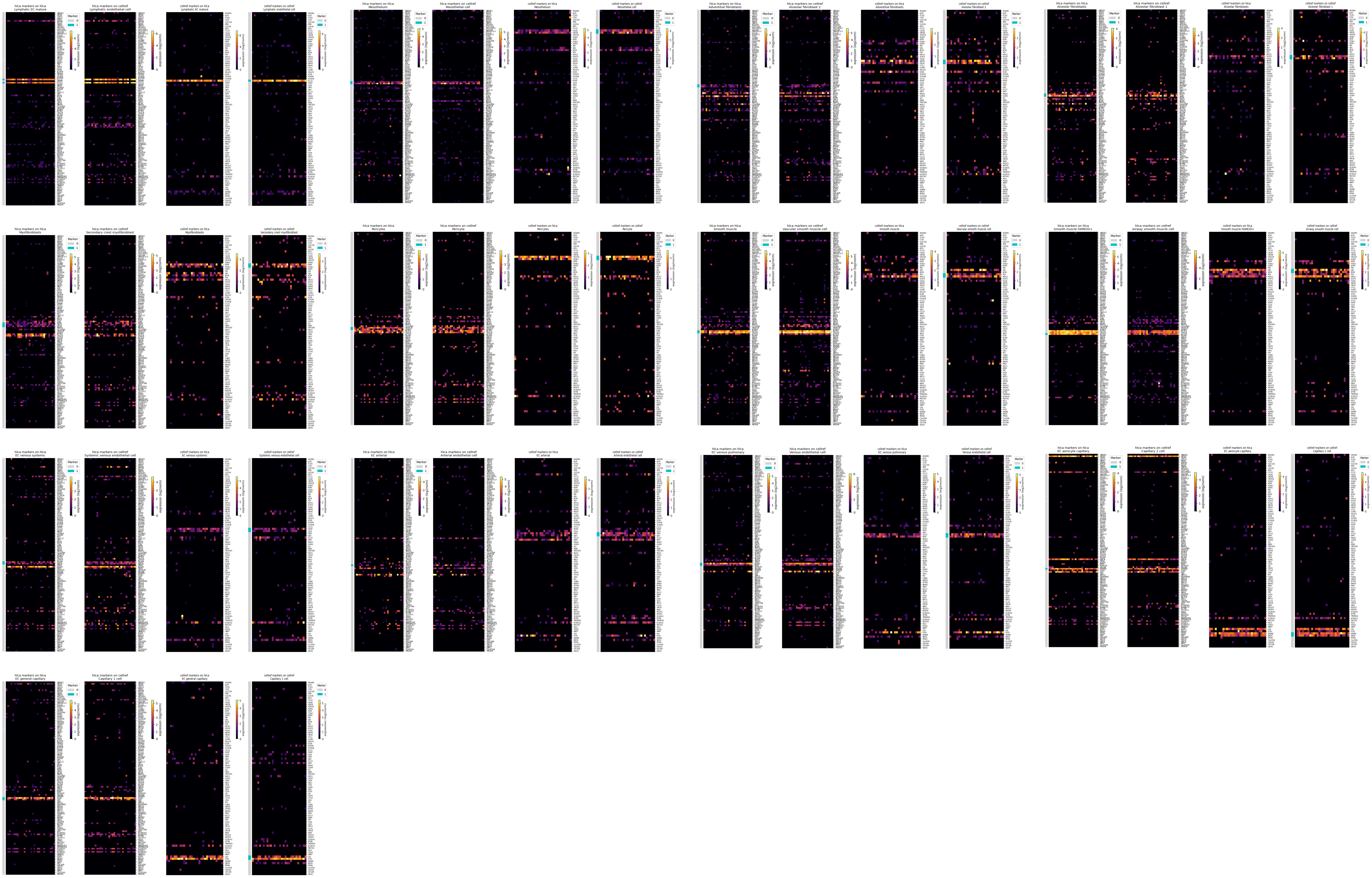

### Supplementary Figure 22

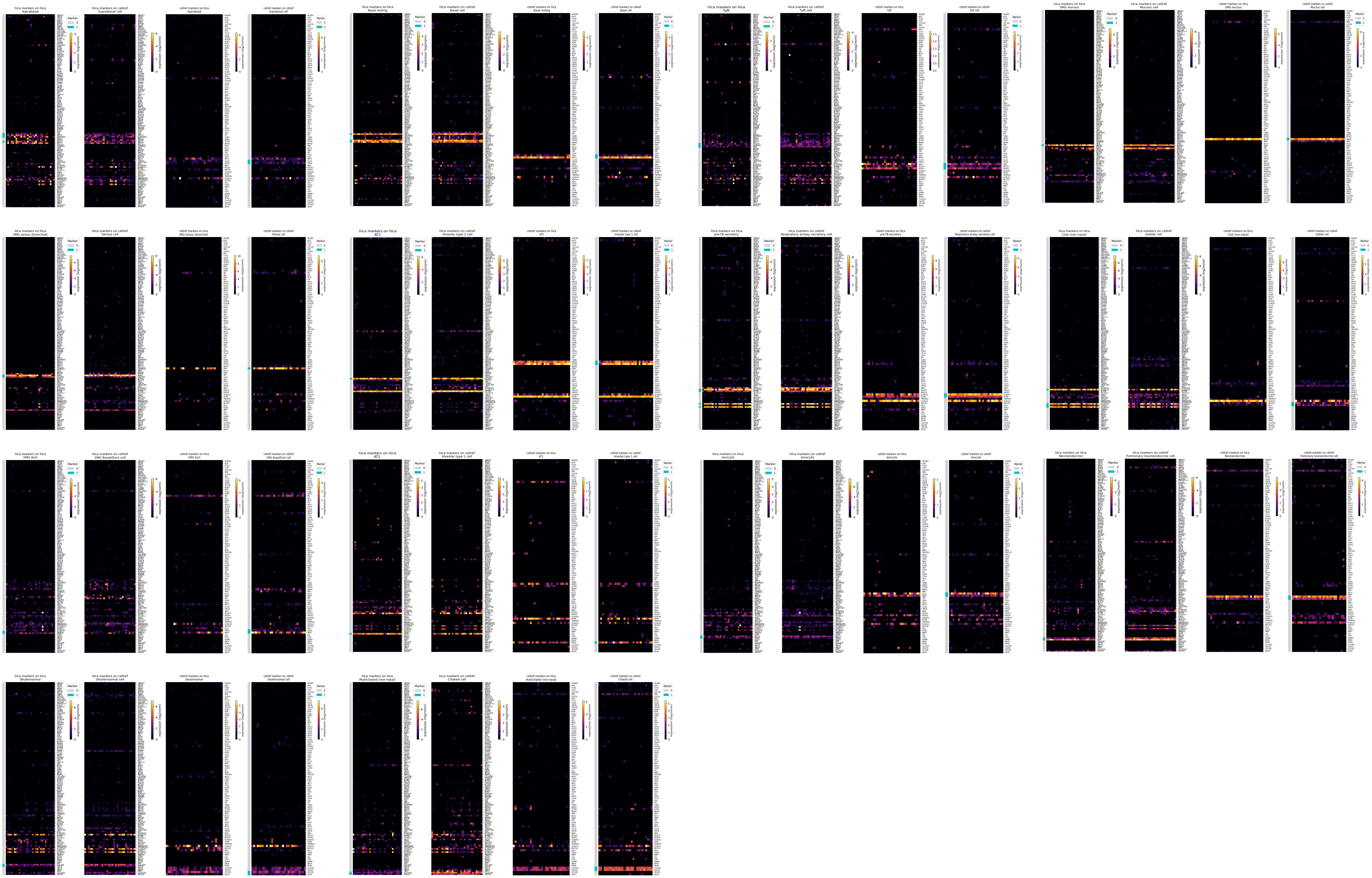

### Supplementary Figure 23

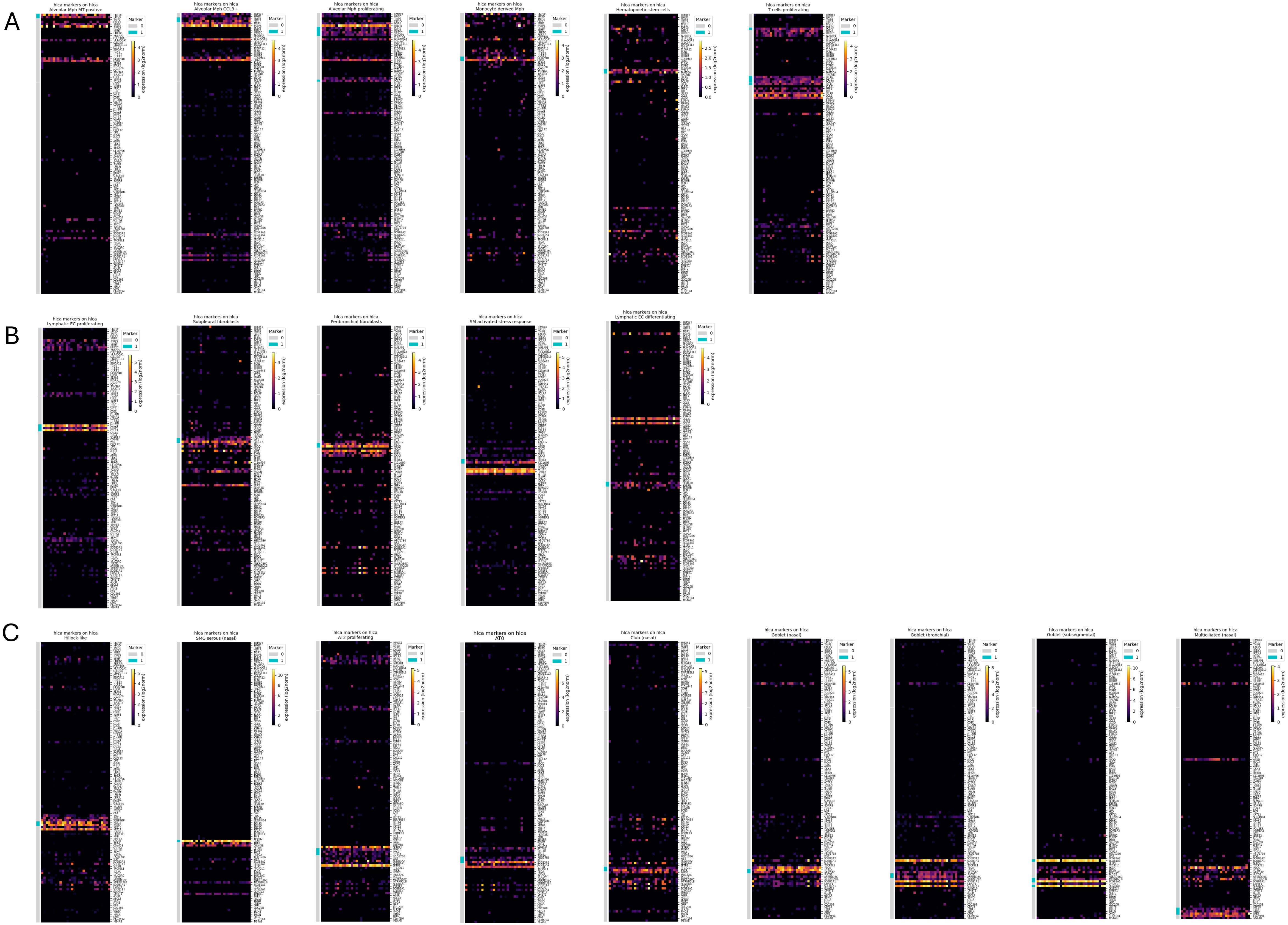

### Supplementary Figure 24

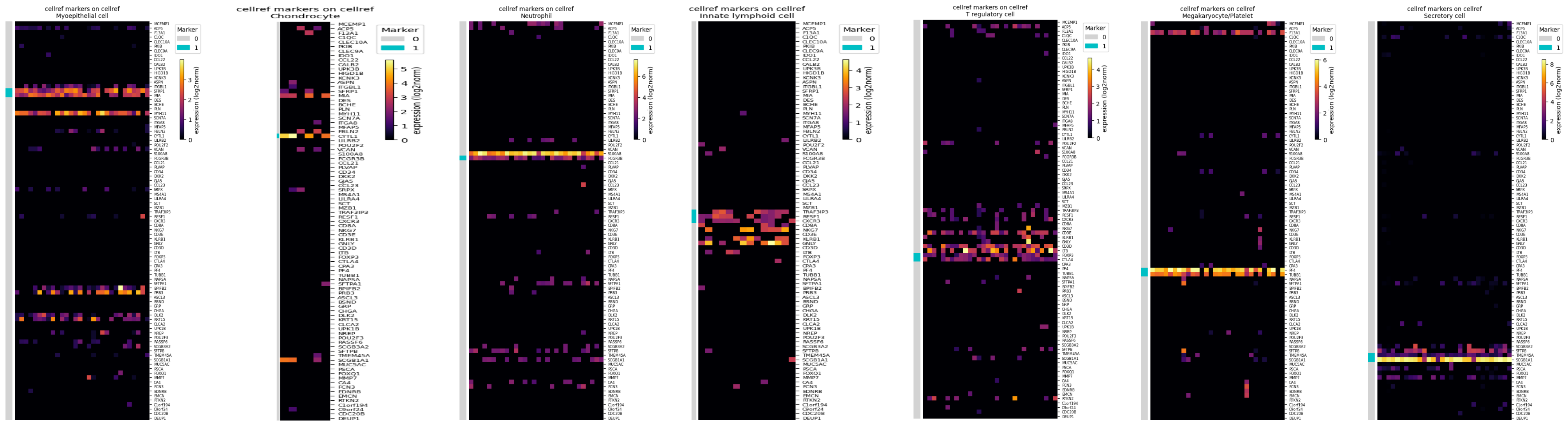
