## Supplementary Figure Legends for "Benchmarking single cell transcriptome matching methods for incremental growth of cell atlases"

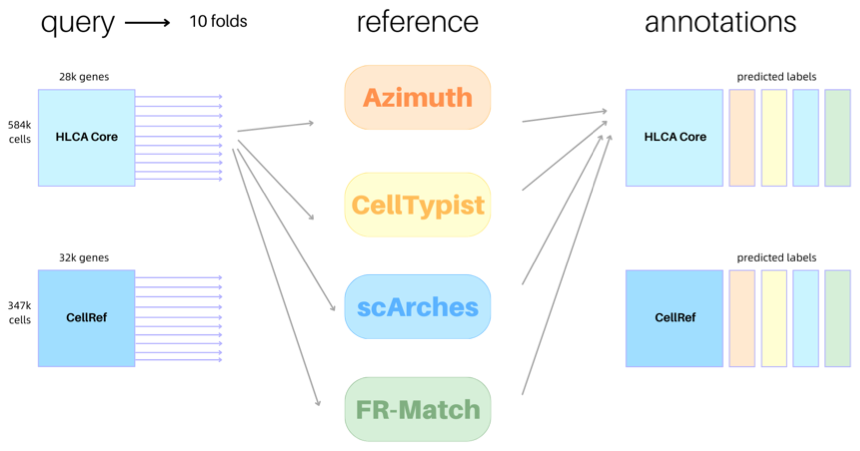
**Supplementary Figure 1. The benchmarking scheme follows a 10-fold cross-validation set up.** Similar scheme is used in the CellRef matching for computational efficiency.

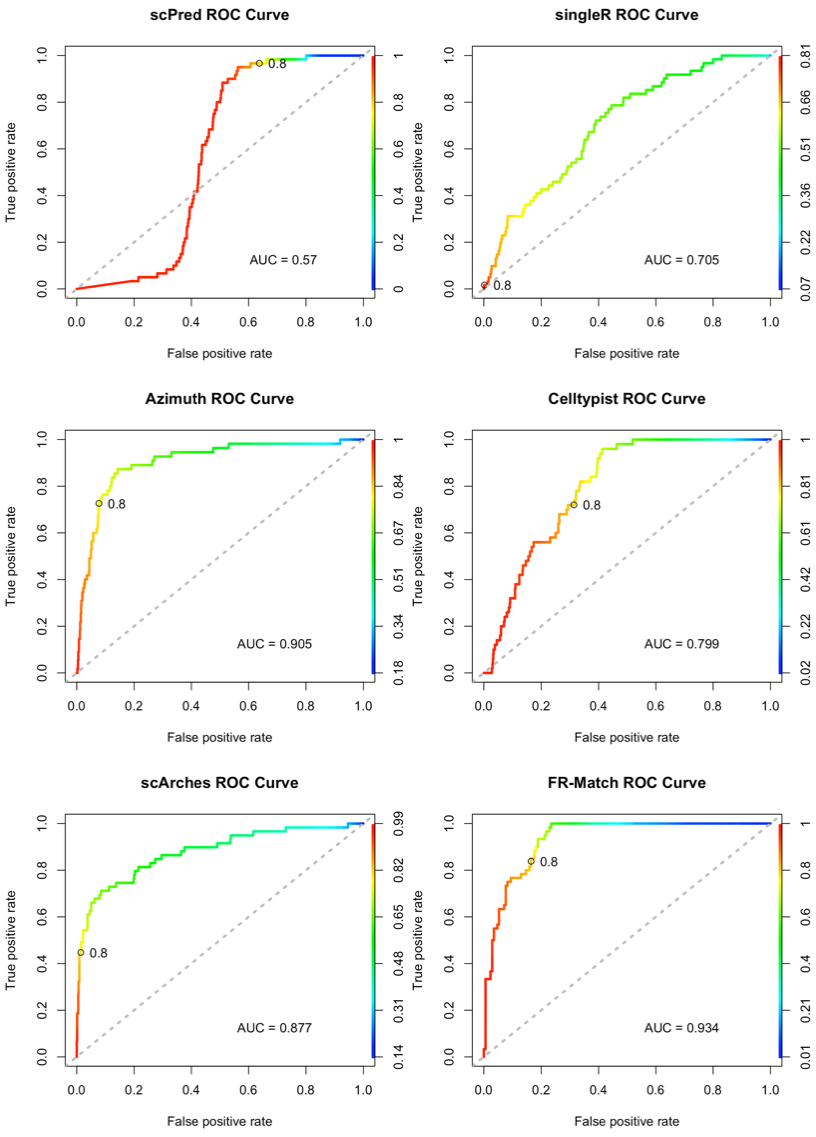

**Supplementary Figure 2. Colorized ROC curve for each method in the benchmarking results.** The plot is generated using the plotting function with “colorize=TRUE” from the ROCR R package.

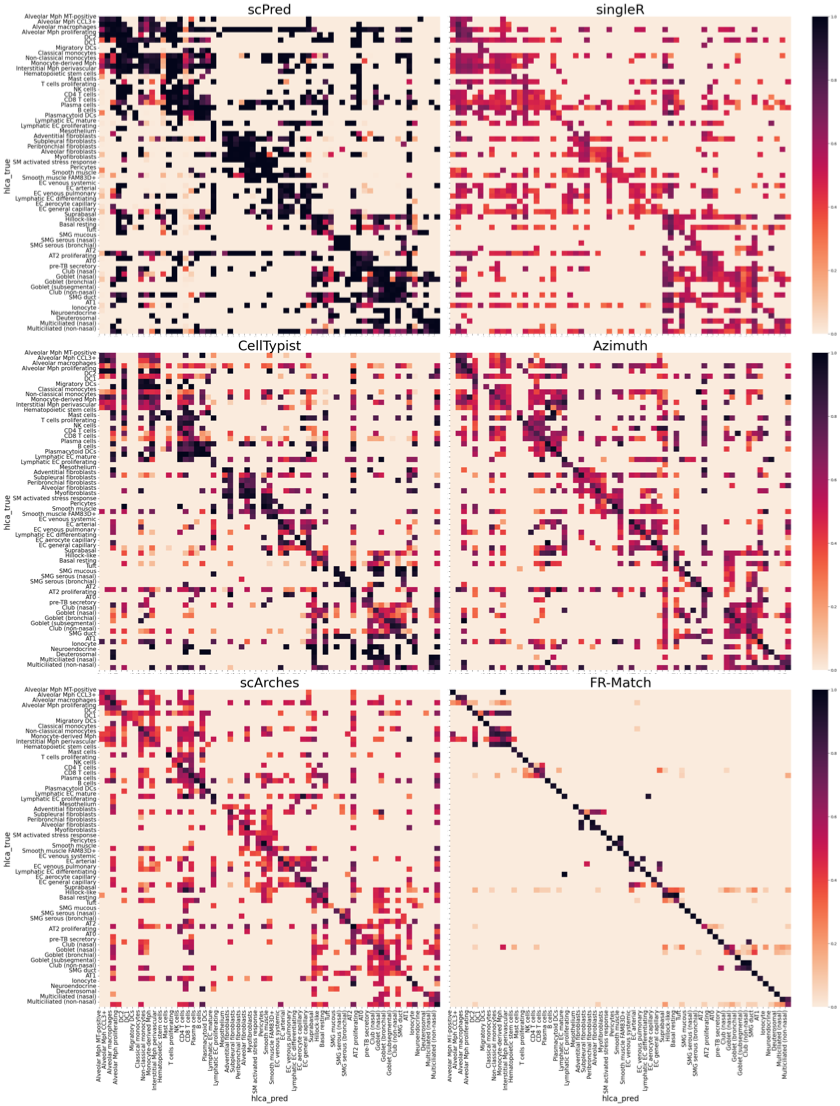

**Supplementary Figure 3. Heatmaps of the average confidence scores for each true-predicted pair for each method in the cross-validation benchmarking results.** Clusters ordered by similarity from the HLCA taxonomy dendrogram (see **Methods**). The heatmaps further elaborate on the results of the colorized ROC curves (**Supplementary Figure 2**) by depicting the mean confidence for each true label and predicted label pairing generated by the corresponding method. A perfect heatmap would display a clean distinct diagonal line, signifying a perfect alignment between the prediction and true labels at a high confidence level (dark color). Darker squares in the heatmap correspond to the red portion of the colorized ROC curve; lighter regions correspond to the blue portion.

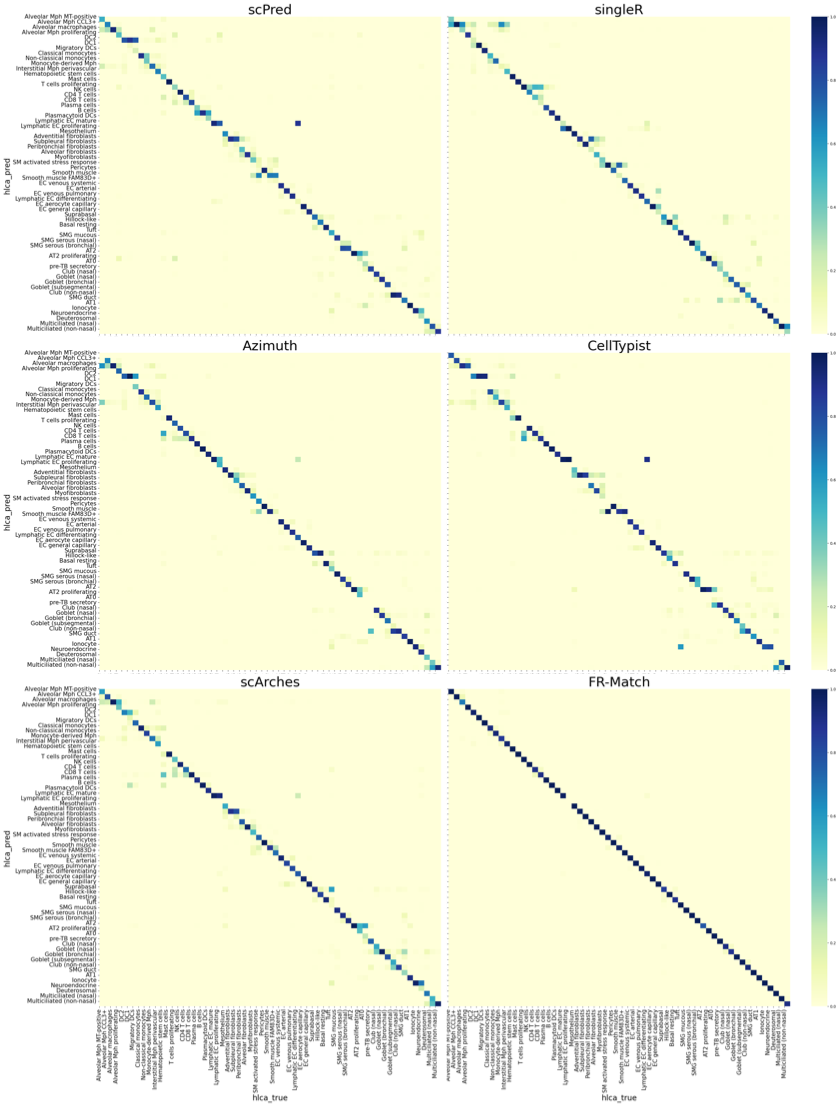
 **Supplementary Figure 4. Heatmaps of cell type-to-cell type confusion matrix in the HLCA cross-validation results.**

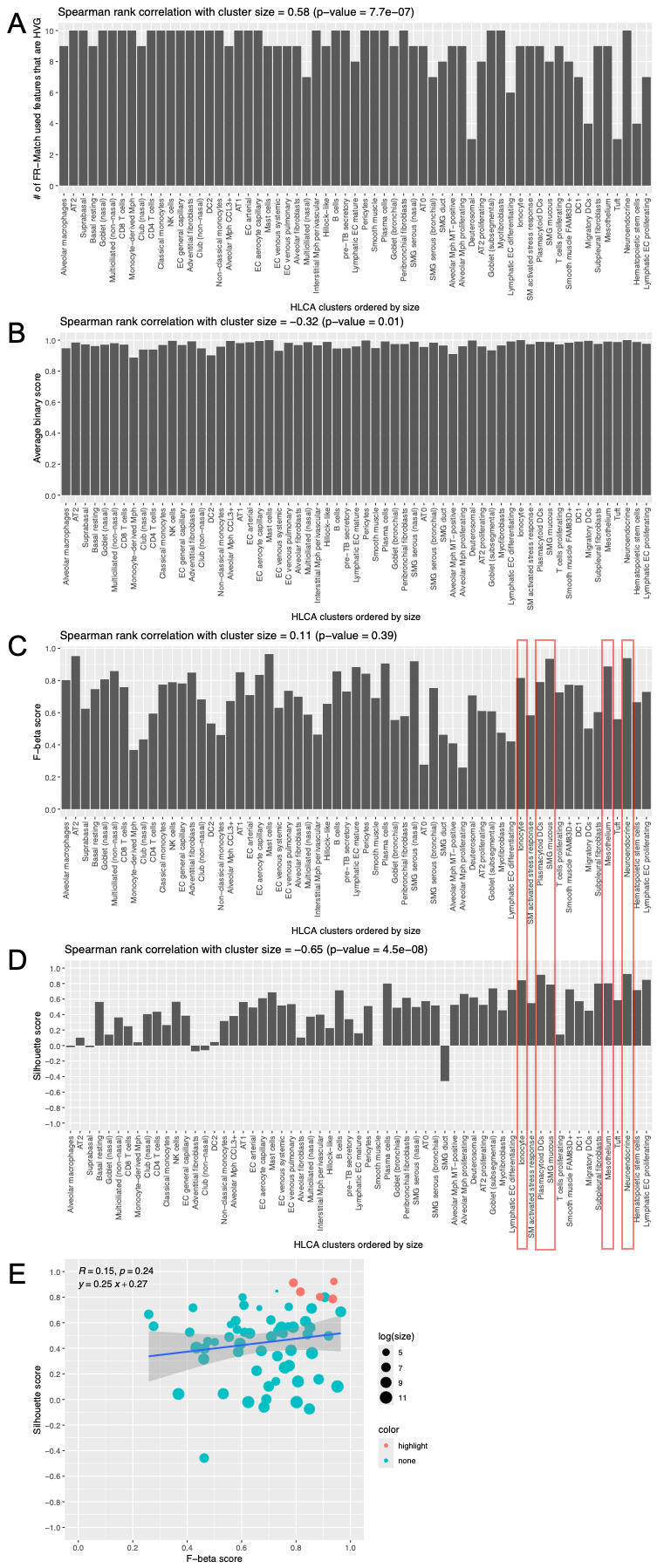

**Supplementary Figure 5. Per-cluster analysis of feature spaces and cluster quality metrics of the HLCA clusters. (A-D)** HLCA clusters are ordered by their cluster sizes. Quantity plotted in the bar plot is shown in the y-axis title. Correlation with cluster size and correlation test p-value are shown in the plot title. **(E)** Scatter plot of F-beta score (C) and silhouette score (D). Red dots correspond to highlighted rare cell types in (C) and (D). Dot size reflects cluster size.

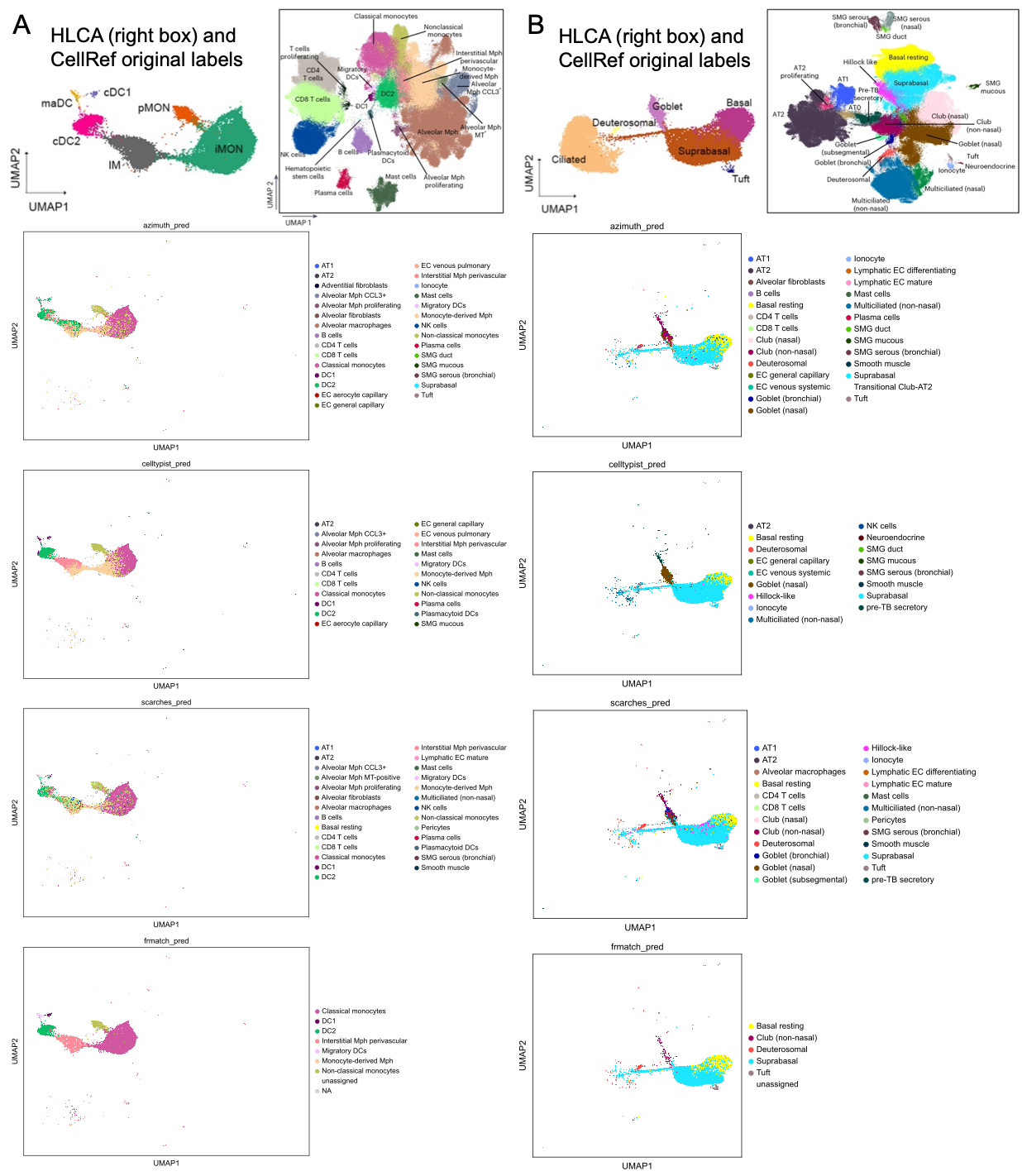

**Supplementary Figure 6. Detailed UMAPs of the “IM” region in (A) and “basal” region in (B) with all predicted labels printed out.** “Unassigned” is shown as white color, implying a clean-up of the low confidence cells (**Supplementary Figure 5**). The HLCA and CellRef original labels in the original UMAPs are shown on the top.

**Supplementary Figure 7. UMAPs deciphering more differences of the cell-based methods. (A)** CellRef cells in the UMAPs are colored by the prediction confidence scores by each method. **(B)** The Azimuth-only labels (due to versioning of the HLCA annotations) and the FR-Match “unassigned” label are denoted in the UMAPs. Black arrows indicate examples where isolated clusters are both having low scores and “unassigned” in FR-Match.

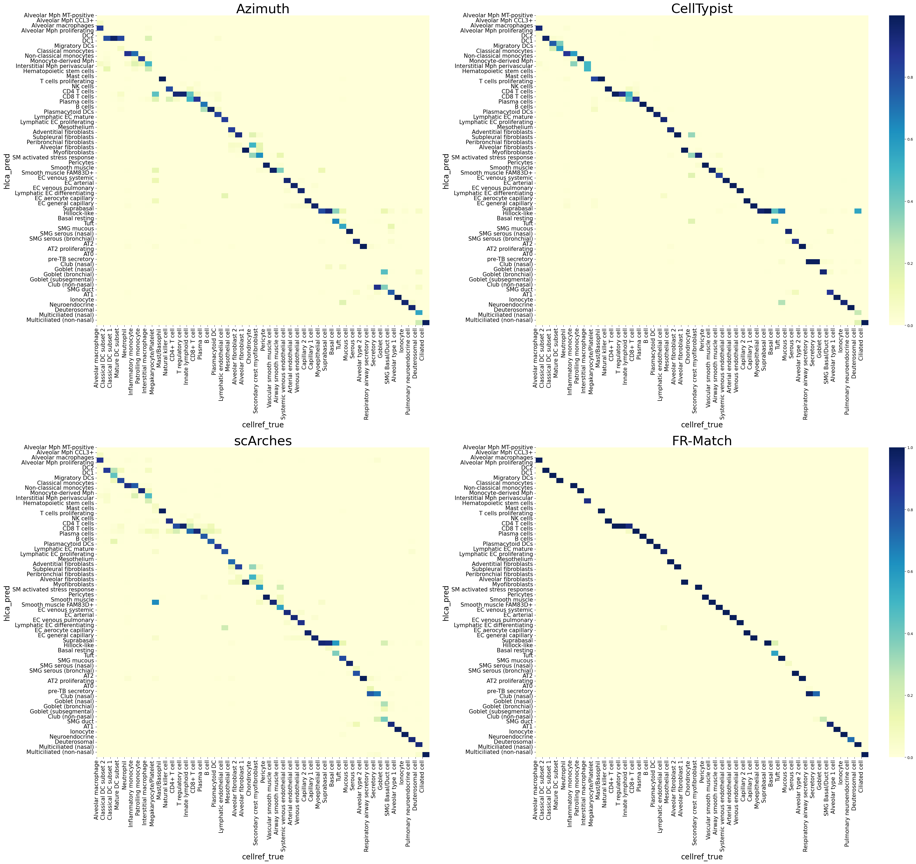
**Supplementary Figure 8. Heatmaps of cell type-to-cell type confusion matrix in the CellRef and HLCA matching results.**

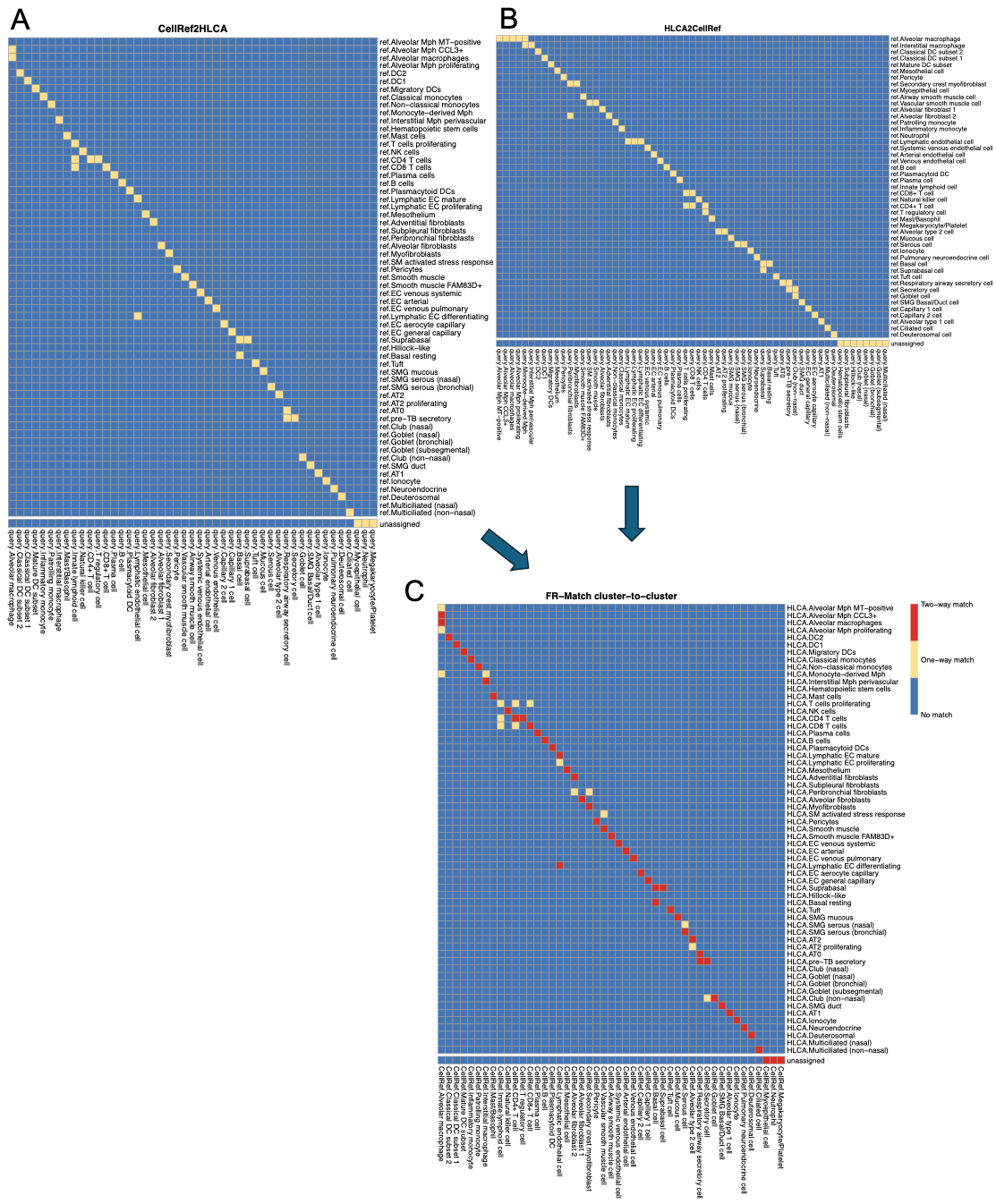

**Supplementary Figure 9. FR-Match cluster-to-cluster results. (A)** CellRef 🡪 HLCA matching results. **(B)** HLCA 🡪 CellRef matching results. **(C)** Combined one-way and two-way matches from both directions.
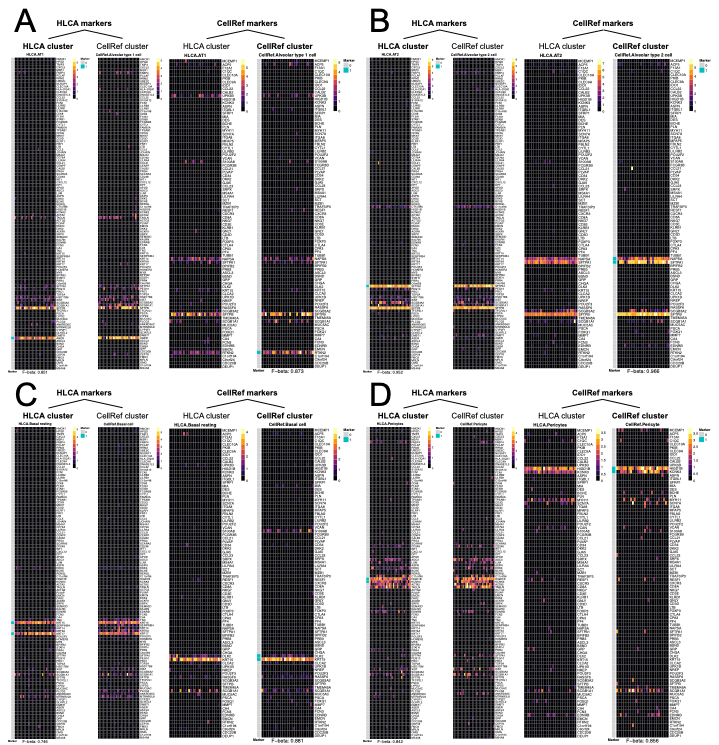

**Supplementary Figure 10. Example “barcode” plots of matched lung cell types. (A)** AT1 in HLCA is matched with alveolar type 1 cell in CellRef. **(B)** AT2 in HLCA is matched with alveolar type 2 cell in CellRef. **(C)** Basal resting in HLCA is matched with basal cell in CellRef. **(D)** Pericytes in HLCA is matched with pericyte in CellRef.

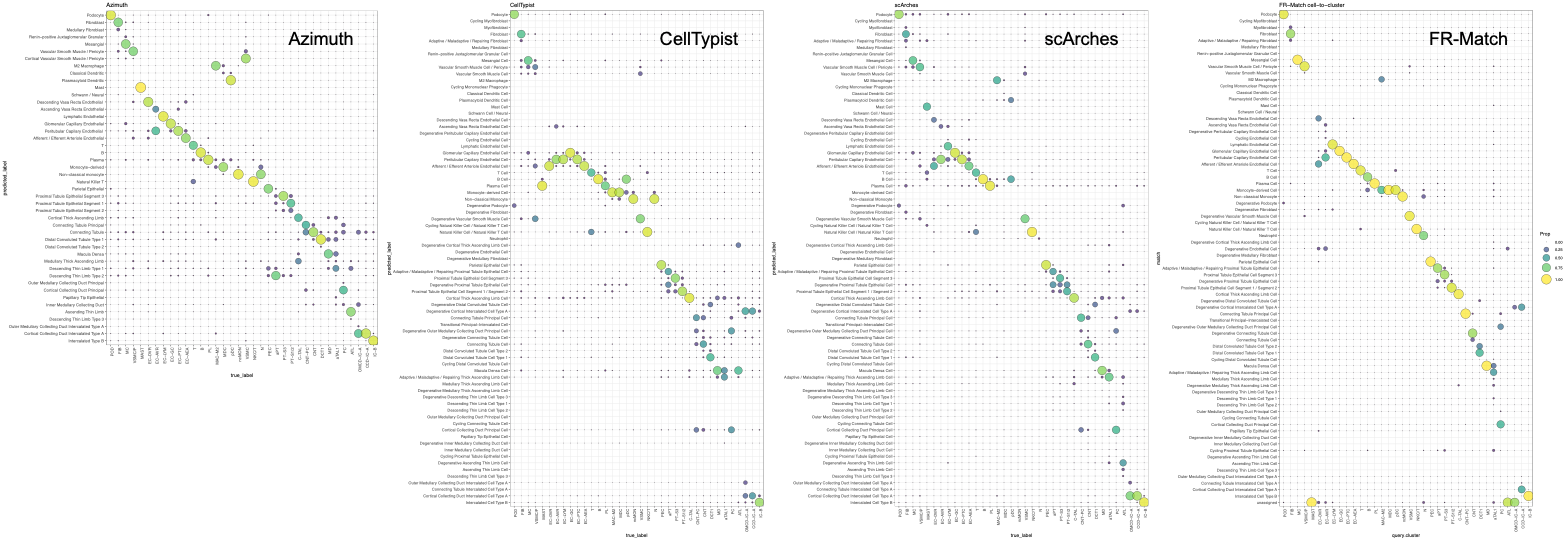

**Supplementary Figure 11. Cell-based matching results for the kidney analysis.**

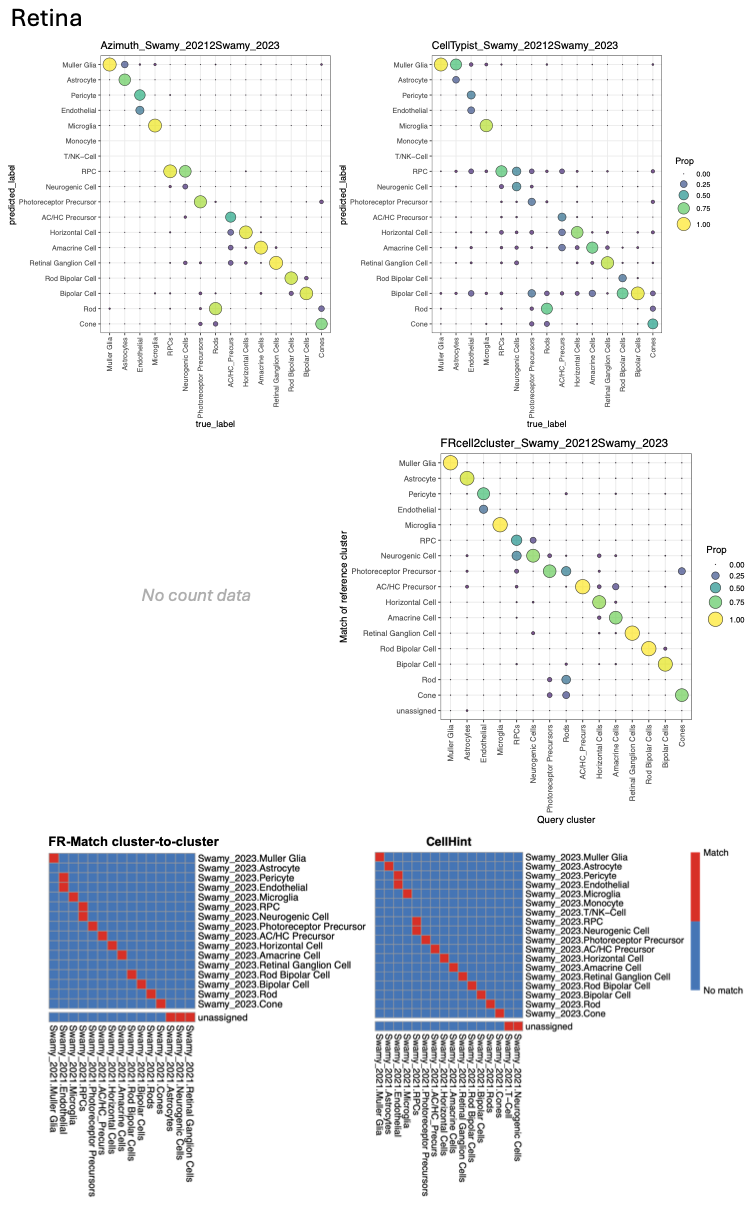

**Supplementary Figure 12. Matching results of retina cell types.**

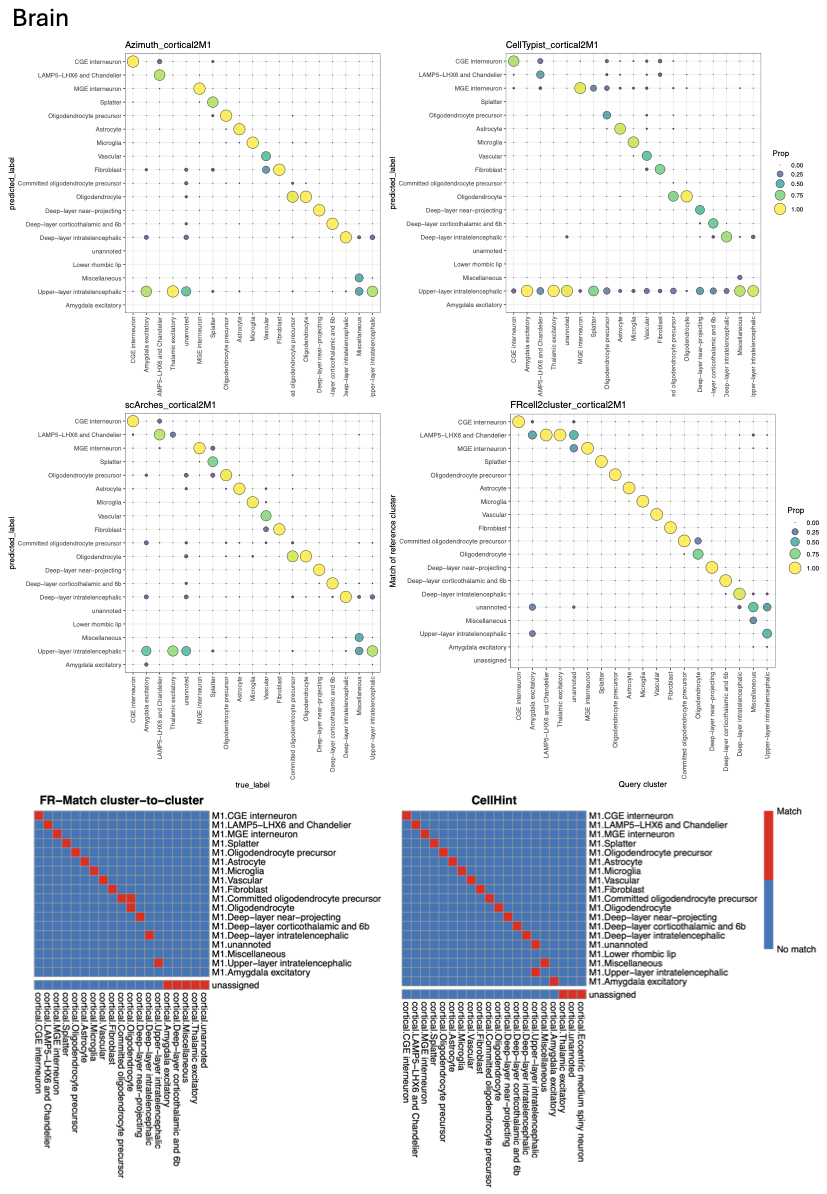
**Supplementary Figure 13. Matching results of brain cell types.**

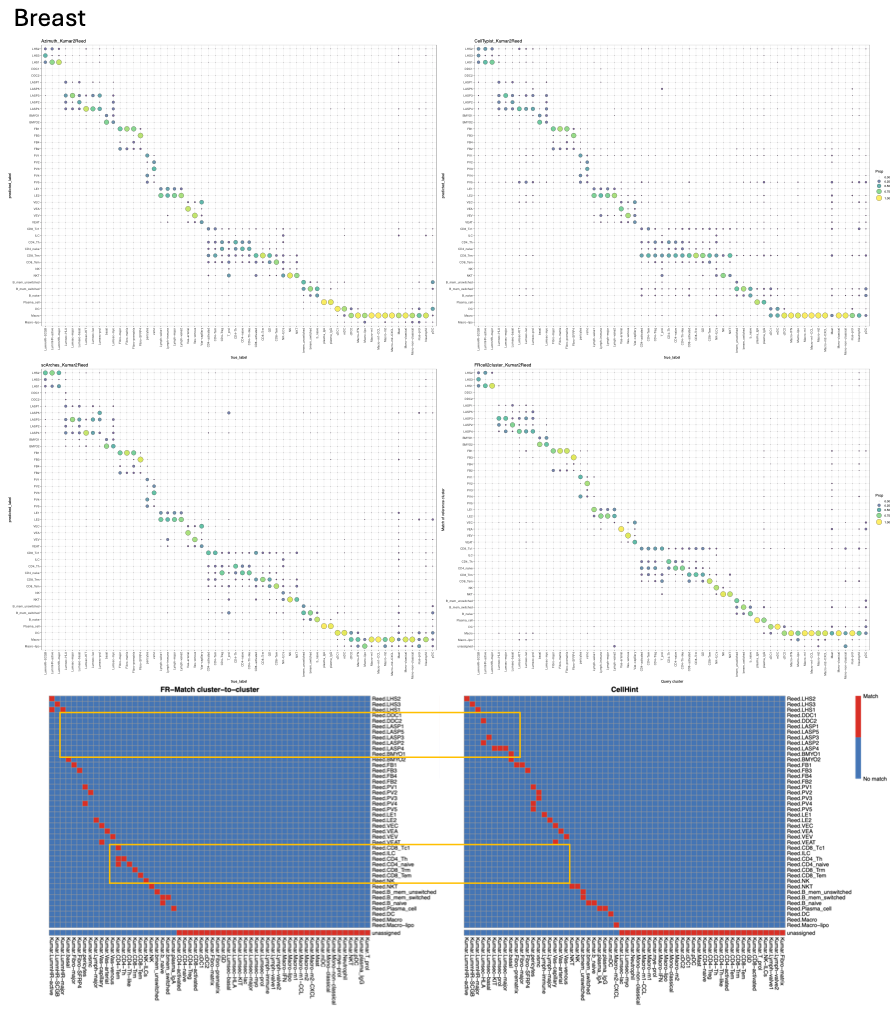
**Supplementary Figure 14. Matching results of breast cell types.** Highlighted boxes show the complementing matching between FR-Match and CellHint.

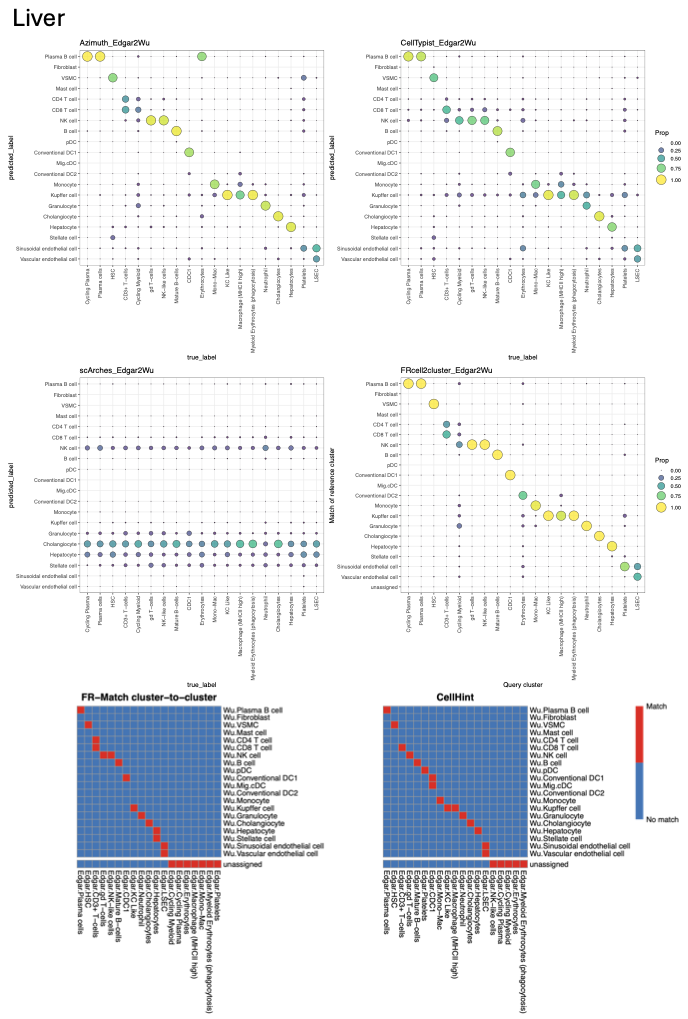

**Supplementary Figure 15. Matching results of liver cell types.**

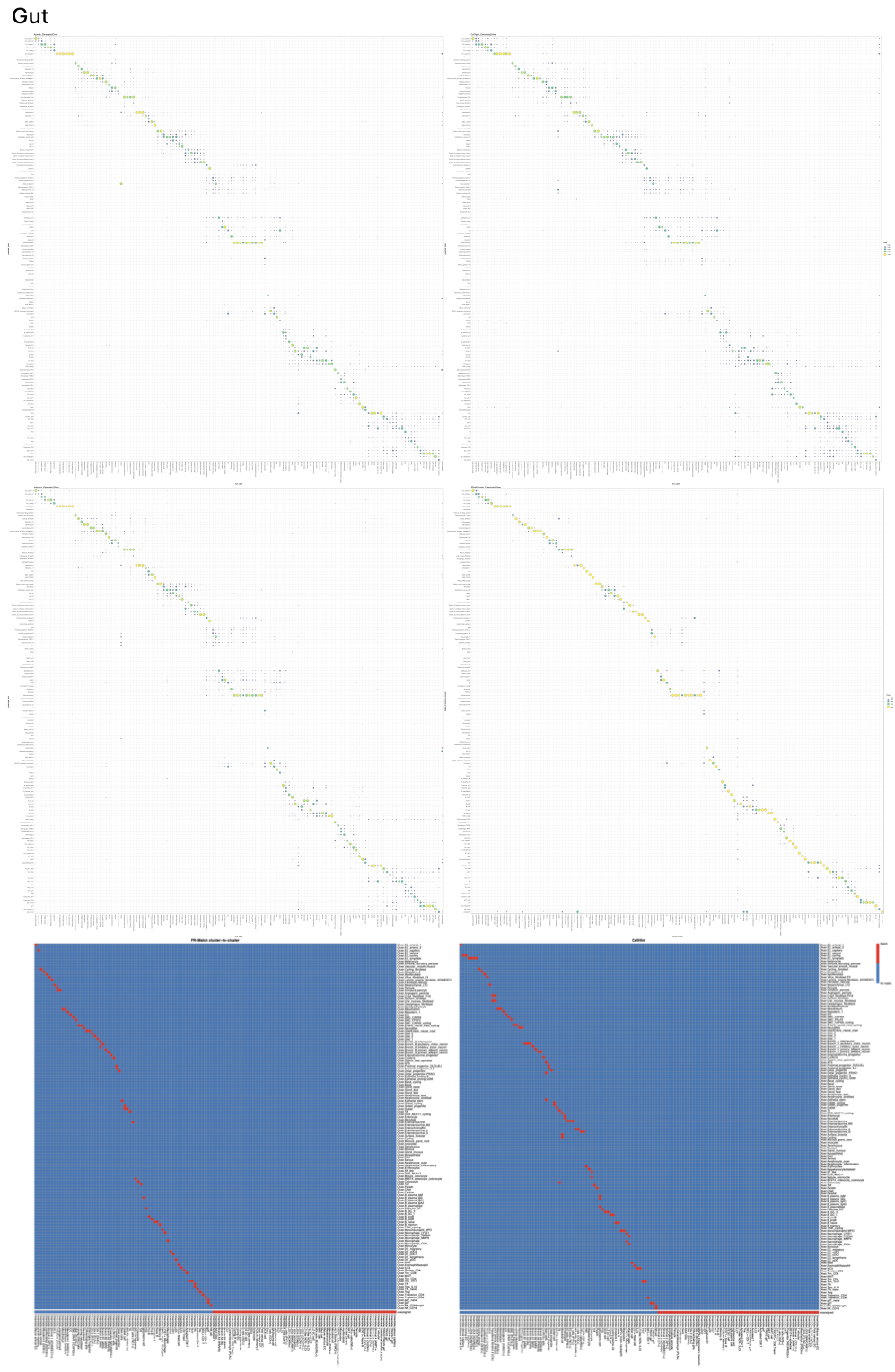
**Supplementary Figure 16. Matching results of gut cell types.**

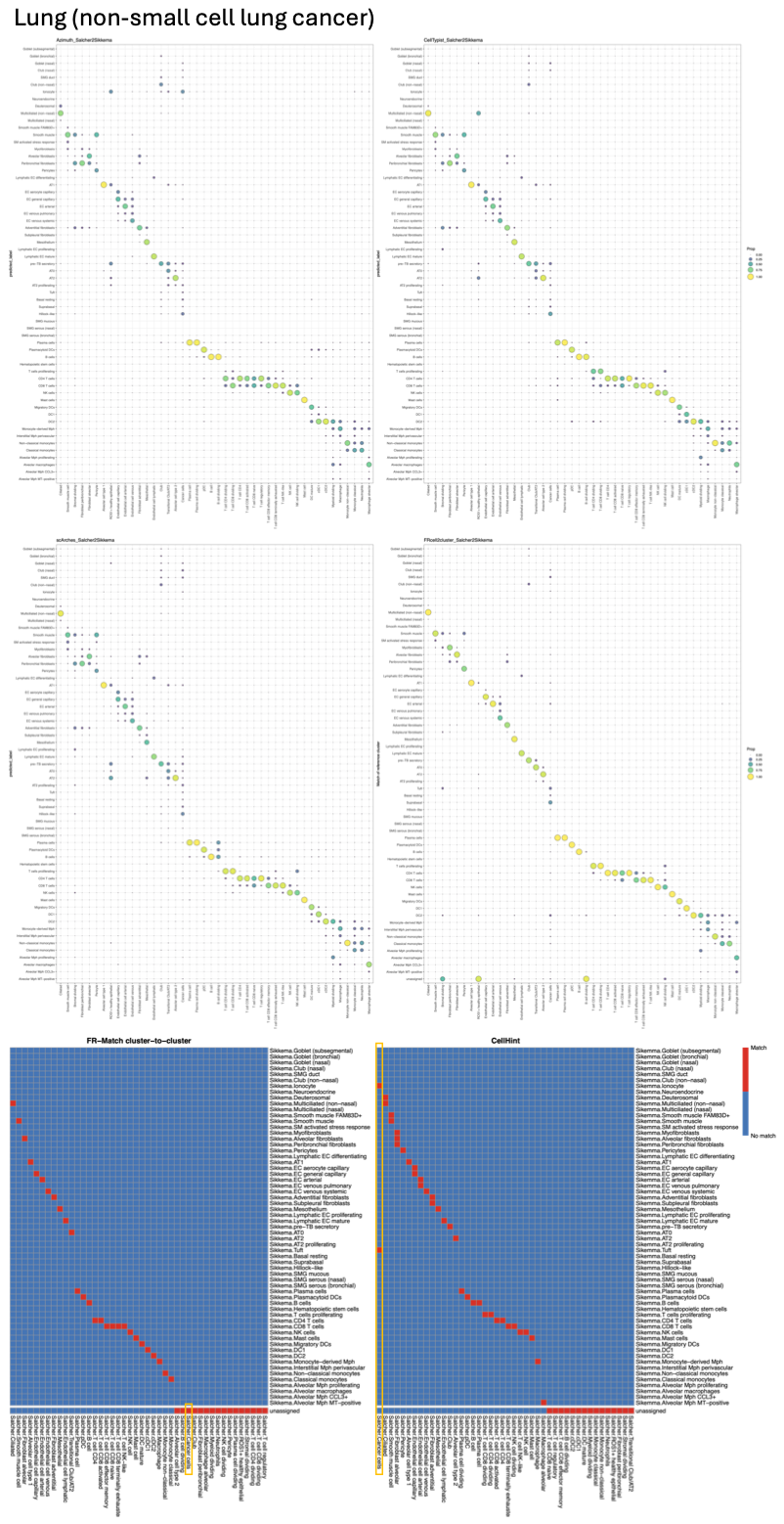

**Supplementary Figure 17. Matching results of small cell lung cancer cell types.** Highlighted boxes indicate the cancer cells.

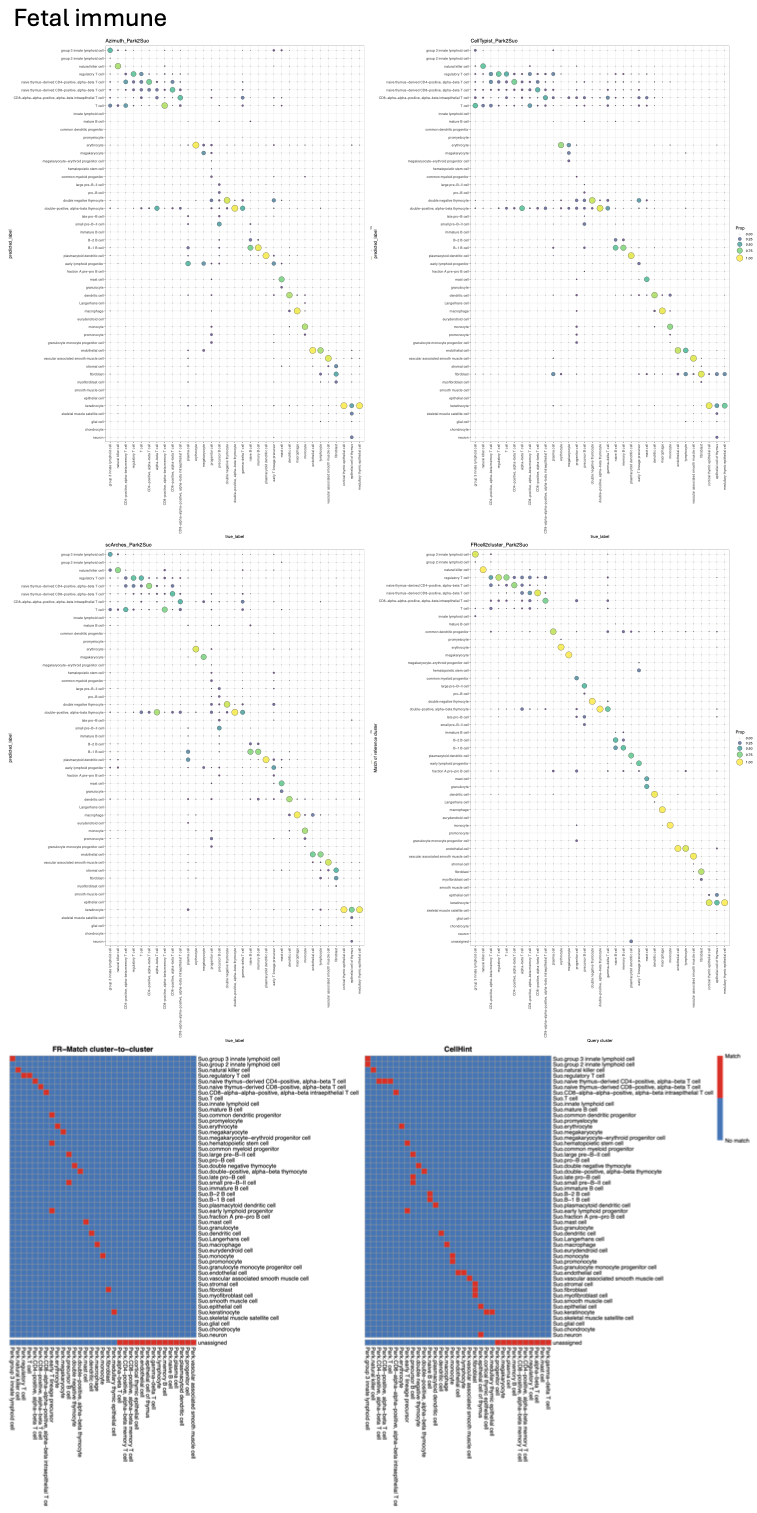

**Supplementary Figure 18. Matching results of fetal immune cell types.**

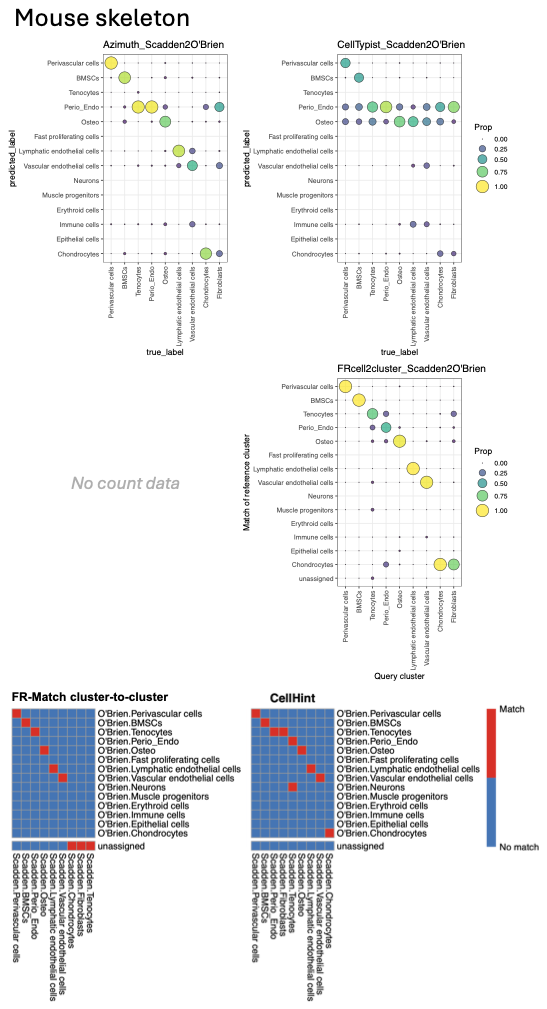

**Supplementary Figure 19. Matching results of mouse skeleton cell types.**

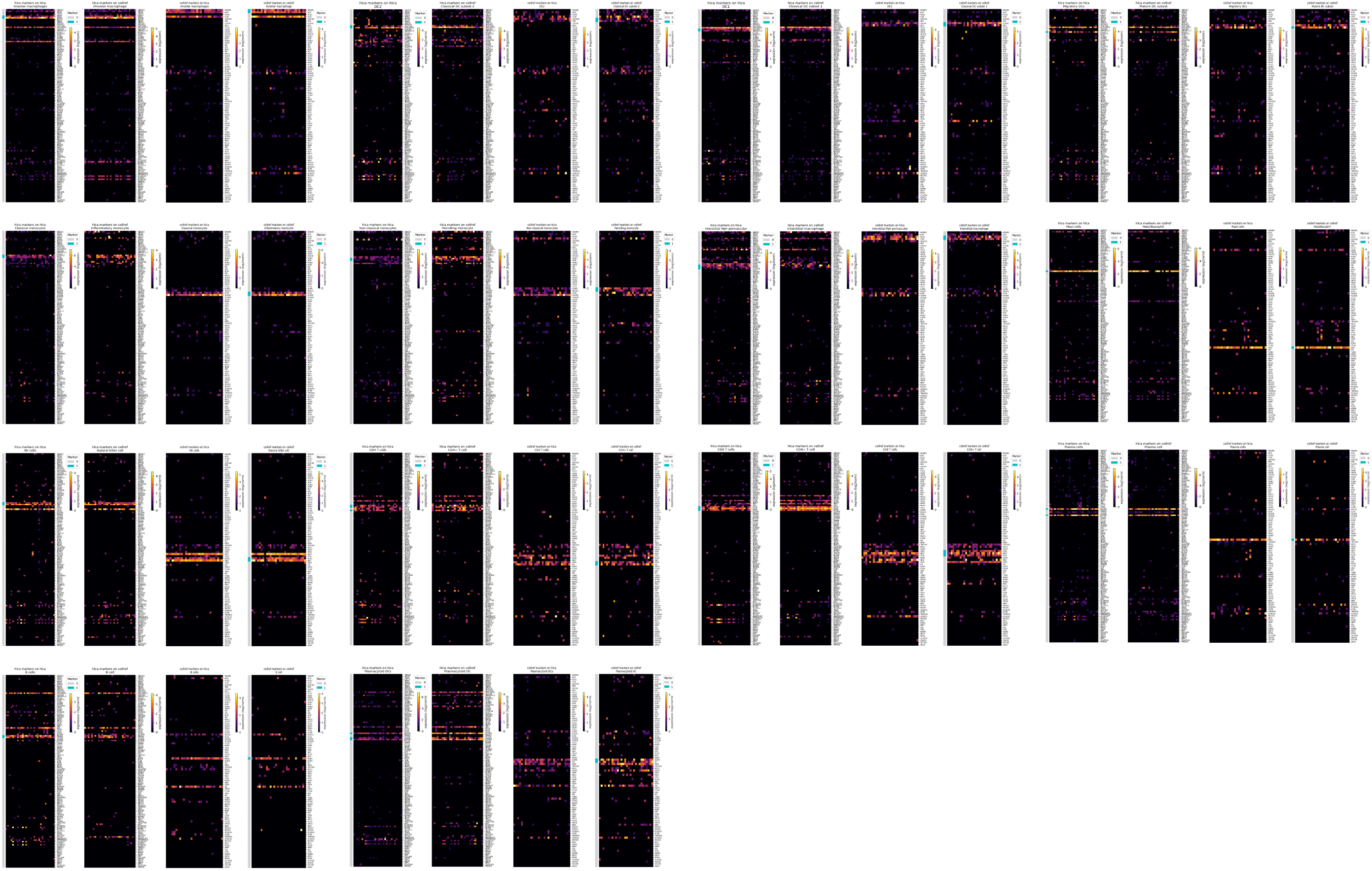

**Supplementary Figure 20. Cell type barcode plots for the recommended matches of cell types in the immune branch of the HLCA dendrogram.**

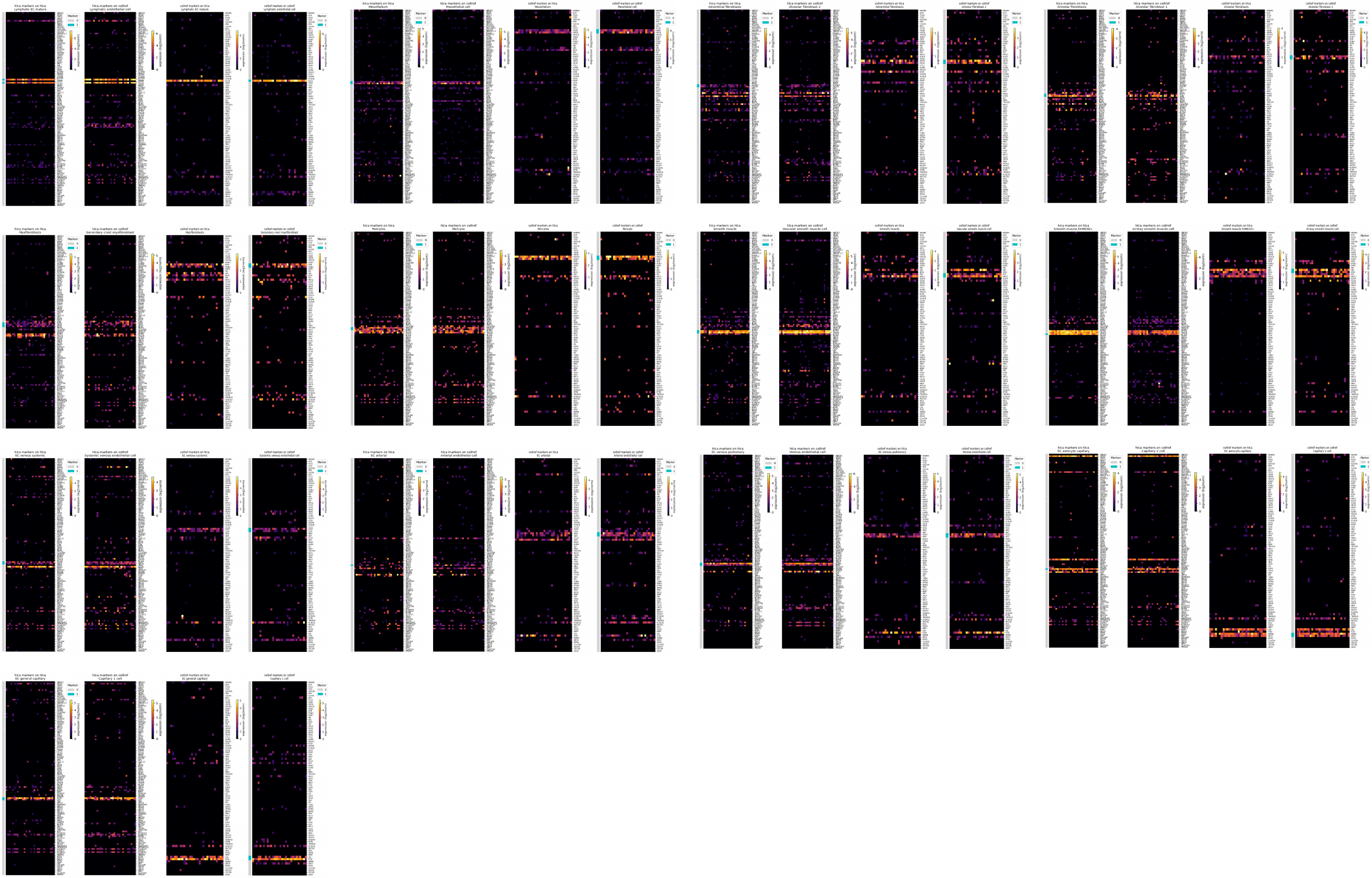

**Supplementary Figure 21. Cell type barcode plots for the recommended matches of cell types in the endothelial and stromal branch of the HLCA dendrogram.**

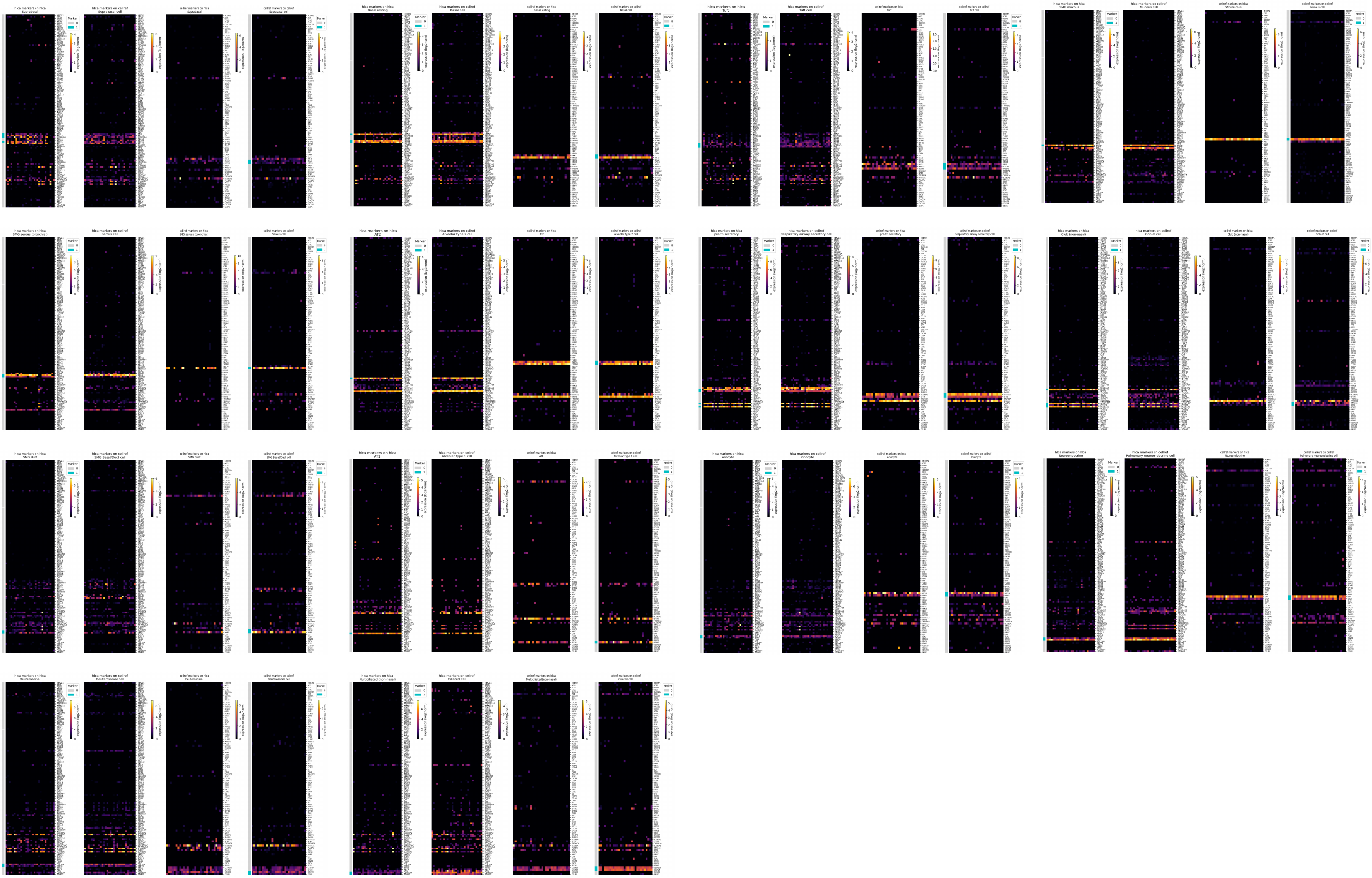

**Supplementary Figure 22. NS-Forest barcode plots for the recommended matches of cell types in the epithelial branch of the HLCA dendrogram.**

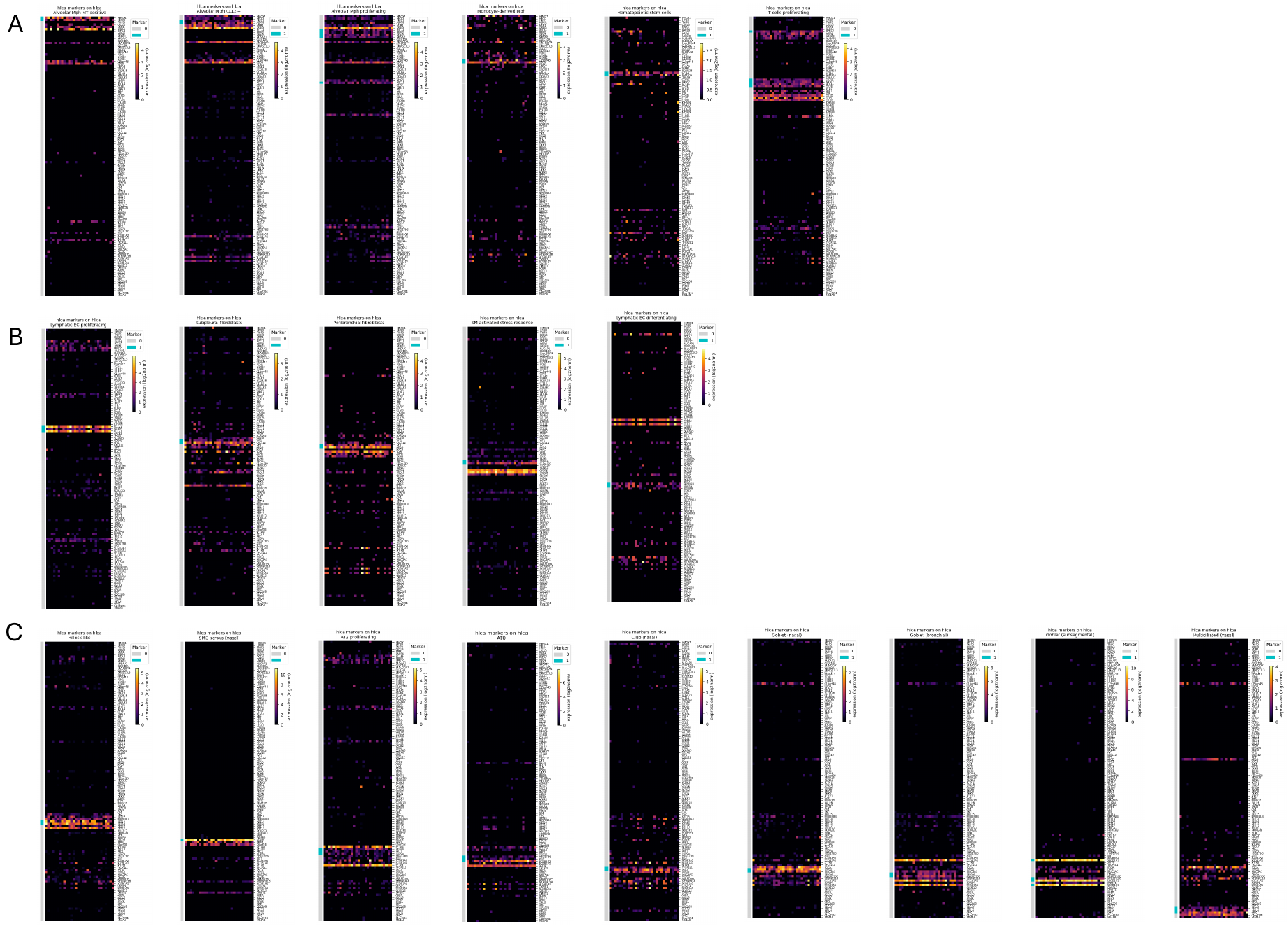

**Supplementary Figure 23. NS-Forest barcode plots for the 20 HLCA-specific cell types. (A)** Immune cell types in dendrogram order. **(B)** Endothelial and stromal cell types in dendrogram order. **(C)** Epithelial cell types in dendrogram order.

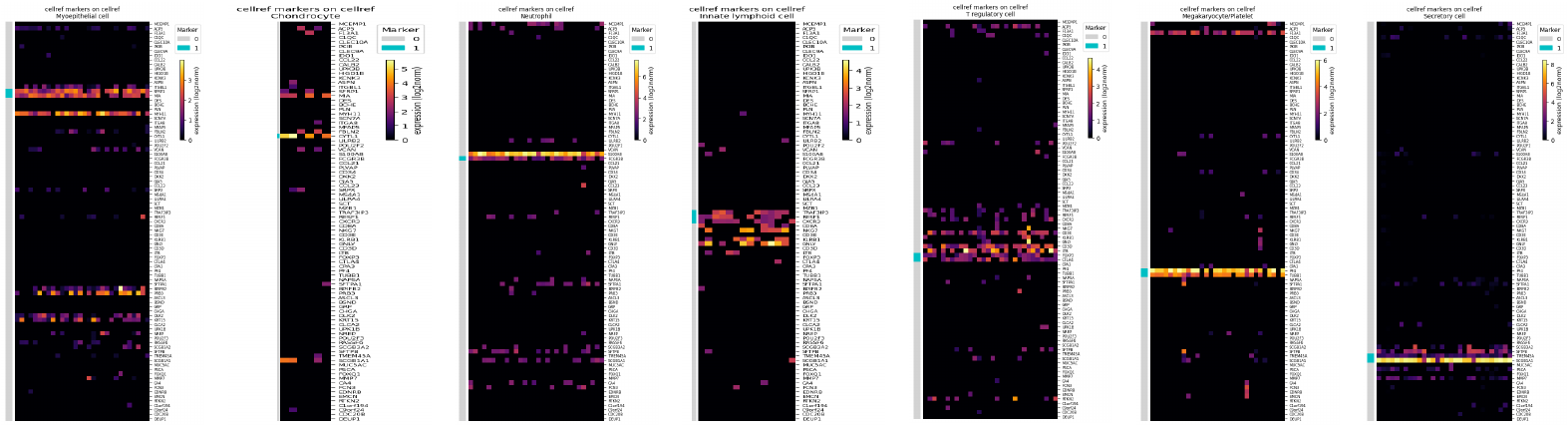

**Supplementary Figure 24. NS-Forest barcode plots for the 7 CellRef-specific cell types.**

**Supplementary Table 1. Cell type predictions and scores for matching the CellRef cells to the HLCA reference.**

**Supplementary Table 2. The healthy human lung cell type meta-atlas across CellRef and HLCA and characterizing marker genes for each cell type.**

**Supplementary Table 3. High confidence match values for the recommended matches.**

**Supplementary Table 4. All high confidence matches identified by Azimuth.**

**Supplementary Table 5. All high confidence matches identified by CellTypist.**

**Supplementary Table 6. All high confidence matches identified by scArches.**

**Supplementary Table 7. All high confidence matches identified by FR-Match.**

**Supplementary Table 8. Dataset and result summary of more studies.**
